# Bias in the arrival of variation can dominate over natural selection in Richard Dawkins’ biomorphs

**DOI:** 10.1101/2023.05.24.542053

**Authors:** Nora S. Martin, Chico Q. Camargo, Ard A. Louis

## Abstract

Biomorphs, Richard Dawkins’ iconic model of morphological evolution, are traditionally used to demonstrate the power of natural selection to generate biological order from random mutations. Here we show that biomorphs can also be used to illustrate how developmental bias shapes adaptive evolutionary outcomes. In particular, we find that biomorphs exhibit phenotype bias, a type of developmental bias where certain phenotypes can be many orders of magnitude more likely than others to appear through random mutations. Moreover, this bias exhibits a strong Occam’s-razor-like preference for simpler phenotypes with low descriptional complexity. Such bias towards simplicity is formalised by an information-theoretic principle that can be intuitively understood from a picture of evolution randomly searching in the space of algorithms. By using population genetics simulations, we demonstrate how moderately adaptive phenotypic variation that appears more frequently upon random mutations will fix at the expense of more highly adaptive biomorph phenotypes that are less frequent. This result, as well as many other patterns found in the structure of variation for the biomorphs, such as high mutational robustness and a positive correlation between phenotype evolvability and robustness, closely resemble findings in molecular genotype-phenotype maps. Many of these patterns can be explained with an analytic model based on constrained and unconstrained sections of the genome. We postulate that the phenotype bias towards simplicity and other patterns biomorphs share with molecular genotype-phenotype maps may hold more widely for developmental systems, which would have implications for longstanding debates about internal versus external causes in evolution.

## INTRODUCTION

### Three versions of the infinite monkey theorem

In his influential book, *The Blind Watchmaker* [1], Richard Dawkins illustrates how natural selection can efficiently find fitness maxima in ‘hyper-astronomically large’ [2] search spaces by introducing an intriguing twist on the famous infinite monkey theorem. He frames his argument by first introducing the classic case (See Fig 1) with a question: How likely is it that a monkey randomly typing on a typewriter produces Hamlet’s 28-character phrase “METHINKS IT IS LIKE A WEASEL”? For a monkey typing on an *M* -key typewriter, the probability to produce a specific string of *n* characters will scale as 1*/M^n^*, which rapidly becomes unimaginably small with increasing *n*. By analogy, random mutations on their own are unlikely to produce meaningful biological novelty. Dawkins contrasts this picture with his second version of the infinite monkey theorem, where the novel twist is to include a fitness function that acts on each letter independently. The output stops changing once the correct letter is found, so that on average only *M* random keystrokes are needed for each letter. Thus, any *n* letter phrase can be produced in a number of keystrokes that scales as *n × M*, which is exponentially smaller than in the classical case. This simple but evocative example illustrates an important property of biological sequence spaces. For a given alphabet size *K*, their size grows exponentially with sequence length *L* as *K^L^*, but genomic distances remain linear in *L* because on the order of *L* mutations can be used to link any two sequences. By using fitness functions of the kind that Dawkins introduced, an evolutionary search algorithm can exploit this linearity and locate a fitness maximum in an exponentially large high-dimensional search space within a relatively small number of randomly generated steps.

**Figure 1.**
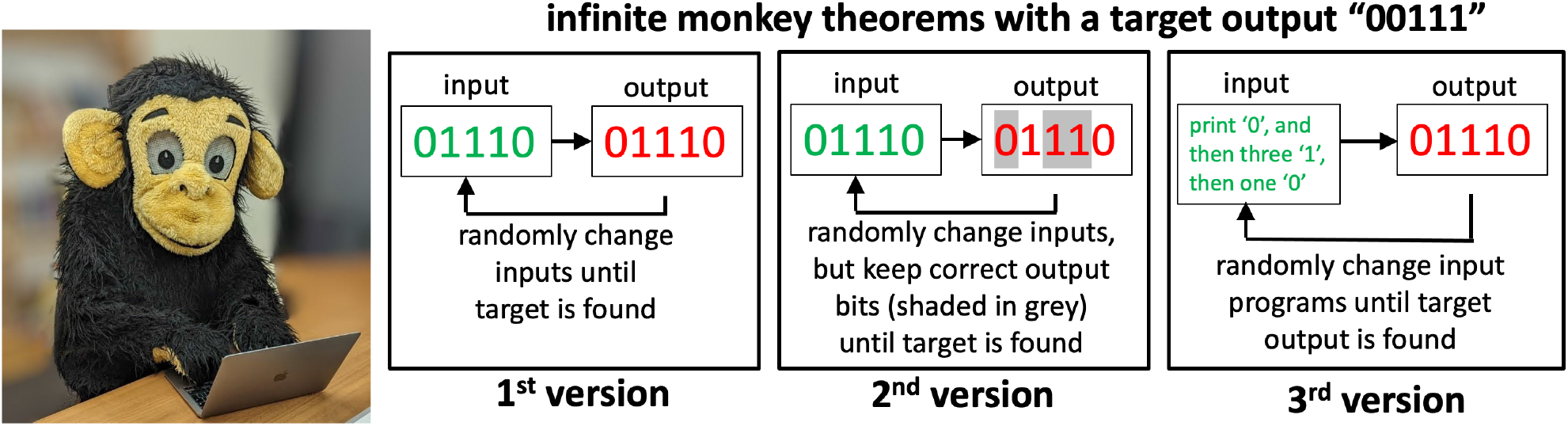
Three versions of the “infinite monkey theorem” compared. In the 1^st^ version, or the classic case, all outputs of length *n* are equally likely, and thus the probability of obtaining a specific output scales as 1*/M ^n^*, where *M* is the number of keys, and *n* is the length of the desired output. In the 2^nd^ version, introduced by Dawkins [1], a fitness function fixes each correct partial output, so that a desired *n*-length string is likely to be found in a timescale that scales as *n × M*, which is linear instead of exponential in *n*. In the 3^rd^ algorithmic version, the probability of obtaining an output scales as 1*/M ^K^*, where *K* is the length of a program that generates it [3]. Outputs that can be generated by short programs are therefore exponentially more likely to obtain. The length of the shortest program that generates an output is related to the famous Kolmogorov complexity measure [4] so that the algorithmic monkey theorem also implies a bias towards simplicity.

In this paper, we explore the evolutionary consequences of a third (algorithmic) version of this famous trope of monkeys on keyboards (see Fig 1). In Dawkins’ version, the monkeys directly type out components of the outputs, i.e. the phenotypes. In evolution, however, novel phenotypic variation is generated indirectly by random mutations which are then “decoded” through the process of development. To capture this mapping from genotypes (the inputs) to phenotypes (the outputs), consider instead monkeys generating outputs by typing at random into a computer programming language [3]. In contrast to the classical version of the infinite monkey theorem, where all output strings of length *n* are equally likely (with probability *p* = 1*/M^n^*), in the algorithmic picture, certain outputs are exponentially more likely to appear than others. For example, a string of length *n* = 1000 of the form “010101…” would appear when typing the 21-character program “print “01" 500 times;” [3]. Therefore, its probability *p* = 1*/M* ^21^ is many orders of magnitude larger than the probability *p* = 1*/M* ^1000^ for the classical version. Thus, within this algorithmic picture, there are certain kinds of outputs, namely those that can be described by short programs, which have an exponentially higher probability than outputs without such short algorithmic descriptions. Interestingly, an algorithmic picture of evolution is also introduced in a famous passage from chapter 5 of *the Blind Watchmaker* [1], where Dawkins describes seeds falling from a tree: *“It is raining instructions out there; it’s raining programs; it’s raining tree-growing, fluff-spreading, algorithms. That is not a metaphor, it is the plain truth. It couldn’t be any plainer if it were raining floppy discs.”*.

Can the intuitive link between our simple algorithmic picture and the mapping from genotypes to phenotypes be made more rigorous? To this end, we turn to the field of algorithmic information theory (AIT) [4] where the central concept is the Kolmogorov complexity *K*(*x*) of a string *x*. *K*(*x*) is the length of the shortest program that generates *x* on a universal Turing machine, a computing device that can perform any possible computation. There are profound mathematical relationships in AIT that link probability and Kolmogorov complexity [4]. Here we apply a recently-derived AIT theorem [5] which predicts an upper bound for the probability *P* (*x*) that an output *x* is obtained upon random sampling of inputs of a computable input-output map:

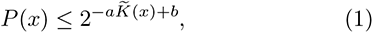

where the descriptional complexity *K*(*x*) is a suitable approximation to the (uncomputable) Kolmogorov complexity, and two constants *a* and *b* are independent of the outputs *x*. Interpreting *K̃* (*x*) as a measure of the length of the shortest program that generates output *x* connects directly to the algorithmic picture of monkeys typing into a computer programming language. Probabilities scale exponentially with program length so that the shortest program has the highest probability. This kind of bias towards short descriptions is called “simplicity bias” in the context of computable input-output maps [5]: Outputs with high *P* (*x*) will have small *K̃* (*x*), and outputs with large *K̃* (*x*) will have low *P* (*x*) (but not necessarily vice-versa because Eq (1) is an upper bound). In [3, 5, 6] it was shown that the bound (1) works remarkably well for a wide range of input-output maps, suggesting that this information-theoretic link between probability and complexity holds widely.

### Simplicity bias in genotype-phenotype maps

It has recently been argued [3] that many genotype-to-phenotype (GP) maps obey the mathematical conditions needed for Eq (1) to be satisfied, formalizing the intuitive connection between GP maps and the 3^rd^ algorithmic monkey theorem. GP maps typically exhibit redundancy due to neutral mutations [7], where ’neutral’ simply means that the mutation does not change the phenotype, which is a simpler definition than the classical notion introduced by Kimura [8]. This redundancy naturally leads to the concept of a *neutral set* made up of all the genotypes that map to a given phenotype *p*. We can define the associated probability *P* (*p*) that a randomly selected genotype belongs to the neutral set of *p*, which is also referred to as the phenotype frequency *f_p_* of *p*. It is directly proportional to the size of the neutral set. *Phenotype bias* occurs when there are large differences in the *neutral set sizes* (or equivalently in the *f_p_*) associated with different phenotypes *p* [9]. The AIT simplicity bias bound (1) above predicts that GP maps will exhibit an Occam’s razor-like phenotype bias where the largest frequencies correspond to low-complexity phenotypes^1^. These simple phenotypes are therefore more likely to appear upon random mutations than individual complex phenotypes are.

Strong evidence for this hypothesis of a “biological Occam’s razor” was found at the molecular scale for the GP maps of RNA secondary structure, the polyomino model for protein quaternary structure, and a popular model of the yeast cell-cycle gene regulatory network [3]. For example, phenotype bias towards simplicity can explain key patterns in nature such as an observed strong preference for symmetry in protein complexes, and the fact that the most frequent RNA secondary structures found in nature have structures that are highly compressible, and therefore are simple with low descriptional complexity *K̃* (*p*) [3].

To illustrate in more detail how phenotype bias can affect patterns observed in nature, we discuss the best-understood molecular GP map, namely that of the secondary structures of RNA. The advantage of RNA secondary structure is that an abundance of data is available in standardized databases of functional RNA, which include catalysts and structural elements in the cell. The total number of possible sequences grows extremely rapidly with length. For example, the set of all 4^126^ possible RNA sequences of length *L* = 126, which is not that long for functional RNA, would have more mass than the observable universe [2]! Given the size of these possibility spaces, one would not expect to observe the same structure twice when randomly searching such a hyper-astronomically large genetic search space. But this expectation is mistaken: strong phenotype bias means that some structures are found with high frequency even in a relatively small random sequence sample [9, 11–14]. For example, if the secondary structures are coarse-grained using level-5 of the RNAshapes method [15], then the 68 evolved secondary structures of length *L* = 126 found in the RNAcentral database [16] of functional RNA are among the 96 structures with highest phenotypic frequencies out of a much larger set of 10^12^ topologically possible level 5 structures [9]. Interestingly these 68 observed secondary structures are typically found by sampling less than 10^6^ random sequences, an unimaginably small 10*^−^*^70^th fraction of the total number of distinct possible sequences of that length^2^. The mechanisms by which strong phenotype bias is predicted to influence adaptive evolutionary outcomes includes the “arrival-of-the-frequent” effect [17], which captures the simple fact that natural selection can only act on the structures that are introduced sufficiently frequently into the population through random mutations, see also [18]. Depending on the relevant time scales and mutation rates, concepts such as “free-fitness” [19, 20], or the “survival of the flattest” [21] are similarly predicted to favor the evolution of high-frequency structures. To sum up, for RNA, the evolved structures dramatically reflect the strong phenotype bias in this system. This observation does not negate the role of selection. Each functional RNA structure in the database will have fixed due to natural selection, and a randomly selected sequence would be unlikely to perform a given biological function (see [22] for a recent discussion). But it does mean that nature was able to produce the “*endless forms most beautiful* ” [23] that grace the living world from only a minuscule fraction of the set of all RNA structures, namely those that are most likely to appear as variation.

While molecular GP maps such as the RNA model above can be interpreted as a stripped-down version of developmental bias [24, 25], historically much of the interest in the effects of bias on the arrival of variation has focused on morphological evolution. Could simplicity bias also have a dramatic impact on this larger scale? A recent study of an abstract morphological model of tissues found that random developmental mechanisms are more likely to be associated with simple morphologies and moreover, that complex morphologies are less robust to parameter changes [26]. Similarly, in a model of digital organisms [27], it was found that simple phenotypes are generated by a higher number of genotypes and are more likely to evolve from another phenotype. Higher phenotypic frequencies for simpler phenotypes were also found in a model of digital logic gates [28, 29], Boolean threshold models for gene regulatory networks [30] and a highly simplified model of neural development [31]. As a further example, models based on Lindenmeyer systems, a recursive model that can generate plant-like shapes [32] or sequences of symbols, indicate that simple phenotypes are more robust to mutations [10] and have higher neutral set sizes [5].

In order to address the status of phenotype bias in systems beyond the molecular scale, we will focus on another important innovation from *The Blind Watchmaker* [1], a developmental model of two-dimensional shapes called *biomorphs*. As illustrated in Fig 2, these are made up of vectors, which are defined by the (numeric) genotypes and combined into a biomorph phenotype in a recursive developmental process. This model produces a rich array of forms. In his book [1], Dawkins was able to gradually steer the evolution of biomorphs towards particular desired shapes in a relatively small number of generations by carefully choosing phenotypes that appear upon random mutations. In this way, he used biomorphs to illustrate the power of natural selection in a more complicated system than the simple “WEASEL” program. The main aim of this paper will be to analyze the generation of phenotypic variation more systematically in this system and test the hypothesis that this iconic model of morphological development also exhibits simplicity bias and other phenomena similar to those observed for molecular GP maps, and furthermore, to analyze the effect that these biases in the arrival of variation have on evolutionary dynamics.

**Figure 2.**
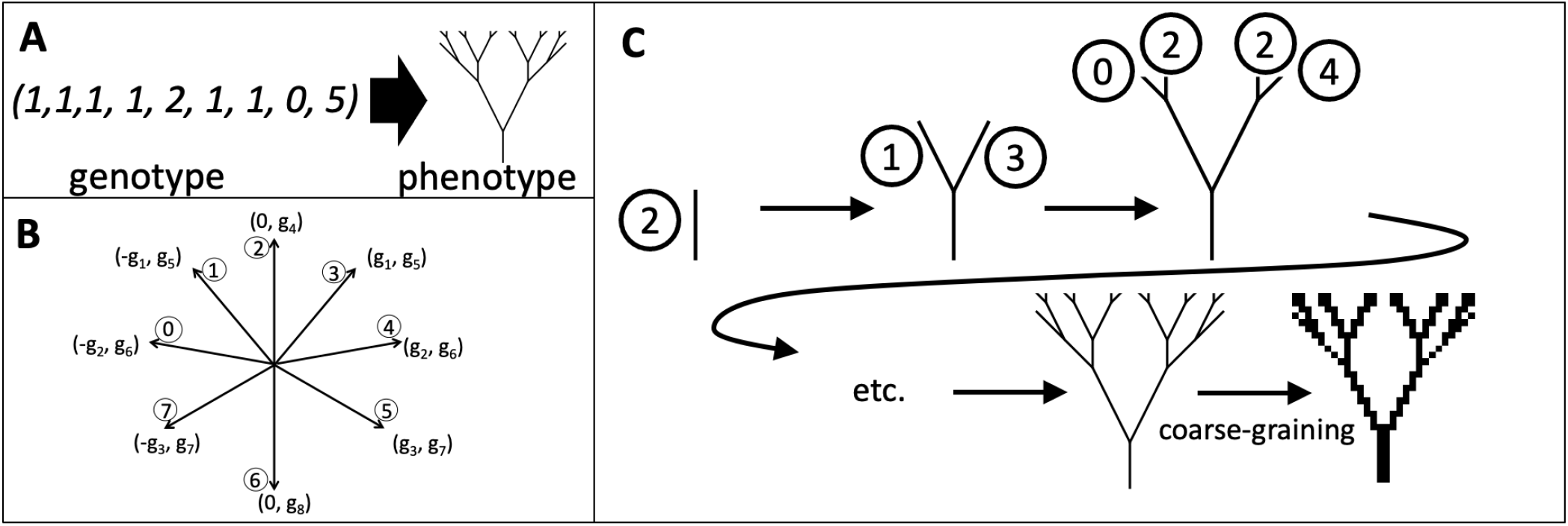
The biomorphs GP map: A) Following [1, 33] each genotype, a set of nine integers, is linked to the corresponding biomorph phenotype, a 2D shape. B) To obtain the phenotype from a given genotype, the integers at the first eight positions of the genotype (labeled *g*1 - *g*8) are used to define eight two-dimensional vectors, numbered from 0 to 7, as shown schematically above. C) The 2D biomorph shape is created from these vectors by applying a set of simple rules for several developmental stages, the number of which is set by the integer at the ninth position of the genotype (see Methods). In order to discretize the phenotypes for our computational GP map analysis, we coarse-grain the final image on a 30 *×* 30 grid, as shown in the final step.

We analyze the biomorphs GP map as follows. Firstly, to take into account the fact that many biomorph phenotypes are nearly indistinguishable to the eye, we define a coarse-graining that maps them onto a discrete 30 *×* 30 pixel grid, as shown in Fig 2 C. We then exhaustively analyse all genotypes within a fixed parameter range, and use an approximate descriptional complexity measure [34] to show that the frequency-complexity relationship of biomorph phenotypes is indeed consistent with the simplicity bias of Eq (1). We show that the GP map of biomorphs exhibits many other properties that resemble those commonly found in molecular GP maps, as reviewed in [7, 35]. For example, the phenotype robustness *ρ_p_*, defined as the mean mutational robustness of all genotypes *g* that map to a given phenotype *p*, scales as the logarithm of the frequency *f_p_*of the phenotype. Evolvability, a measure that counts how many novel phenotypes are accessible by point mutations, correlates negatively with the mutational robustness *ρ_g_* of an individual genotype *g*, but positively with phenotype robustness *ρ_p_* of the whole neutral set [36]. We can rationalize these effects in both existing GP maps and in the biomorphs system through a simple analytically tractable model based on separating genotypes into constrained and unconstrained portions [37–40].

Another big question is to what extent these structural GP map characteristics, which determine the spectrum of novel variation that appears upon random mutations, affect evolutionary outcomes, especially when natural selection is also at play. We first show that in the absence of selection, biases in phenotypic frequencies (which are calculated on a uniform random sampling of genotypes) are reflected in the average rates with which each biomorph phenotype appears in an evolving population. Next, we turn to a scenario that is adapted from refs [17, 18] and includes both variation and selection: Two adaptive phenotypic changes are possible and for a range of fitness values, we find that the more frequent phenotype fixes first even though it is not the fittest phenotype. We also study a scenario from Dawkins’s book [1] where he finds it hard to reconstruct an evolutionary pathway to an ‘insect’-shaped phenotype. He argues that for such rare phenotypes, while short paths exist, these are only a tiny fraction of a much larger set of potential paths, and so they are hard to reliably find. We illustrate these shortest paths and note that if neutral mutations are included, fewer phenotypic changes are needed, making it easier to create fitness functions that lead to monotonically increasing fitness paths to the final desired phenotype.

Finally, we situate our specific findings for the biomorph system within the longstanding debate about the relative contributions of internal (or structuralist) causes versus external (or adaptationist) causes in determining evolutionary outcomes. In particular, we explore whether phenotype bias could join natural selection as an ultimate, rather than a proximate evolutionary cause in Mayr’s [41] influential classification. To clarify and unravel some of the complex strands that have historically had an impact on his broader argument, we first review in some detail how the better-understood example of RNA secondary structure for functional RNA fits into this dichotomy. Given that that biomorphs exhibit similar phenotype bias, we then explore how the hypothesis that such bias holds more widely for developmental systems would impact on the contentious question of where developmental constraints and biases fit in within the causal structure of evolutionary biology [42–56].

## MATERIALS AND METHODS

### Dawkins’ biomorphs model

In Dawkins’ biomorphs model [1, 33], phenotypes are two-dimensional figures, recursively constructed from genotypes, which consist of nine genes *g*_1_-*g*_9_, represented by integer values, (Fig 2A). This construction is performed in two steps: first, a set of eight vectors is constructed from the genotypic information and then these vectors are combined recursively to form the final figure, as described in [33]:

1. **Definition of eight two-dimensional vectors:** The x- and y-coordinates of eight two-dimensional vectors are set by the values of the first eight genes, *g*_1_-*g*_8_, as shown in Fig 2B: for example, the vector number 3 has an x-component set by the integer at the first gene (*g*_1_) and a y-component set by the integer at the fifth gene (*g*_5_). The allocation of specific genes to vector components is fixed by Dawkins’ definition of the biomorphs system, as described in [33].
2. **Recursive construction of the final biomorph from eight two-dimensional vectors:** These eight vectors, denoted vector 1 to vector 8, form the basis of a recursive developmental process, where vectors are added to the figures in several stages, as sketched in Fig 2C. The ninth gene determines after how many stages this process terminates. At the first stage, a single vector is drawn (by convention, this is ‘vector 2’ with its length multiplied by the value of *g*_9_). At the next stage, vector (2 + 1) and vector (2 *−* 1) are multiplied by *g*_9_ *−* 1 and both are attached to the endpoint of the existing vector. At each further recursion, vectors *n −* 1 and *n* + 1, where *n* is the number of the preceding vector, have their length multiplied by a successively smaller integer and attached to the current endpoint of the figure, as demonstrated in Fig 2C. The numbering of the eight vectors is treated periodically, i.e. the next-highest vector from vector 7, the final vector, is vector 0 and conversely, the vector corresponding to (0-1) is vector 7.

In this way, the biomorph system corresponds to a scenario where the key developmental mechanisms are fixed: two new lines grow out of each endpoint of the biomorph shape at each developmental stage. However, the number of stages and the details of individual entities, i.e. the lengths and angles of the vectors, vary from genotype to genotype (a vector could even have zero length).

In order to exhaustively analyze the GP map computationally, we restrict the values in the genotypes to a finite range. We take 7 values for each of the ‘vector genes’ (*−*3 *≤ g_i_ ≤* 3 for *i ∈* [1*, ..,* 8]) and 8 values for the ninth gene (1 *≤ g*_9_ *≤* 8). In this range, there are 7^8^ *∗* 8^1^ = 46, 118, 408 genotypes. This range is somewhat smaller than the values in Dawkins’ examples [1], but they are near the limit of what is feasible for exhaustive enumerations. We chose a slightly higher range for the ninth gene than for the first eight genes since changes in the ninth gene affect the number of drawn lines and therefore have the greatest qualitative effect. The effect of extending these ranges further can be investigated with the approximate analytic model introduced in this paper. We find that the qualitative observations are unchanged (section S3.3 in the *Supplementary Information*).

Following Dawkins’ program of artificial evolution [1], a point mutation can increase or decrease a single gene by one integer step. This is a key difference from models like RNA, where each nucleotide can be exchanged for any other nucleotide. We do not include mutations that go beyond our fixed range of genotypes.

### Quantifying the biomorphs GP map

We use two different approaches to study the relationship between biomorph genotypes and phenotypes on a large scale. The first approach is computational: we simply consider all genotypes within a fixed range and generate their phenotypes computationally. In order to be able to manipulate, analyze and compare the phenotypes computationally, we coarse-grain them on a 2D grid, as explained below. The second approach is an analytic model based on separating the genome into constrained and unconstrained parts, a simplification which makes it possible to (semi)analytically calculate some key properties of a GP map [37, 38, 40].

Throughout the paper, where we do not specify, the results are based on the computational approach. For example, our computational simulations of evolving populations use only the computational GP map. However, many steps of the GP map analysis are performed for both the computational and the analytic approaches. Good agreement is found between the two approaches.

#### Computational model with discrete phenotypes

For our computational analysis, we need a clear definition of when two biomorphs share the same phenotype. This definition should mimic the conditions in the original evolution experiments by Dawkins [1], who applied artificial selection based on the entire appearance of a biomorph (rather than just a specific feature). Moreover, the biomorphs were drawn on a computer screen of limited size, such that very small features may have appeared indistinguishable. Thus, biomorphs should only be treated as distinct phenotypes if they display clear visual differences. To reproduce this delineation, we project the 2D shape onto a limited-resolution 30 *×* 30 pixel grid as illustrated in the final step of Fig 2C. In detail, this procedure works as follows:

- First, we go through the lines and merge any coinciding line segments (i.e. if the identical line segment is drawn as part of two longer lines, only one instance is kept). We only work with one half of the biomorph since the other half is given by axial symmetry.
- Secondly, we place the lines on the grid - the lines are scaled such that the total size of the grid is 5% larger than the longer dimension of the biomorph shape (either width or height) and the biomorph is placed at the center of the grid.
- Next, we record, how many lines are contained within each pixel on the grid as follows: we simply compute the total length of all line elements within the pixel (for computational reasons, we round to the nearest 10*^−^*^3^ in our calculations). Lines coinciding with the outer boundary of a pixel are assumed to contribute half their length to the pixels on either side of the boundary.
- Finally, we go through each pixel: if the total line length contained within the pixel is *≥* 20% of the side length of the pixel, the pixel value is set to one. Otherwise, it is set to zero.

This coarse-graining method has two parameters: the grid resolution (30 *×* 30) and the threshold for setting a pixel to one (*≥* 20% of the length of the side of the pixel). In the *Supplementary Information* section S3, we show that the qualitative characteristics of the GP map are robust to changes in these two parameters.

#### Analytic model based on sequence constraints

By separating a genotype into constrained and unconstrained positions, it has been possible to (semi-)analytically calculate many properties of GP maps [37, 38, 40]. The simplest of these approximations rely on the fact that mutations at certain positions of the genotype have no effect on the phenotype [37]. These positions are called ‘unconstrained’. Those parts of the genotypes that do affect the phenotype when they are changed are called ‘constrained’.

This technique of sequence constraints can be applied to the biomorphs as follows: The first eight sites in the biomorph genotype encode eight vectors, but not all of these vectors are used in the final shape if the developmental process terminates after a small number of stages, as dictated by gene 9 (Fig 2). Therefore, there are unused vectors and the positions of the genotype that encode such vectors must be fully unconstrained since mutations to these positions can have no effect on the phenotype at all. In our analytic calculations, we assume that all other positions, i.e. positions that affect one or more of the vectors in the final shape in some way, are fully constrained, i.e. that any change in these positions leads to a phenotypic change: this is a simplifying assumption since it is possible that two lines in the biomorph shape are drawn on top of one another, and in this case deleting a piece from one of these lines has no visible phenotypic effect. Thus, this analytic model is only perfectly accurate for a very detailed phenotype description: in the analytic model, any small change in any drawn line corresponds to a phenotypic change. Even if a line that was previously drawn multiple times is now only drawn once, this corresponds to a phenotypic change in the analytic model, and if the shape is rescaled, this also corresponds to a phenotypic change. Thus, the analytic description would be 100% accurate if the biomorphs are drawn with a fixed length scale on a very large screen, if lines that are generated multiple times in the developmental process are drawn as thicker lines, and if length-zero lines are included, for example as a visible dot.

Having determined which sites are constrained and which are unconstrained, we can make analytic predictions for GP map characteristics, such as phenotype frequencies, robustness, and evolvability values (see section S1 in the *Supplementary Information* for detailed derivations). The analytic model complements the computational results since both rely on opposite assumptions: the computational model uses coarse-graining, whereas the analytic model is (overly) fine-grained. In order to compare the data from the two approaches, we restrict the genotypes to the same range of integers in both cases throughout the main text. However, since calculations in the analytic approach are fast, we also use this approach to investigate how the biomorphs GP map would change if we allowed the integer values in the genotype to vary over a wider range. This modification produces qualitatively similar outcomes, as shown in section S3.3 in the *Supplementary Information*.

### Models of evolving populations

To model populations of biomorphs evolving over time, we use the Wright-Fisher model with selection [57] in combination with a GP map, as done in refs. [12, 17]: the fitness of a specific genotype is calculated by mapping it to its phenotype and then using a phenotype-fitness relationship that is fixed for each simulation, as described in the text. Mutations occur at a constant rate *µ* per site at each generation [17]. As an initial condition, we choose a random genotype out of all genotypes that meet the specifications (for example map to a given phenotype) and initialize all individuals with this genotype. To ensure that this choice of initial conditions doesn’t affect our measurements, we follow Schaper & Louis [17] and, for a population of size *N*, let the initial population evolve for 10*N* generations before starting any measurements.

## RESULTS

### Phenotype bias towards simple phenotypes

#### Quantifying the strength of the bias

Having introduced the relationship between biomorph genotypes and phenotypes, the first question is how many phenotypes exist and how many genotypes correspond to each of these phenotypes. In the computational results, there are *≈* 9.8 *×* 10^6^ different phenotypes for the 7^8^ *×* 8 *≈* 5 *×* 10^7^ genotypes that are within the parameter range considered in our analysis (approximately 1.2 *×* 10 different phenotypes in the more fine-grained analytic model). A few examples from the computational approach are shown in Fig 3A: among these are phenotypes that are generated by approximately 10^5^ genotypes, as well as phenotypes that are only generated by two genotypes. These examples illustrate that the biomorph system exhibits strong phenotypic bias: neutral set sizes differ by several orders of magnitude between different phenotypes.

**Figure 3.**
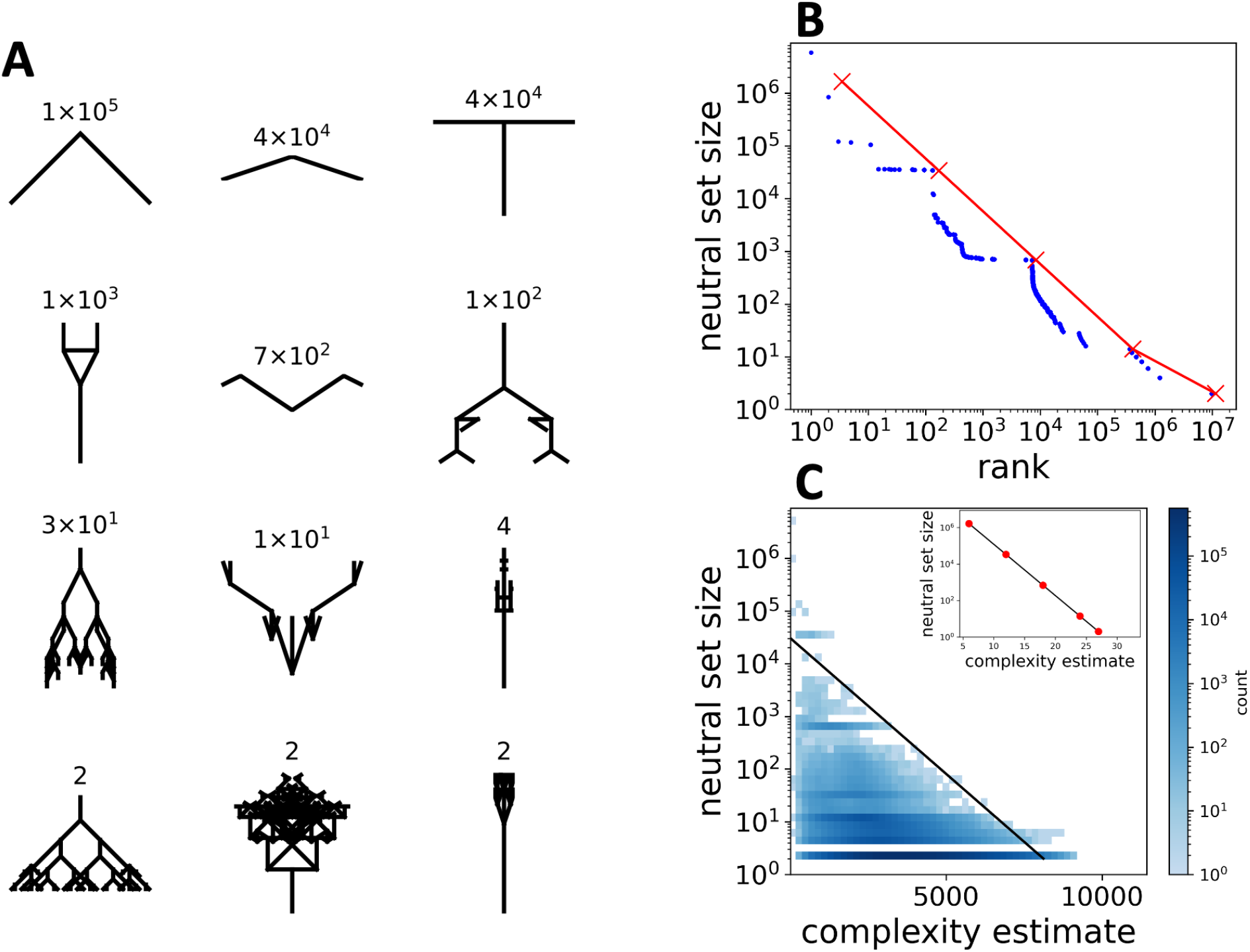
Phenotypic bias: A) Example biomorph phenotypes: the neutral set sizes (i.e. number of genotypes per phenotype) are indicated above each image. The phenotypes shown in the first three rows are chosen to represent a range of neutral set sizes, and the last row shows three phenotypes with a neutral set size of two since *≈* 9 *×* 10^6^ out of *≈* 10^7^ phenotypes have this neutral set size. B) The neutral set sizes of all phenotypes are plotted against their neutral set size ranks (i.e. the number of phenotypes with greater or equal neutral set size). The computational results are shown in blue and the analytic data of Eq (3) in red. In both treatments, neutral set sizes vary over several orders of magnitude, i.e. there is strong phenotype bias. C) The neutral set size of each phenotype is plotted against the estimated complexity of the corresponding coarse-grained binary image (calculated using the block decomposition method [34]). The black solid line is an approximate, but not a perfect upper bound, drawn to illustrate the simplicity-bias prediction from Eq (1). Large neutral set size phenotypes tend to be low-complexity biomorphs and high-complexity biomorphs tend to have small neutral sets (deviations from the upper bound can also be understood within AIT [6]). **Inset:** Neutral set size versus complexity for the analytic model, calculated with the bound of Eq (4) which is based on an alternate complexity measure that measures the size of the constrained part of the minimal genome that generates a given biomorph. The resulting relationship is consistent with the trend in the computational results.

This phenotypic bias can be further observed in Fig 3B where we plot the neutral set sizes for all phenotypes. The sizes vary across more than six orders of magnitude for both for the computational (blue) and the analytic (red) data. Neutral set sizes approximately follow Zipf’s law, where the relationship between neutral set size *N_p_* and phenotype rank *r* (i.e. the number of phenotypes with greater or equal neutral set size) is *N_p_ ∝* 1*/r* for a wide range of *N_p_*. This fat-tailed distribution means that most phenotypes have small neutral sets: in fact, only approximately 4 *×* 10^5^ out of approximately 10^7^ phenotypes have neutral set sizes greater than ten genotypes in the computational results. Note that phenotypic bias is found even without the coarse-graining introduced in the computational analysis, since it is also present in the analytic model, which does not rely on coarse-graining. From the analytic calculations (for details see section S1.1 in the *Supplementary Information*) we find a range of neutral set sizes that depend only on the final site of the genotype *g*_9_:

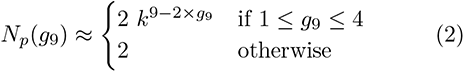

Here *k* = 7 is the number of distinct integers that are in the allowed range for genotype positions *g*_1_ to *g*_8_. Essentially the neutral set size differences in the analytic model are due to the fact that phenotypes with many unconstrained positions can be produced by a large number of genotypes [38]: each constrained site can only take one value within the entire neutral set, but each unconstrained site can take *k* different values and thus each unconstrained site leads to a larger number of distinct neutral sequences, i.e. a higher neutral set size. Specifically, each additional unconstrained site increases the neutral set size by a factor of *k*. Due to this simple relationship between constrained sites and neutral set sizes, there can only be a few phenotypes with large neutral set sizes: it is the constrained positions that define the phenotype, and since phenotypes with large neutral sets only have a small number of constrained positions, only a small number of distinct phenotypes with large neutral sets can exist. This argument gives a relationship between neutral set size *N_p_*and phenotype rank *r* that closely resembles a Zipf’s law (derivation in section S1.2 in the *Supplementary Information*), as in some previous constrained-unconstrained models [38], and is plotted in Fig 3B:

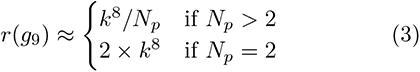

However, note that simplifications were made in the derivation of this equation: the full analytic expression involved a sum over *g*_9_ and we only kept the largest term in each sum. This gives us a simple expression whose *N_p_*-dependence is easy to analyze, but at the cost of underestimating the true rank values.

#### Quantifying the bias towards low-complexity biomorphs

As can be seen visually from the examples in Fig 3A, phenotypes with higher neutral set sizes appear to be less complex. To quantify this trend, we use the block decomposition method [34] to estimate the descriptional complexity *K̃* (*p*) since this method is designed for 2D binary arrays such as our coarse-grained description of the phenotype^3^. This procedure generates the data in Fig 3C.

We find that large-neutral-set-size phenotypes have low complexity, whereas high-complexity phenotypes have small neutral sets. There are phenotypes, which are simple and rare, but we do not find phenotypes that are both complex and frequent. Therefore, the GP map is biased towards a subset of simple biomorph phenotypes. This observation of an upper bound as in Eq (1), with many phenotypes also found below the bound is what Dingle et al. [5, 6] predicted based on their version of the AIT coding theorem. The biomorphs GP map, therefore, presents very similar simplicity bias phenomenology to that found for molecular GP maps in [3]. This conclusion remains unchanged when applying a different Lempel-Ziv-based complexity estimator from [5] to the coarse-grained phenotypes (section S4.2 of the *Supplementary Information*).

In the analytic model, we cannot quantify the visual appearance of a phenotype. Instead, we approximate the complexity of a phenotype by measuring the complexity of a minimal genotype that generates the phenotype. Since not all vectors are used in the final phenotype construction, some are irrelevant and this (unconstrained) part of the genotype has no direct effect on the phenotype. Thus, the full information on the phenotype is contained within the constrained part of the genotype (if the biomorphs construction process is known), and the length of this part of the genotype *K̃* can be used to estimate an upper bound on the description length and hence the complexity. As we have discussed, phenotypes with fewer constrained sites have exponentially higher neutral set sizes. Therefore, the analytic calculations (section S1.3 of the *Supplementary Information*) give a negative log-linear correlation between complexity *K̃* and neutral set size *N_p_* (again with *k* = 7 for the range of values per site):

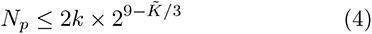

Plotting the analytic neutral set sizes against this estimate of an upper complexity bound leads to the same qualitative conclusions as the computational results (inset of Fig 3C): complex phenotypes have small neutral set sizes, whereas simple phenotypes can have large neutral set sizes. Qualitatively, the conclusions also hold when we quantify the complexity by the number of lines in the biomorph (section S4.3 of the *Supplementary Information*), but the shape of the relationship differs from a simple log-linear curve in this case.

We note that most of the phenotypes Richard Dawkins discusses in his book [1] (for example the ones shown as illustrations) are complex phenotypes, which we estimate to have low neutral set sizes. If all phenotypes of relevance have the same neutral set sizes of (*N_p_ ≈* 2), then there is no bias among these phenotypes. However, in the more general case, where there are no restrictions on which phenotypes evolve, the biases have to be taken into account.

### Further GP map structure that shapes phenotypic variation

Fundamentally, the GP map determines how random mutations produce novel variation. Many molecular GP maps have been shown to share a series of structural features beyond simplicity bias that also shape the spectrum of phenotypic variation [7, 35]. This finding begs the question of whether the biomorph GP map also exhibits these other features.

We will focus on three structural features of GP maps that affect evolutionary dynamics. We first explore mutational robustness which quantifies the likelihood of neutral mutations that leave the phenotype unchanged. Secondly, we study how the mutational robustness of a phenotype correlates with a measure of evolvability that counts how many different unique phenotypes are accessible by point mutations on genotypes within a given neutral set. Thirdly, we analyze the phenotypic mutation probabilities, which measure how likely a mutation will be non-neutral and lead to a specific new phenotype. The definitions of these quantities follow standard practice [17, 35, 36, 58], and are given in table I.

**Table I.** Definitions of key quantities for GP maps: Each line describes one quantity with a symbol and the definition. These definitions are commonly used in the literature [7, 17, 35, 36, 58, 59]. For clarity, we use tildes to distinguish genotypic quantities from corresponding phenotypic definitions.

|  |  |
| --- | --- |
| $N_p$ | The neutral set size of a phenotype $p$ is the number of genotypes that generate $p$. |
| $f_p$ | The phenotype frequency $f_p$ of a phenotype $p$ is the probability that a randomly selected genotype corresponds to the selected phenotype $p$. It is thus a normalized measure of the neutral set size $N_p$ of $p$. |
| $\tilde{\rho}_g$ | The genotype robustness $\tilde{\rho}_g$ of a genotype $g$ is the probability that a random mutation on $g$ leaves the phenotype unchanged. |
| $\rho_p$ | The phenotype robustness $\rho_p$ of a phenotype $p$ is the mean genotype robustness of all genotypes $g$ that correspond to phenotype $p$. |
| $\phi_{pq}$ | The phenotype mutation probability $\phi_{pq}$ from phenotype $q$ to phenotype $p$ is the probability that a random mutation on a random genotype in the neutral set of $q$ leads to a phenotypic change to phenotype $p$. |
| $\tilde{\epsilon}_g$ | The genotype evolvability $\tilde{\epsilon}_g$ of a genotype $g$ is the total number of distinct phenotypes that can be obtained from genotype $g$ through a single mutation. |
| $\epsilon_p$ | The phenotype evolvability $\epsilon_p$ of a phenotype $p$ is the total number of distinct phenotypes that can be obtained from any genotype in the neutral set of $p$ through a single mutation. |

To help quantify these structural features, we use a *random null model* from refs [17, 58] where the neutral set sizes of each phenotype are kept fixed, but the individual assignments of the genotypes to phenotypes are randomized. Comparing to this random null model helps clarify where properties arise from the non-trivial structure in the GP map.

#### Phenotype robustness is high due to genetic correlations

Mutational robustness can be quantified in several ways. Firstly, genotype robustness *ρ_g_* describes what fraction of mutations is neutral for a given genotype *g* [36]. To characterize the robustness of a given phenotype *p*, the phenotype robustness *ρ_p_*of phenotype *p* is defined by averaging the genotype robustness over the neutral set of all genotypes that map to phenotype *p* [36].

In the simple null model with a random assignment of phenotypes to genotypes, one would expect that a mutation on a genotype *g* with phenotype *p* would generate the same phenotype with a probability proportional to the phenotype frequency *f_p_* of *p* [58]. This null expectation is plotted by a solid black line in Fig 4B. However, as can be seen in the same figure, on average we find a completely different scaling, namely that *ρ_p_∝* log(*f_p_*) *» f_p_*. This is seen both in the computational results (blue) and in the analytic (red) calculations. In the analytic calculations, we can rationalize this as follows: each unconstrained site contributes a constant amount of robustness since it can vary freely without changing the phenotype. However, it contributes multiplicatively to the neutral set size since the values at unconstrained sites can be combined in different ways to generate genotypes within the neutral set. Taken together, this gives a log-linear relationship, which is derived in section S1.4 of the SI:

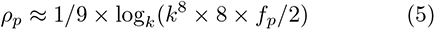

**Figure 4.**
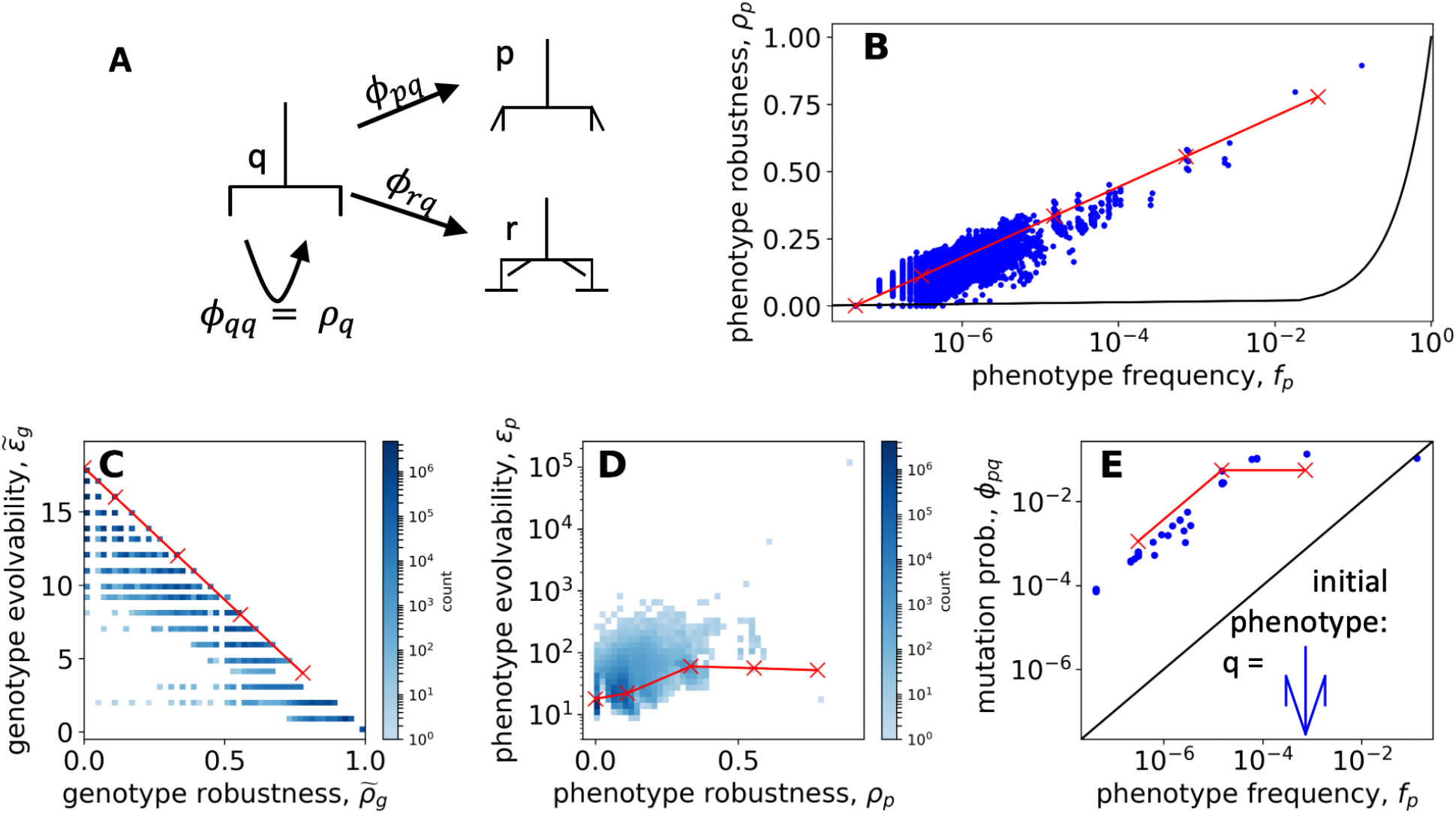
Structure in the GP map - the phenotypic effect of mutations: In every panel, the computational results are shown in blue and the analytic relationships from the constrained-unconstrained model are shown as red lines, with markers indicating the discrete allowed values. (A) Point mutations of a genotype with initial phenotype *q* can either leave *q* intact or lead to a phenotypic change to a new phenotype *p*. The likelihood of the first outcome is given (on average) by the phenotype robustness of *q*, *ρq* . The likelihood of the former outcome is given (on average) by the phenotype robustness *ρq* ; the likelihood of the latter outcome is given by the mutation probability from *q* to *p*, denoted as *φpq* . See Table I for definitions. (B) Phenotype robustness *ρp* vs. phenotype frequency *fp*: the computational results are compared to the prediction of Eq (5). The black line (*ρp* = *fp*) shows the prediction from the uncorrelated null model from refs [17, 58]. The robustness is much higher than this random null model, i.e. there are genetic correlations. (C) Genotype evolvability -*Eg* vs genotype robustness *ρ*-*g* : shows the expected trade-off between robustness and evolvability at the genotype level. The analytic relationship (in red) is from Eq (6). (D) Phenotype evolvability *Ep* vs phenotype robustness *ρp*. The red line is from the prediction of Eq (7). As observed more widely [36], robust phenotypes have large neutral sets and are connected with many other neutral networks, so there is a positive correlation. (E) Mutation probability *φpq* vs. phenotype frequency *fp* for a fixed initial phenotype q (shown in the corner). Again, the black line shows the null model from refs [17, 58], which gives *φpq* = *fp*, The red line denotes a parametric equation from the SI. Data points with *φpq* = 0 are excluded due to the logarithmic scale, even though 99.997% of all biomorph phenotypes have *φpq* = 0 for this particular initial phenotype *q*.

Note that robustness values in the analytic model are discrete because neutral set sizes and hence phenotype frequencies are discrete in Eq (2): the allowed values are *ρ_p_*= 0 and *ρ_p_*= (1 + 2*n*)*/*9 with integer *n* in the range 0 *≤ n ≤* 3.

This log-linear scaling of the robustness and frequency has been reported in many other GP maps [7], including the RNA secondary structure GP map [13, 58, 60], Boolean threshold models for gene regulatory networks [30], a multi-level GP map model called Toylife [61], the polyomino GP map for protein quaternary structure [58, 62], the HP model for protein tertiary structure [58], empirical data on sequences binding transcription factors and RNA binding proteins [63] and the Fibonacci or stop-codon GP map [37] on which the constrained/unconstrained approach is founded. It may hold for a wider set of input-output maps as well [64], and is close to the maximum possible robustness for this class of systems [65]. Because robustness is higher than in the null model in all of these cases, two genotypes that differ only by a single mutation are typically orders of magnitude more likely to correspond to the same phenotype than two randomly chosen genotypes. Such deviations from the (correlation-free) null model have been referred to as *genetic correlations* [58]. The high robustness provided by genetic correlations means that evolving populations can much more easily explore a neutral network than in an uncorrelated model [58], implying enhanced evolvability and navigability of fitness landscapes [66].

#### Phenotype robustness and evolvability are positively correlated

We next analyze the link between mutational robustness and non-neutral mutations. It is clear that there must be a trade-off on the genotypic level [36]. There are only a fixed number of possible mutations per genotype and the more that are neutral, the fewer non-neutral mutations are possible. This trade-off can be quantified by defining the genotype evolvability *c_g_*as the total number of distinct phenotypic changes that are possible through random mutations starting from a given genotype [36]. In Fig 4C) we illustrate this predicted tradeoff between genotype robustness *ρ_g_*and evolvability *c_g_*in the biomorphs system. This pattern is seen both in the computational results (blue) and in the analytic predictions (red) where every non-neutral mutation from a given genotype gives a distinct phenotype, leading to a simple trade-off derived in section S1.5 of the SI:

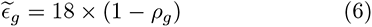

In his “Robustness and evolvability: a paradox resolved” paper, Wagner [36] argued that this picture looks markedly different if we consider the neutral set mapping to a phenotype instead of individual genotypes. A phenotype with high robustness *ρ_p_* is likely to have a large neutral set size. Even if, due to the high robustness, only a relatively small number of non-neutral mutations is possible from each of the genotypes in this neutral set, the higher the number of genotypes, the higher the number of novel phenotypic changes accessible through mutations [36]. This concept is quantified by the phenotype evolvability *e_p_* of phenotype *p* (Table I), which counts the total number of alternative phenotypes accessible from the entire neutral set. We find that, just as for other GP maps [36, 62], this argument holds for the biomorphs GP map: phenotypes with higher phenotype robustness tend to have higher phenotype evolvability. Again, this is seen both in the computational results (blue) and the analytic calculations (red) in Fig 4D.

In the analytic calculation, the positive relationship between evolvability and robustness on the phenotypic level has the following origin: genotypic changes at the unconstrained positions of *p* are neutral and thus occur within the neutral set of *p*. These changes can accumulate and contribute to evolvability because they might become important if a mutation raises the value of *g*_9_ and a new phenotype with a higher number of developmental stages emerges, for which these positions might be important. Thus, different genotypes within the neutral set of *p* can give rise to different phenotypic changes and the evolvability of the neutral set can be higher than the evolvability of an individual genotype in the neutral set. Thus, the phenotype evolvability in the biomorphs system can be higher than the genotype evolvability because unconstrained sites can become constrained (and thus phenotypically relevant) after mutations, as has been shown [40] for other abstract GP map models, including an RNA-inspired model. The full calculation in section S1.6 of the Supplementary Information gives the following relationship:

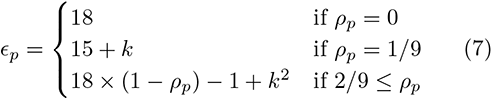

While the trend is the same in the computational results (blue) and the analytic predictions (red), clear deviations between the two approaches exist in the phenotype evolvability calculation. This deviation may be partly due to the nature of the definition of evolvability: all possible phenotypic transitions *p* to *q* contribute equally to *c_p_*, even if they are only possible from a single genotype in the neutral set of *p*. Thus, phenotype evolvability is much more sensitive to small changes in the GP map than quantities like phenotype robustness, which are given by the average over a neutral set.

Note that in both the analytic calculations and the computational results, the phenotype evolvability is typically several orders of magnitude lower than the number of phenotypes (*≈* 10^7^). Thus, while the number of possible phenotypic changes from the neutral set of an initial phenotype can be much larger than the number of possible phenotypic changes from a single genotype, not all phenotypic changes can be achieved in a single mutation even if we include the entire neutral set of an initial phenotype. The reasons for this behavior can easily be understood: for example, if the initial biomorph *q* contains a line pointing in the positive y-direction, at least two point mutations are needed to change this to a vector pointing in the negative y-direction (1 *→* 0 *→ −*1 in the relevant gene).

We hasten to point out that the word evolvability encompasses a much broader set of concepts than the particular measure we discuss above. Evolvability [67–69] is often defined as the potential for “viable and heritable phenotypic variation” [68]. Because many different aspects of biology touch on this capacity, evolvability can be measured in many different ways [70] and thus the genotype and phenotype evolvability measures used here are just one of the ways this concept can be unpacked for biomorphs. Interestingly, although the word appears in the literature at least as far back as 1931 [71], Richard Dawkins’ famous paper on the evolution of evolvability [33], which builds on the biomorphs model, kicked off the modern use of the word [72]. In Dawkins’ paper, he notes that evolvability depends on the developmental process. He contrasts the classic biomorphs studied here with variations to the model that have additional developmental steps, such as segmentation. This perspective on evolvability differs from the one we have analyzed here, where we compare the phenotype evolvability of biomorph phenotypes that all originated from the same fixed developmental system. The rich concept of evolvability has many facets [70].

#### The mean probability φpq of a non-neutral mutation from phenotype q to phenotype p is higher for target phenotypes of high fp

Our phenotype evolvability calculations only tell us how many different phenotypic changes are possible, but not how likely they are. This latter concept is quantified by the phenotypic mutation probability *φ_pq_*, which measures how likely a mutation is to produce phenotype *p*, given that the phenotype before the mutation is *q* [17]. It is an average quantity computed over the neutral set of all genotypes mapping to *q*. The random null model predicts that *φ_pq_* = *f_p_*, indicating that the probability of phenotype *q* mutating to phenotype *p* is largely independent of the source phenotype *q* [17, 58]. This simple prediction has been shown to work reasonably well for several molecular GP maps if the initial phenotype *q* has a large neutral set [58].

Fig 4E plots the mutation probabilities *φ_pq_* for an initial phenotype *q* of intermediate neutral set size (*N_p_* = 3.5 *×* 10^3^ in the computational results, *N_p_* = 6.9 *×* 10^2^ in the analytic model). While it is clear that for accessible phenotypes, *φ_pq_*indeed increases with the frequency of the target phenotype *p*, the data deviates from the simple relationship *φ_pq_* = *f_p_*. One deviation is that most phenotypic transitions are impossible (i.e. *φ_pq_*= 0 and thus these *φ_pq_*values are not shown on this log-log plot): for the initial phenotype *q* shown in Fig 4E, we have *φ_pq_* = 0 for *≈* 99.997% of all possible biomorph phenotypes *p*, and this figure is even higher for other less evolvable choices of *q* – the phenotype *q* in Fig 4E, which is based on a genotype drawn at random from all genotypes with *g*_9_ = 3, has a comparatively high evolvability of 261 phenotypes in the computational results (60 in the analytic model) and thus a higher number of possible phenotypic changes than most other phenotypes. As noted in our discussion of phenotype evolvability, the fact that many phenotypic changes are impossible through single mutations is a feature of the biomorphs system that may not be shared across all GP maps. Interestingly, the allowed phenotypic transitions, i.e. those with non-zero *φ_pq_*, are mostly transitions to phenotypes whose phenotypic frequency is within two orders of magnitude of the phenotypic frequency of *q*. In the analytic model, this is easy to explain: each gene, including *g*_9_ can only vary by *±*1 in a single mutation and thus the neutral set size, which depends on *g*_9_ (Eq (2)), can only vary by a limited amount.

If we consider the possible phenotypic transitions shown in Fig 4E, we find that transitions to target phenotypes with high phenotypic frequency tend to be more likely, i.e. a higher *f_p_* tends to be associated with a higher *φ_pq_*. There is a linear regime (*φ_pq_ ∝ f_p_*), but also a regime at a higher frequency where the relationship plateaus. This pattern is observed both in the computational results (blue scatter points in Fig 4E) and the analytic calculation (red line - this is given by a parametric equation derived in section S1.7 in the *Supplementary Information*). This parametric equation summarizes the following relationships: high *φ_pq_*values are predicted for phenotypic changes to phenotypes with the same or fewer constrained values, which are known to have equal or larger phenotypic frequencies than the initial phenotype *q*. Low *φ_pq_*values are predicted for phenotypic changes to phenotypes with a higher number of constrained values. These transitions are rare because they can only happen on a specific genetic background because of the additional constrained values. Since these phenotypic transitions correspond to a higher number of constrained sites, they have lower phenotypic frequencies than the initial phenotype *q*. While the computational and analytic data show good agreement, the computational data includes additional transitions at very high and very low values of *f_q_*: the transition with *f_q_ >* 10*^−^*^2^ corresponds to the simple ‘line’-shaped phenotype. This phenotype’s neutral set is highly affected by the treatment of overlapping vertical lines along the y-axis and by rescaling, and therefore shows large deviations between the two models. Similarly, the computational data contains additional transitions with low values of *φ_pq_* and *f_q_*. As we argued when comparing evolvability predictions, this is because phenotypic transitions that are only possible from one or a small number of specific genotypes in the initial neutral set are particularly sensitive to a change in GP map definition^4^. These differences between the computational and the analytic data mean that the bias in the mutation probabilities *φ_pq_* is higher in the computational data.

Overall, our main takeaway is that most phenotypic transitions are not possible through single mutations, but out of the possible phenotypic transitions, those to phenotypes with high neutral set sizes tend to be more likely. The second aspect is in agreement with results from a series of other GP maps [58], even though the exact shape of the relationship with its two distinct regimes is different. Because complex phenotypes have low phenotypic frequencies, this implies that the most likely phenotypic changes tend to be towards low-complexity phenotypes (as confirmed in Fig S12 in the *Supplementary Information*). This agrees with previous research that has argued that transitions from high-complexity phenotypes to low-complexity phenotypes are more likely than the reverse, both for an L-system-based GP map [73] and a GP map for digital organisms [27]. However, it is important to note that these arguments only hold for initial phenotypes with a relatively high neutral set size: if the initial phenotype is one of the *≈* 9 *×* 10^6^ phenotypes with a neutral set size of *N_p_*= 2, then there are only 36 possible distinct mutations (18 per genotype for *N_p_*= 2 genotypes), and since typically at least ten phenotypic transitions are found^5^ among these 36 distinct mutations, all non-zero *φ_pq_*values are of a similar order of magnitude and no strong bias among possible phenotypic transitions is expected.

### Phenotype bias and adaptive evolution

Having analyzed what the GP map can tell us about the phenotypic effect of mutations in general, we next investigate how this structure in the arrival of variation affects an evolving population. Modeling evolving populations requires us to make assumptions about the way in which fitness depends on the biomorph phenotype and so we study several scenarios. All data in the following sections rely on computer simulations that use the computational treatment of the biomorphs GP map.

#### Scenario 1: Neutral evolution on a flat fitness landscape

We start with the simplest scenario: a population of size *N* = 2000 evolves under Wright-Fisher dynamics without the effect of selection, i.e. all phenotypes are equally fit and there is only genetic drift. Each individual genotype in each generation could carry any of the approximately 10^7^ different phenotypes, so we simplify our analysis by focusing on three phenotypes with different neutral set sizes, as highlighted in Fig 5A. We recorded each time that one of these phenotypes was found in the population (Fig 5C). Out of these three phenotypes, the one with the highest neutral set size appears most frequently in the population, followed by the phenotype with an intermediate neutral set size. The phenotype with the lowest neutral set size only appears twice. The takeaway from this scenario is the intuitive result that, on average, the rate at which individual phenotypes appear in a neutrally evolving population is well predicted by their global phenotypic frequencies *f_p_*(Fig 5B), as previously seen for molecular GP maps [3, 74]. It is not hard to imagine that these large differences in the rates can also affect adaptive evolutionary scenarios where fitness plays a role, as we will see later in this paper.

**Figure 5.**
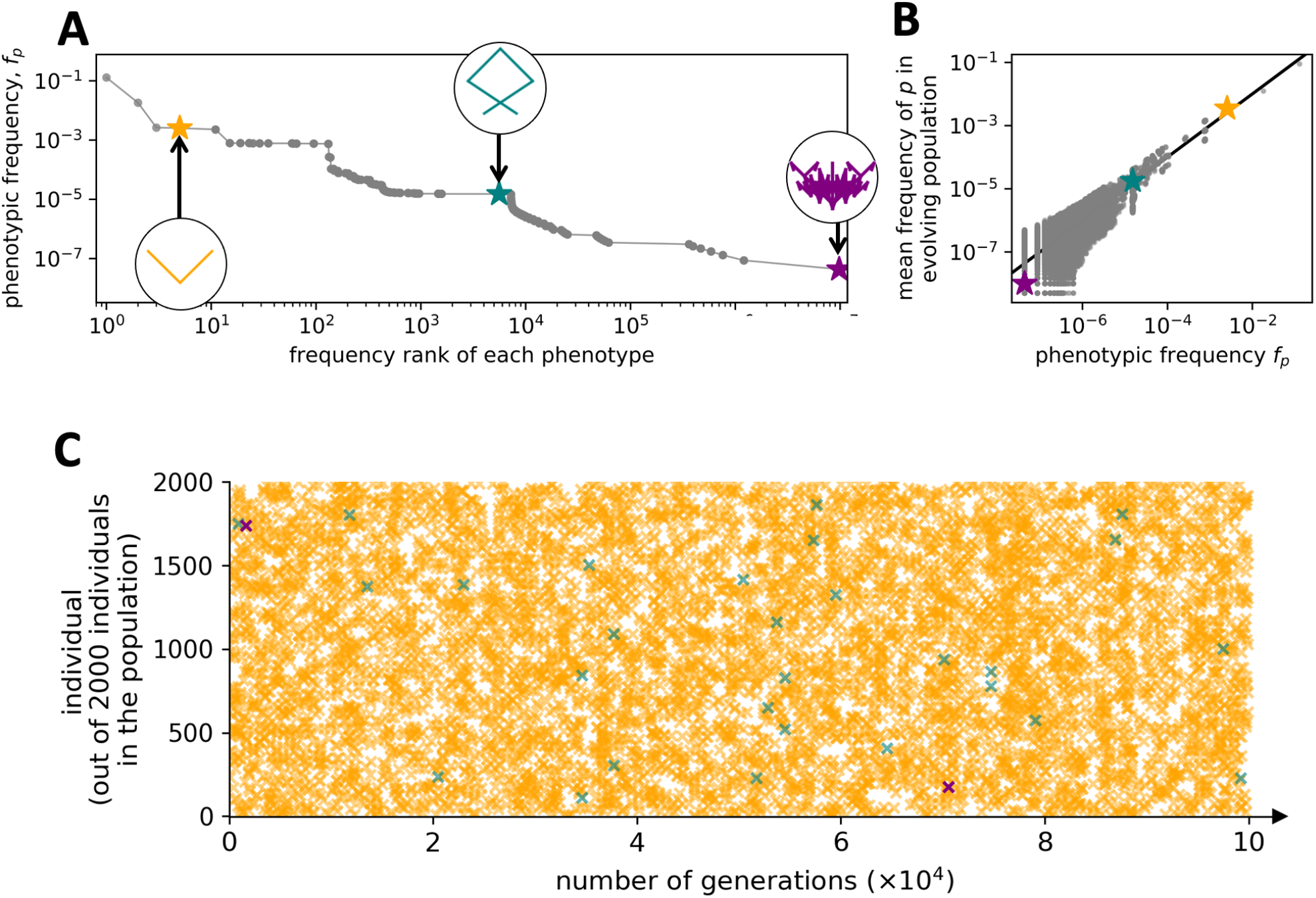
Evolution in a flat fitness landscape. We simulate a scenario where all biomorph phenotypes are equally fit so that there is no selection, just neutral drift. (A) This plot of phenotypic frequency versus rank highlights three phenotypes that are chosen for more detailed analysis: one with high frequency in genotype space (yellow), one with medium frequency (teal), and one with low frequency (purple). (B) The normalized number of times each phenotype occurs in the population is plotted against its phenotypic frequency (for all phenotypes; the chosen phenotypes are highlighted in color). As might be expected in the absence of selection, we see an approximate one-to-one correspondence (as indicated by the black line), with some fluctuations at low values of *fp*. Zero values are not shown due to the logarithmic scale. (C) We plot the occurrence in the population of each of the three phenotypes highlighted in (A) once every 100 generations. This representation highlights the relative frequencies with which the different phenotypes appear in an evolving population of 2000 individuals. The most frequent phenotype (yellow) appears in the population with an average of about 7 individuals per generation. The intermediate frequency phenotype (green) appears in the population on average only once every 28 generations, so about 200 times less frequently than the yellow phenotype. The rarest phenotype (purple) only appears twice in all 10^5^ generations. To take this into account, we plot the two times they appear separately. **Parameters**: Population size *N* = 2000 individuals, with a mutation rate of *µ* = 0.1 per genome, evolving for 10^5^ generations, initialized on a random initial genotype - we run the simulation for 10*N* = 20000 generations before starting the analysis to minimize artifacts of the initial conditions.

A slightly more complex version of this scenario is included in the *Supplementary Information*(Fig S15): here all tree-like biomorph phenotypes are equally fit, but all biomorphs that are not tree-like are unviable. This scenario approximates a situation, where some phenotypic features are under extremely strong selection, whereas others are irrelevant for survival and therefore neutral. Qualitatively, we observe the same trends: there is phenotypic bias over several orders of magnitude among the viable phenotypes and this bias is reflected in the evolving population.

#### Scenario 2: Two peak fitness landscape

Next, following [17, 18], we investigate a more complex adaptive scenario, a two-peak fitness landscape, where two phenotypes have different selective advantages over an initial source phenotype. As illustrated in Fig 6, the population starts at an initial phenotype *p*_0_ and most alternative phenotypes are unviable, with two exceptions, phenotypes *p*_1_ and *p*_2_. For simplicity, the population is chosen such that we are approximately in the strong- selection weak-mutation regime^6^, where adaptive mutations are a limiting factor.

**Figure 6.**
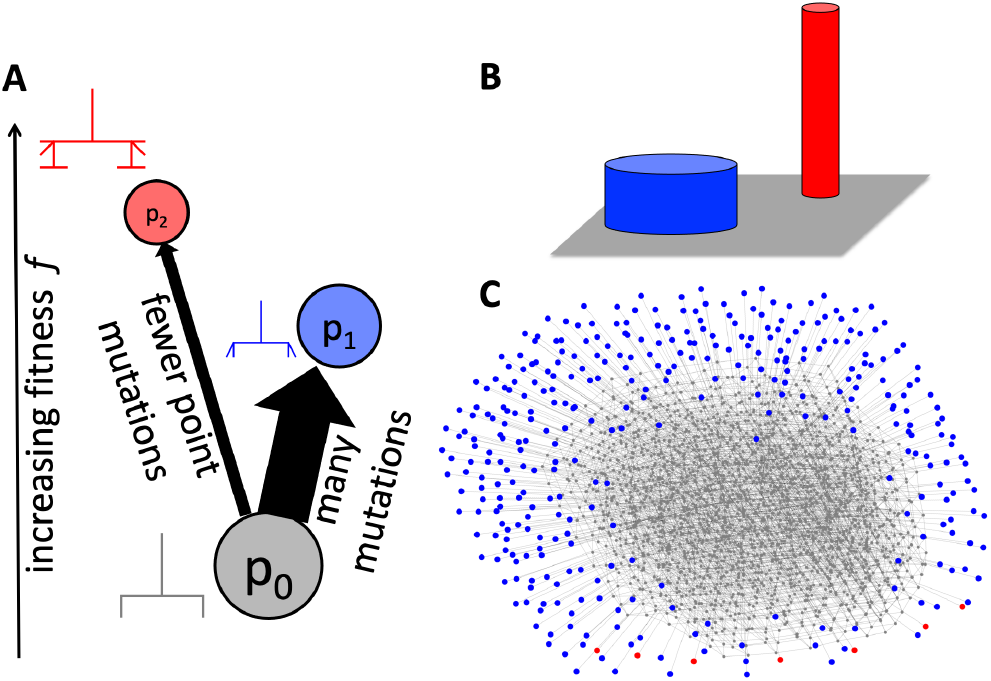
Schematic of the two-peaked fitness landscape (following a similar example for RNA. [17]): (A) Only three phenotypes have nonzero fitness - the initial phenotype *p*0 (grey) and the two adaptive phenotypes, *p*1 (blue) and *p*2 (red). Phenotype *p*1 is more frequent and has a higher mutation probability *φp*1 *p*0 from *p*0 than *p*2 does, but *p*2 has higher fitness. A single point mutation can convert *p*0 into *p*1 or *p*2, but not *p*1 into *p*2. (B) The same scenario is sketched as a schematic ‘fitness landscape’ - the population starts with the *p*0 phenotype (grey area) and there are two fitness peaks, corresponding to *p*1 (blue) and *p*2 (red). While *p*1 forms a broader fitness peak and thus a larger mutational target, *p*2 forms a higher fitness peak. (C) A schematic representation of the relevant part of the GP map: each genotype is drawn as a node in the color of the corresponding phenotype and two genotypes are connected by an edge if one can be reached from the other through a single point mutation. This representation illustrates that there are many different genotypes for each phenotype and that the population will therefore evolve neutrally on the neutral component of *p*0, until moving to either *p*1 or *p*2. Genotypes are only included in this network if they belong to the initial neutral component of *p*0, or if they are direct mutational neighbors of that neutral component and map to *p*1 or *p*2. The initial neutral component can be found by starting from the genotype (-2, 0, 2, -2, 0, 0, -2, -2, 3).

In this particular example, phenotype *p*_1_ has a frequency *f*_1_ = 1.5*×*10*^−^*^5^, and phenotype *p*_2_ has a frequency *f*_2_ = 9.1 *×* 10*^−^*^7^ *≈* 0.06 *f*_1_. However, since the initial condition is known, the relevant quantities are the probabilities of obtaining *p*_1_ and *p*_2_ through mutations from our initial conditions: the whole population is initially undergoing neutral exploration starting on one particular genotype *g*_0_ in the neutral set of *p*_0_ and can therefore drift through the entire part of the neutral set of *p*_0_ that is accessible from *g*_0_ through neutral mutations. This part is known as a neutral component (NC) of *p*_0_ [76]. The phenotype mutation probabilities for that NC determine the rates at which the two adaptive phenotypes are expected to appear [17]: these are also biased towards *p*_1_, with 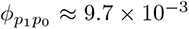 and 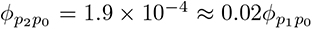. The fitness is traditionally expressed as *F_p_* = 1 + *s_p_* in terms of the selection coefficient *s_p_*. For the neutral phenotype, *s*_0_ = 0, and we vary the two other fitnesses, but we are only interested in the non-trivial case, where the rarer phenotype has larger fitness, in other words, *s*_2_ *> s*_1_ *>* 0.

In our simulations of the fixation dynamics, both *p*_1_ and *p*_2_ can evolve from the initial phenotype *p*_0_ and both are fitter than *p*_0_.If selection alone was the deciding factor, we would expect *p*_2_ to evolve in every simulation since it has the highest selective advantage. However, the more frequent phenotype *p*_1_ also has a selective advantage over the initial phenotype *p*_0_, albeit a smaller one, and so *p*_1_ can reach fixation before *p*_2_ appears in the population as potential variation. Since it is not possible to go from *p*_1_ to *p*_2_ through a single point mutation, but only via a two-step process from *p*_1_ back to *p*_0_ and then to *p*_2_, we focus only on the first fixation event. This is a good approximation since the population is unlikely to go back to *p*_0_ via drift due to the strong selection, as shown in the Supplementary Information. In Fig 7 we analyze how likely it is that (A) the fitter and rarer phenotype *p*_2_ has appeared at least once before the first fixation event and (B) the first fixation event is a fixation of *p*_2_.

**Figure 7.**
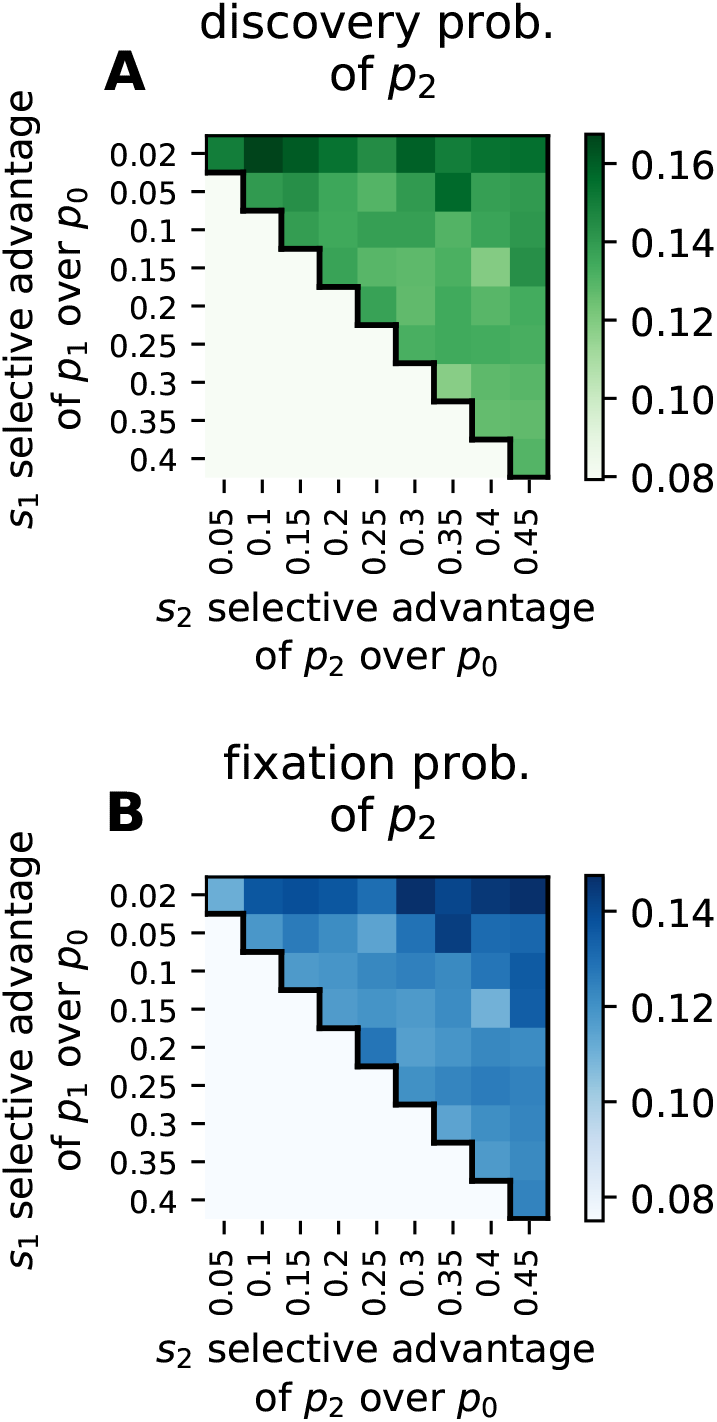
Probability of discovery and fixation of the fitter and less frequent phenotype p2: (A) Probability that the rarer phenotype *p*2 appears in the population before the first fixation event. As long as the more frequent phenotype *p*1 has a sufficiently high selective advantage *s*1 over the initial phenotype, it is likely to appear and fix quickly so that it becomes unlikely that the rarer phenotype *p*2 appears even once before *p*1 fixes. (B) The probability that the rarer phenotype *p*2 is the first to fix. *p*2 can only fix once it appears and so its fixation probability is even lower than the probabilities in (A). These results for the phenotypic bias are consistent with the trends expected from mutational bias [18]. Parameters: population size *N* = 500, mutation rate *µ* = 0.0001, 10^3^ repetitions per parameter set; we consider a phenotype to have fixed if *>* 70% of the individuals carry that phenotype. The populations start from randomly chosen initial genotypes *g*0 that all belong to the NC shown in Fig 6 and then evolve neutrally for 10*N* generations before the three-peak simulations begin.

Fig 7A shows a heatmap of the probability that the rarer phenotype *p*_2_ appears at all in the population before the first fixation event of *p*_1_. This probability is low in the entire range of selective advantages we consider, but it increases slightly if the high-frequency, lower fitness phenotype, *p*_1_ has a low selective advantage (i.e. *s*_1_ = 0.02). This effect occurs because if the high-frequency phenotype *p*_1_ takes longer to go to fixation, this leaves more time for *p*_2_ to appear. Note that the selective advantage of the low-frequency phenotype *p*_2_ does not play a role here: *p*_2_ could be infinitely fit, but when it appears in the population for the first time is unaffected by its fitness. It is clear that *p*_2_ can only achieve fixation if it appears in the simulation at some point, but even if it appears, it could still be lost due to stochasticity. Thus, we now turn to the probability that *p*_2_ reaches fixation (Fig 7B). This probability is of a similar order of magnitude to the probability that *p*_2_ appears, indicating that *p*_2_ is likely to fix once it appears. However, since the fixation probability cannot exceed the probability of discovery, it remains low for the entire range of selective advantages we consider. Interestingly, the impact of varying *s*_1_ and *s*_2_ is not as strong here as in the original paper by Yampolsky and Stoltzfus [18] that first studied such effects, because their calculations focus on a simpler case with only three genotypes. For evolution on GP maps, where there are many genotypes mapping to *p*_0_, the constant-rate assumptions underlying existing work are merely an approximation to the true dynamics [17].

To sum up, in this particular example, the higher rate with which *p*_1_ is introduced into the population due to random mutations dominates over the difference in selective advantage, which would favor *p*_2_. This does not mean that selection does not play a role: selection is the reason why each simulation leads to one adaptive fixation (*p*_1_ or *p*_2_) and due to selection the probability of a *p*_2_ fixation is highest if the selective advantage of *p*_2_ is much higher than that of *p*_1_.

#### Scenario 3: Finding Dawkins’ beetle

The previous subsection analyzed how the balance between selection and phenotype bias affects a single fixation step. In general, however, phenotypic adaptation is a multi-step process, and this is one of the key themes of *The Blind Watchmaker* [1]. Here we re-visit one example which helps highlight the connection between multistep paths in genotype space and fitness landscapes. In the book, Dawkins recounts how he had not recorded the genotype of an insect-shaped phenotype he had observed [1]. When he tried to find the insect-shaped phenotype again by artificial selection, this took a long time, even though he remembered what phenotypes were visited on the original evolutionary trajectory to the insect-shaped phenotype [1]. He explains the difficulty of finding the exact correct phenotype in terms of the shortest evolutionary paths between two phenotypes. Here we revisit this discussion. Since Dawkins doesn’t write down the exact phenotype, we choose to pick one insect shape, a “beetle”^7^, and illustrate one of the paths with the smallest number of mutations in Fig 8A. To stay on this path, the ‘correct’ one out of 18 possible mutations (two possible changes for each of the nine genotype positions if we ignore boundary effects) has to be chosen at each step, so that the probability of obtaining this particular 13 step path is 1*/*18^13^ *≈* 5 *×* 10*^−^*^17^. Of course, there are many other paths that lead from the initial to the final genotype with the mutations arranged in a different order, so that the real probability of obtaining this phenotype by a random walk is closer to its phenotype frequency of 4*/*(7^8^ *×* 8) *≈* 9 *×* 10*^−^*^8^. Clearly, the probability of obtaining the final beetle phenotype by random mutations is extremely small [1]. By contrast, as illustrated by Dawkins’ second infinite monkey example [1], if there is a fitness function that allows each correct intermediate step to increase fitness, then the probability of success can become exponentially larger. Dawkins uses this example to argue that selection by many small steps is much more efficient at finding a fitness maximum than a naive mutationist picture where the final biomorph shape appears directly in a population [1]. One weakness of this example, and one shared schematically by his WEASEL program, is that it relies on a fitness function that is uphill for a large number of intermediate phenotypes.

**Figure 8.**
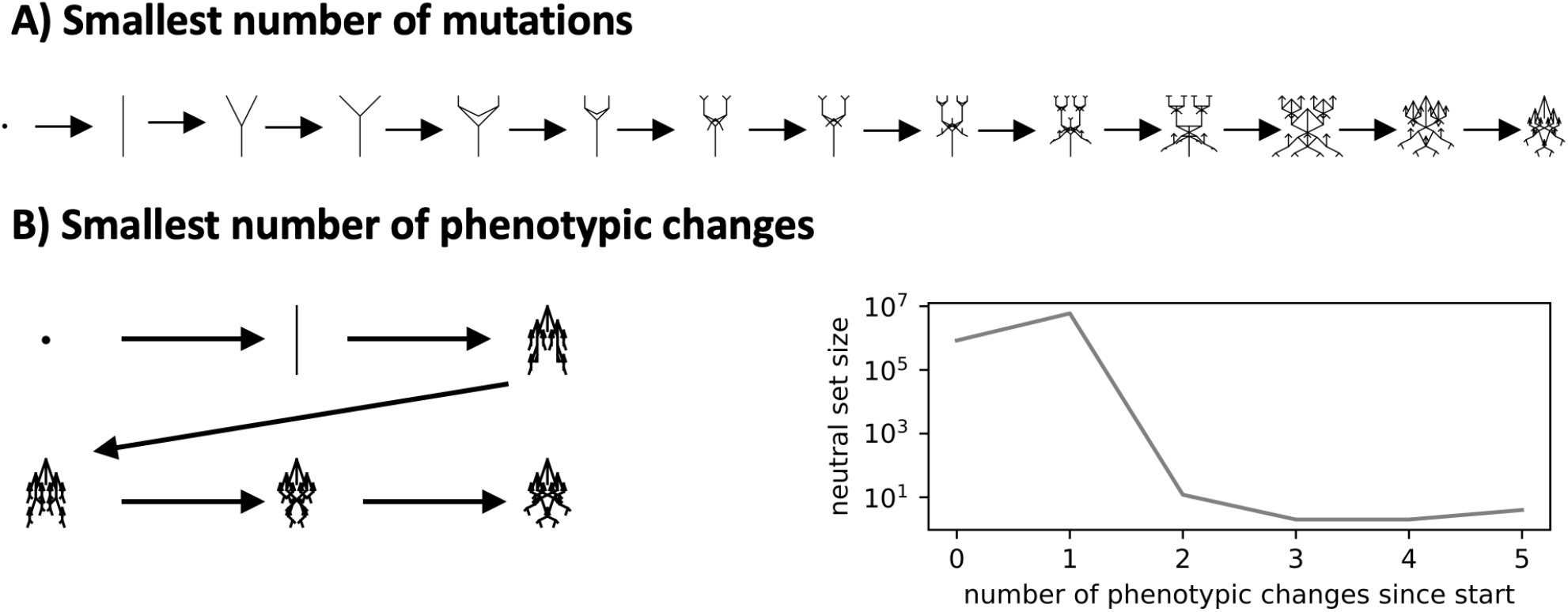
Reconstructing Dawkins’ search for a rare beetle-shaped phenotype: A) An evolutionary path from a dot to the final beetle-shaped biomorph that uses the smallest number of mutations from a given initial genotype that maps to a dot’ phenotype to the final beetle-shape phenotype. B) An evolutionary path from a dot to the final beetle shape with the smallest number of phenotypic changes (but several neutral mutations may be required between each of the phenotypic changes sketched in the figure). The neutral set sizes of the phenotypes along this path are shown on the right. A key advantage of this second scenario is that it increases the probability of a path without any fitness valleys for all intermediate steps (as recently demonstrated for molecular GP maps in Greenbury et al. [66]).

The GP map perspective allows us to study a different kind of minimal path that explicitly includes neutral mutations, and which may facilitate stepwise evolutionary adaptation. Neutral mutations enable genetic drift and cryptic variation [78, 79]. These can facilitate adaptation because, although each genotype in a neutral set maps to the same phenotype, different genotypes will have different sets of accessible alternate phenotypes in their one-mutation neighborhoods [36]. With enough time, a population can in principle explore the entire neutral space, and so find all accessible phenotypes, the number of which is captured by *e_p_*, Wagner’s [36] measure of phenotype evolvability (See Fig 4 (D))^8^. In this context then, rather than ask what the absolute minimal number of mutations is, the more relevant question may be what minimum number of phenotypic changes a population has to pass through in order to evolve from a dot to a beetle. We illustrate an example of such a path in Fig 8B^9^. Allowing the exploration of neutral networks significantly reduces the number of phenotypic transitions from a dot to a beetle when compared to the absolute shortest path shown in Fig 8A. Importantly, such pathways make it much easier to imagine how fitness could increase for all steps since the number of intermediate phenotypes is smaller. This scenario illustrates how neutral correlations in the GP map permit neutral exploration, which may facilitate the emergence of advantageous phenotypic transitions [36, 66]. In this example, concepts related to both the second and the third versions of the infinite monkey theorem defined above interact synergistically.

## DISCUSSION

### The biomorphs GP map shows many similarities to molecular GP maps

GP maps quantify exactly how mutations get translated into a highly anisotropic exploration of the morphospace of phenotypes [3, 9, 80]. A key message of this paper is that the manner in which the biomorphs GP map structures the arrival of variation upon random mutations exhibits many qualitative similarities to what molecular GP maps do [7, 35]. The main similarities observed are listed below:

1. Biomorphs exhibit a strong phenotype bias: upon random sampling of genotypes, certain phenotypes are orders of magnitude more likely to appear than others. However, for biomorphs, a larger fraction of the morphospace of all structures have small neutral sets than is typically seen for molecular GP maps.
2. The particular form of the phenotype bias in biomorphs is typically towards phenotypes with short descriptions. Such simplicity bias [3, 5] means that high-frequency phenotypes have low descriptional complexity, and only low-frequency phenotypes can have high descriptional complexity.
3. The mutational phenotype robustness *ρ_p_* scales as the log of the frequency *f_p_* that a phenotype is obtained upon random sampling of genotypes, and so is much higher than in a random null model without correlations between genotypes.
4. The probability of non-neutral mutations *φ_pq_* tends to increase with increasing frequency *f_p_* of the target phenotype *p*, (if the initial phenotype *q* has a large enough neutral set). However, compared to molecular GP maps [58], biomorphs have an unusually high number of disallowed mutational links between phenotypes, so the positive correlation only holds for the small fraction of phenotypes that are linked by point mutations.
5. 5. The mutational robustness *ρ _g_* of an individual genotype *g* is negatively correlated with a measure of its evolvability *c_g_*that counts the number of alternate phenotypes within a one-mutation neighborhood.
6. By contrast, the mutational phenotype robustness *ρ_p_*, calculated by averaging *ρ_g_* over the neutral set of *p*, is positively correlated with the phenotype evolvability *c_p_*, which counts the number of different phenotypes accessible from the neutral set of all genotypes mapping to phenotype *p*.
7. Many of the relationships above can be analytically derived from a simple model that partitions the genomes into constrained regions that affect the phenotype and unconstrained regions that do not.
8. The many orders of magnitude difference in the rate at which variation arrives in a population can lead to ‘arrival-of-the-frequent’ scenarios [17, 18] where a more frequent, but only moderately fit phenotype will fix in a population because the fitter phenotype either does not appear at all within the relevant time scales, or appears with too low a rate to have a significant probability of sweeping to fixation.
9. Neutral exploration can reduce the number of intermediate phenotypes needed to reach a fitness peak, increasing the likelihood that there are pathways that monotonically increase fitness.

The large number of similarities between biomorphs and molecular GP maps is at first sight surprising since the models have important qualitative differences. The molecular models most studied in the literature are typically based on minimum-free-energy folding (for example protein lattice models [81] and RNA folding models [82]), molecular self-assembly (for example models of protein quaternary structure [62, 83, 84]) or network topologies of gene regulatory networks (for example [30]). By contrast, the biomorphs model’s organization is quite different. It imitates biological development through recursive local branching patterns [1]. The many observed similarities between the manner in which these different GP maps structure the arrival of variation suggest that there may be deeper mathematical principles that are much more widely shared across GP maps. Indeed, the simplicity bias in the GP map follows from very general arguments from AIT, which should hold for a much wider class of input-output maps, of which GP maps are a subset [3, 5]. Similarly, analytic models based on a dual constrained/unconstrained structure of genotypes, while certainly oversimplified, have been used to explain universal behavior observed in molecular GP maps [37, 38, 40]. The fact that this same approach also captures key properties of the biomorphs may help explain why these “universal” GP map characteristics are found for the biomorphs model of biological development as well.

### Can simplicity bias in development be considered as an ultimate evolutionary cause?

If, as we hypothesize above, the phenotype bias towards simplicity observed in molecular GP maps and in the biomorphs model holds for more realistic developmental systems, then this should have implications for the long-standing debate about the relative explanatory power of developmental processes and natural selection [42–46, 48–56]. This debate continues into the present. For example, in a heavily cited recent paper, two teams of scientists argued different sides of the question “Does Evolutionary Theory Need a Rethink?” [45]. The side in favor of the thesis called for developmental bias to be given more weight in evolutionary theory (among other aspects). The other side responded by saying “*Lack of evidence also makes it difficult to evaluate the role that developmental bias may have in the evolution (or lack of evolution) of adaptive traits.*” [45]. While the paper was too short to allow extensive deliberation on this point, it is likely that much of the disagreement is not just about evidence or the lack of it, but rather about (sometimes unacknowledged) differences in their approaches to the interpretation of evidence, influenced by the tangled intellectual history of this debate between internalism and externalism.

Before moving further, we briefly sketch out some of the broader philosophical and historical backdrop to this debate, pointing the reader to Supplementary Text section S9 for a longer (albeit still highly compressed) discussion. The first theme concerns the language of constraints (see also Section S.9.1) which has been heavily influenced by a famous review paper by Maynard-Smith et al. [42] which distinguished between *universal constraints* (such as physical constraints) which act everywhere, and *local constraints* which are limited to certain taxa. Developmental constraints were considered to be local, on the grounds that they don’t hold across taxa. The rise of evo-devo [54, 55, 85–87] means that the term developmental bias is now typically favored over developmental constraints, in part to capture how development can both limit and facilitate certain types of variation [88].

Another longstanding source of confusion in this debate arises from the emphasis by the founders of the modern synthesis of evolutionary biology on the existence of a gene pool of standing variation [89]. In such scenarios, mutations do not supply directionality but only act as fuel [53]. By contrast, in a regime where both the introduction of a new allele by mutation and its fixation or loss are explicitly modelled, biases in the introduction of mutation can substantially affect adaptive outcomes [17, 18, 75, 90, 91]. Detailed evidence for the effect of mutational biases – where certain mutations can be as much as two orders of magnitude more likely than others – on adaptive outcomes has recently been quite firmly established in the field of molecular evolution [92–98]. See Section S.9.2. for further background.

In Section S.9.3. we describe Ernst Mayr’s famous distinction between ultimate “why is it like this” causes of evolutionary biology and proximate “how something works” causes of functional biology [41]. Developmental biases have traditionally been labeled as proximate causes, in part they are thought to ultimately be sculpted by natural selection [99, 100]. How one judges this contentious debate depends in part upon one’s views on contingency [2] (see Section S.9.4.) and on subtle differences between philosophical stances on adaptationism [56, 101, 102] (see Section S.9.3.). Here we build on an approach by Ariew [103] who defines ultimate causes as statistical population-level phenomena that “*answer questions about the prevalence and maintenance of traits in a population*”. These include natural selection, but also other evolutionary causes such as genetic drift. We will add to these evolutionary causes the stipulation that they are universal so that they are not limited to certain taxa. If a developmental bias can be shown to be a universal population-level phenomenon, then it is hard to see why it should not be classed as an ultimate cause.

Interestingly, in the *Blind Watchmaker* [1], Dawkins is happy to concede significant roles to processes such as neutral mutations or developmental bias. Nevertheless, in chapter 11 he writes “*Of course, large quantities of evolutionary change may be non-adaptive, in which case these alternative theories [neutralism, mutationism] may well be important in parts of evolution, but only in the boring parts of evolution . . .* ”. In Godfrey-Smith’s influential three-fold typology of adaptationism [101], Dawkins is an explanatory adaptationist for whom the origins of adaptations are the most important questions in evolutionary biology. On this philosophical stance, even if anisotropies in the introduction of variation significantly affect evolutionary outcomes, they remain “boring” because they are random w.r.t. improvement of the organism and so don’t explain the adaptations Dawkins is primarily interested in.

To prefigure what an argument for simplicity bias in development as an ultimate cause would look like, we will first explore in some detail how this all plays out for the RNA secondary structure GP map where the analysis is simpler and clearer than for larger-scale developmental systems. We then apply the lessons learned to the more ambitious question of what it would take for developmental bias to be classed as an ultimate cause.

#### Phenotype bias as an ultimate evolutionary cause: the example of RNA secondary structures

In addition to the difficulties listed above, the debate on whether developmental bias could be an ultimate cause is hard to resolve because counterfactuals play a key role in the structure of this debate [2], and exploring these reliably is hard. The RNA secondary structure GP map [3, 9, 14], which can be viewed as a stripped-down developmental model [24, 25], has the advantages of being tractable enough to allow counterfactuals to be reliably calculated, and of having abundant and easily accessible data.

#### Constraints and RNA secondary structures

As discussed in the Introduction, strong phenotype bias over many orders of magnitude in the RNA map severely limits the phenotypic secondary structure variation that is accessible to natural selection, the consequences of which can be observed in nature [9, 11–14]. On the one hand, such a strong restriction on the spectrum of potential variation is not that different in principle from the effect of morphological constraints. For example, in the addendum on evolvability in *The Blind Watchmaker* [1], Dawkins argues that: “*You can’t make an elephant by mutation if the existing embryology is octopus embryology* ”. However, these types of constraints are local and may be subject to adaptive change, as Dawkins emphasizes in his writings on the evolution of evolvability [1, 33]. By contrast, we classify simplicity bias in RNA as a universal constraint since it does not depend on the organism, nor does it depend that much on evolutionary history. Moreover, to first order, the bias does not hinge on the detailed chemistry of RNA, but rather arises from deeper information-theoretic principles that hold more widely [5].

The effects of phenotype bias in RNA can then be situated within discussions about the language of constraints and biases [9]: On the one hand, the phenotype bias in RNA ranges over many orders of magnitude, which means that in practice most low-frequency phenotypes are extremely unlikely to ever appear as potential variation on reasonable time scales. In Schaper & Louis [17], this effect was called the “arrival of the frequent”. At this level of analysis, phenotype bias can be viewed as a fairly hard constraint, since it produces a “*limitation on phenotypic variability* ” [42] to an exponentially small fraction of the full morphospace of all biophysically possible structures. On the other hand, within the subset of structures that are frequent enough to appear on evolutionary time scales of interest, there remain differences of several orders of magnitude in the frequencies. These can be said to both enhance or suppress (w.r.t. the mean) the relative rates at which structures that do appear fix in a population [3, 9, 14, 17]. At this second, finer-grained level of analysis, the language of bias seems more appropriate than the language of constraint. The literature on mutation bias [18, 90, 91, 97, 98, 104, 105] is much more similar to this second regime.

*a. Contingency, convergence and RNA secondary structure possibility spaces.* The hyper-astronomically large size RNA sequence space does not necessarily imply contingency for RNA secondary structures, as one might naively think [2]. The RNA GP map exhibits “shape space covering”[82], which means that given a sequence of length *L* mapping to a given structure, the vast majority of common structures are accessible within a relatively small number (*« L*) of mutations. (See Sec. S2 for an analysis of shape-space covering for biomorphs.). Moreover, neutral sets for RNA can have maximal sequence dissimilarity, meaning that completely different RNA sequences might produce the same secondary structure [82]. These properties imply that potentially fruitful RNA secondary structures could appear from almost any genetic starting point.

Similarly, the vast size of the possibility space of all RNA secondary structures does not prevent us from predicting that if one were to replay the tape of life again from the time that RNA appeared, only a minuscule fraction of the set of these possibilities would be utilized, and this convergence would be toward high-frequency structures [9] with low descriptional complexity. At the same time, this picture also implies that as more functional RNA secondary structures are discovered and deposited in databases, we should expect further rare structures to be found.

##### Phenotype bias and selection in RNA secondary structure evolution

Phenotype bias differs from the mutational biases discussed above and in Section S.9 because it can vary over many more orders of magnitude [9]. The fact that much smaller mutational biases can be used to predict adaptive evolutionary outcomes [98] suggests that the much larger phenotype biases will also be predictive in such evolutionary regimes. Indeed, it was shown in [14] that the distributions of inferred neutral set sizes (or equivalently frequencies *f_p_*) of secondary structure databases of functional RNA can be accurately predicted over more than 15 orders of magnitude by random sampling over sequences. Similarly, the inferred distribution of mutational robustnesses closely followed predictions based on the same kind of sampling [14]. In [9], the frequencies of coarse-grained secondary structures of functional RNA sequences from the RNACentral database [16] were directly compared to the *f_p_*’s calculated by random sampling, again finding close agreement over five orders of magnitude for the subset of structures that occur in the database. This close agreement is quite similar in spirit to the correlations found for mutational biases [97], although the range of frequencies is higher, and the agreement is tighter for RNA.

Structures are deposited in a database such as RNA- Central [16] when they are typically thought to have (putative) functions. Obtaining desired RNA function from a randomly picked sequence is highly unlikely (see [22] for a recent discussion). Instead, complex adaptive histories mark each structure deposited in a database of functional RNA. The nature of this selection will have differed from structure to structure. The signatures of such adaptations are likely to average out in database analyses which include many different types of RNA from a variety of organisms. The point is that once these adaptations average out, we are not left with an isotropic pool of structures, as we might expect for evolution without a causal role of bias, but with a sample that closely follows the phenotypic bias. Thus, evolutionary outcomes are shaped both by selection and phenotypic biases^10^.

##### Phenotype bias as an ultimate cause for RNA secondary structures

A good way to tease out the big question of whether or not phenotype bias in RNA should be classified as an ultimate or a proximate cause is to consider the evolution of the hammerhead ribozyme [14], versions of which have been found across the tree of life [108] suggesting convergent evolution. In an important set of in-vitro SELEX experiments, Salehi-Ashtiani and Szostak [109] selected on self-cleaving enzymatic activity, and repeatedly found the hammerhead ribozyme structure, finding convergent evolution in the laboratory. Interestingly, this convergent structure has high phenotypic frequency [14] (and low descriptional complexity) suggesting that it is favored by RNA phenotype bias. Since nature and the SELEX experiments will only have explored a tiny fraction of the morphospace of all possible structures, it is reasonable to expect that, within the vast unexplored space of counterfactuals, there are many alternate structures that could also support self-cleaving function [109], possibly even some with higher self-cleaving catalytic activities than the hammerhead ribozyme.

We can identify two contrasting examples of ultimate causation in this example [2, 14]. Firstly, the ultimate reason why a self-cleaving ribozyme repeatedly appears is of course natural selection for this function. Secondly, and more controversially, the ultimate (population-level [103]) cause for the convergent evolution of the hammerhead ribozyme *structure* is the highly anisotropic manner in which phenotype bias sculpts the variation upon which natural selection can act. Given its profound impact on determining which structures we can observe in nature [9], we would argue that phenotype bias is acting here as an ultimate cause of the emergence of the hammerhead ribozyme structure.

As mentioned above, the phenotype bias in RNA acts both as a constraint, limiting what can realistically appear, and as a bias, influencing the rates at which possible structures fix. Both categories of cause would be ultimate in this case, as they are both universal. By contrast, the situation is more subtle with mutational biases, because detailed mutational biases can vary from organism to organism, and so are less clearly universal (Some mutational biases may be universal, but that is a more subtle question we won’t address here.). If these mutational biases can change under selection, which seems likely, then they may provide an interesting example akin to Dawkins’ principle of the evolution of evolvability. They would then not be classified as ultimate causes by thinkers such as Mayr [99]). By contrast, to first order, simplicity bias is not based on mechanistic detail that may more easily be modified. Instead, it is based on an information-theoretic principle that is much harder to overcome.

### Phenotype bias in development as an ultimate cause?

Finding clear evidence for phenotype bias more generally, or simplicity bias more specifically, in developmental systems will be harder than for molecular systems. Problems typically studied in evo-devo are far from being as tractable or having the abundant data that the GP maps for RNA secondary structures or protein complexes have. Nevertheless, the many orders of magnitude difference in the frequencies found for the biomorph system, which is consistent with what was found for other GP maps and for input-output maps more generally [5], suggests the hypothesis that similar strong bias in the generation of variation may be present for other developmental systems. Moreover, the fact that this bias plays out so clearly in adaptive outcomes for functional RNA, and the fact that mutational biases can also significantly affect adaptive outcomes for a wide range of organisms, suggests that there may be many evolutionary regimes where even rather modest biases for developmental systems will similarly affect adaptive outcomes. If this is true, it has many interesting evolutionary implications. For example, strong developmental bias could be an important explanatory factor for certain kinds of parallelisms and/or convergences, which are commonly found [110] in the history of life, just as was argued above for the structure of the hammerhead ribozyme, and has been found for mutational biases on the sequence level [93]. In summary, phenotype bias in a developmental system may then as we argued for RNA, function in some cases more like a constraint, limiting the arrival of variation to a small fraction of the space of all theoretically possible phenotypes the system can produce. And, on the other hand, it may act more subtly as a bias, both enhancing and suppressing the relative frequencies with which phenotypes that do appear are fixed.

But even if the hypotheses above about strong developmental bias affecting evolutionary outcomes turn out to be correct, the question of whether or not developmental bias is an ultimate cause remains more ambiguous than it is for RNA. One key conceptual difference is that the RNA GP map describes a system that appeared so early in evolution that it is hard to imagine life without it. By contrast, morphological developmental systems such as those underlying body plans are relatively much more recent. One can more easily imagine an evolutionary history where a particular developmental system does not occur, or one where it can change over time due to natural selection, just as may be the case for some mutational biases. Whether one considers such specific developmental biases to be candidates for ultimate causes will also depend on one’s stance on adaptationism and contingency, further complicating this question.

Nevertheless, if our general hypothesis holds, then even though the details of the bias in each developmental system would differ in ways that may be shaped by natural selection, they would each still exhibit an overall universal phenotype bias towards simplicity in the variation they produce. This would make simplicity bias a universal constraint in the language of Maynard-Smith et al. [42]. It would exist across taxa and not be a product of selective pressures. Depending on the exact population genetic regime, developmental biases may strongly affect what fixes in a population, in which case a general developmental bias towards simplicity fulfills our stringent criteria for being an ultimate cause.

What kind of evidence would one expect to find if simplicity bias is at play? One example where it has been invoked as a non-adaptive explanation is for the prevalence of high symmetry protein complexes [3]. The basic idea is easy to understand from the algorithmic picture of evolution. Since less information is needed to describe bonding patterns that lead to higher symmetry, such phenotypes are much more likely to appear upon random mutations [3]. One could easily imagine extending this preference for symmetry to larger-scale developmental processes (see [111, 112] for a discussion).

More generally, we hypothesize that any biological process that can be understood from an algorithmic perspective – consider for example branching morphogenesis [113] – should exhibit a bias towards simplicity, e.g. towards processes that can be described by shorter algorithms which are easier to find by random mutations. In some cases, this bias would be evidenced by the prevalence of symmetries, modulated by processes such as symmetry breaking [114]. In other cases, including the RNA secondary structures and branching morphologies, different signatures of simplicity need to be employed as evidence for this bias. An interesting line of research that needs further development is exactly how to characterize and analyze such derivative signals. For example, simplicity bias also predicts that random mutations should lead to simpler structures. Indeed, experiments on developmental pathways for mouse teeth suggest that mutations leading to simpler tooth shapes are more common than those that lead to increased tooth complexity because the latter scenario requires a coordinated change in several pathways [115]. The consequences of such simplicity bias for morphological changes are also discussed in an exciting recent study on shark teeth [116]. And finally, phenotypic changes observed in phylogenies of angiosperm leaf shapes [117] tend to be strongly biased towards simpler phenotypes.

### Future work for the biomorphs model

In his work on evolvability, Dawkins used the biomorphs “*as a generator of insight in our understanding of real life*” [33]. We believe that this tractable toy model of development has been understudied in the literature, and show that the biomorphs GP map displays a remarkably rich structure in the mapping from genotypes to phenotypes. These discoveries suggest a number of new directions in which our work on biomorphs could be extended. First of all, for computational efficiency, we only used a specific version of the model with nine genes, the same number that Dawkins used in *The Blind Watchmaker*. But the number of genes can easily be expanded, and several of the rules can be adapted [33]. Such changes to the genotype structure and the phenotype construction can allow the model itself to evolve, in other words, future simulations should not just model evolution *on* the GP map, but also evolution *of* the GP map, as advocated in ref [35]. With such an approach, one could study Dawkins’ formulation of the evolution of evolvability [33] and link it to some of the other ways that the concept evolvability is used [69]. For example, certain types of structure in the arrival of variation may facilitate the evolution of phenotypic novelty [118, 119]. Such changes to GP maps are likely candidates for being under positive selection, and biomorphs may form a good model system to investigate some of these proposals. These investigations could be supplemented with a second toy model introduced by Dawkins, the arthromorphs from his book “Climbing Mount Improbable” [120]. The arthromorphs produce a range of segmented 2D body plans inspired by arthropods such as Derocheilocaris [120]. Since the arthromorphs model produces its shapes from numeric genotypes much like the biomorphs system does, it may be that the same tools we develop here could be used to analyze not just development within one bodyplan, as for the biomorphs, but also the evolution of bodyplans.

Much of the astounding progress in the field of evodevo revolves around understanding key genes and genetic pathways that affect development. These are highly conserved and are often given evocative names [54, 55, 85, 87]. By contrast to the RNA model, where the exact identity of the mutations is clear, in the biomorphs model, the mutations act on parameters and do not have as clear a biological identification. This more coarsegrained approach presents a challenge for modeling developmental systems [78]. Nevertheless, schematic models such as the biomorphs model have a long track record of success in evo-devo. Perhaps the most famous are recursive growth models that have successfully been used to study developmental bias in plants [121]. Interestingly, gene-regulatory networks may also generically exhibit simplicity bias [30] and can display arrival-of-the- frequent like phenomena [122, 123]. Nevertheless, further work is needed to connect the results of schematic models to the underlying gene-regulatory networks.

Another direction for future research would be to look at the likelihood of phenotypic transitions (*φ_pq_*) in more detail. We found that transitions to high-neutral set size phenotypes tend to be among the most likely transitions, but also that many transitions are not possible in a single mutation so that *φ_pq_* = 0. Future work could investigate whether these impossible phenotypic changes correlate with larger visual changes than possible phenotypic changes do. Recent arguments from algorithmic information theory [124] predict that phenotypes with smaller conditional complexity *K*(*p|q*) (so phenotypes that are more similar to one another) are more likely to be connected by mutations. It is reasonable to expect that a mutation-induced change between more similar phenotypes will result in smaller fitness differences, lowering the probability of deleterious mutations, and increasing the likelihood of finding pathways with small incremental changes. Such correlations between the likelihood of phenotypic changes and their fitness are essentially what Dawkins exploited in the artificial selection experiments in The Blind Watchmaker [1]. By making incremental changes, he was able to evolve rare high-complexity structures such as his insect-shaped phenotypes. It would be interesting to study in more quantitative detail the interplay of random mutations and these phenotypic correlations on incremental adaptive evolution for biomorphs. This research program would entail combining the power of natural selection, demonstrated by Dawkins’ 2^nd^ infinite monkey theorem, with an algorithmic account of how structured variation arises, illustrated by the 3^rd^ monkey theorem. Such an interplay can help illustrate that phenomena such as developmental bias and natural selection are not in opposition, but should rather be seen as dual causes in a richer explanatory landscape. We believe that taking both creative forces into account should be far from “boring”. Instead, their interaction opens up exciting new avenues for understanding how the remarkable power of evolution generates “*endless forms most beautiful* ” [23].

## ACKNOWLEDGMENTS

The authors acknowledge useful discussions with Charlie Hamilton, James Malone, Joshua Payne, Joshua Sharkey, and Malvika Srivastava. We thank Richard Dawkins for pointing out the arthromorphs model to us.

## DATA AVAILABILITY

The code for this analysis can be found at https://github.com/noramartin/biomorphs_GPmap.

## Supplementary Information

### S1 Analytic constrained-unconstrained model

In the biomorphs model [1, 2], each integer in the genotype either affects the vectors from which the figure is built or the number of developmental stages after which the developmental process terminates, as illustrated in Fig 2 in the main text. The assumption we make when modelling the GP map analytically is that a mutation changes the phenotype if and only if it either affects a vector that is used in the final figure or if it changes the number of developmental stages. As discussed in the main text, this is accurate in an extremely detailed representation of the biomorphs, where the biomorphs are drawn with a fixed length scale on a very large high-resolution screen, lines that are generated multiple times in the developmental process are drawn as thicker lines and length-zero lines are somehow represented in the figure, for example as a dot. Otherwise, our assumption is simply a well-motivated approximation that we will use in the following to determine analytically whether two arbitrary genotypes share the same phenotype.

With this ansatz, we can build an analytic model similar to previous analytic GP map models, which rely on a division of sequences into constrained and unconstrained parts (for example in [3–5]): for a given phenotype, there are some positions in the genotype, where any mutation leads to a phenotypic change (constrained positions). Other positions can mutate without changing the phenotype (fully unconstrained positions). These definitions allow us to investigate the GP map analytically.

In our analytic treatment of the biomorphs GP map, we have the following division into constrained and unconstrained parts: *g*_9_ is always constrained since mutations in the value of *g*_9_ change the number of developmental stages and thus always lead to a phenotypic change. Since *g*_9_ is constrained for all phenotypes, all genotypes in a given neutral set have the same value of *g*_9_. The remaining eight genes, *g*_1_ - *g*_8_, which define the vectors in the biomorphs construction process, are constrained only for some phenotypes: whether the vector(s) encoded by a certain genotype position *g_i_* appear in the final figure, depends on the value of *g*_9_. If the vector(s) appear in the figure, any mutation to *g_i_* changes the phenotype, and *g_i_* is fully constrained. If the vector(s) do not appear in the figure, mutations to *g_i_* have no effect on the phenotype in the analytic model and *g_i_* is fully unconstrained. Thus, we can deduce the number of constrained positions *n_u_* by studying Fig 2 in the main text and counting, how many genotype positions only appear in vectors that are not used in the final figure. This only depends on the number of developmental stages and thus on the value of *g*_9_ in the neutral set of the given phenotype:

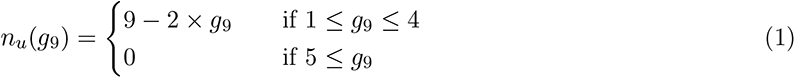

Thus, *g*_9_ sets the number of unconstrained positions in a genotype and plays a similar role to the stop codon in existing analytic constrained-unconstrained models (for example refs [3, 5]).

#### S1.1 Neutral set sizes

The neutral set size can be computed following ref [4] if we know the number of unconstrained positions: in the neutral set of a given phenotype, each constrained genotype position is the same for all genotypes and each unconstrained position can take on any value. In our case, the values in the ‘vector’ part of the genotype are restricted to integers between *−*3 and 3, so unconstrained positions can take one of *k* = 7 values. Thus, there are *k^nu^*^(*g*9)^ possible sequences for the unconstrained part of the genotype.

Hence, there are *k^nu^*^(*g*9)^ different genotypes that give the same phenotype, based on the constrained- unconstrained calculations. In addition, the biomorphs system has an axial symmetry that applies even to the constrained parts of the genotype^1^: flipping all x-coordinates in the figure does not change the phenotype. We approximate this by including a factor of two in our neutral set size estimates. This is only an approximation since the factor of two should not be applied if a genotype had zeros at all x-coordinates, but it gives the following simple expression for the neutral set size *N_p_*(*g*_9_):

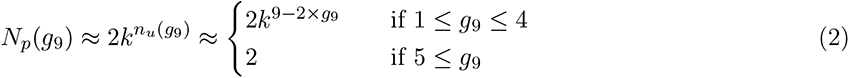

For our plot in the main text, we also need to compute the rank, i.e. the number of phenotypes with greater or equal neutral set size. We can deduce the rank as follows: since neutral set size decreases monotonically with *g*_9_ (eq 2), we can express the rank as a sum over *g*_9_. Since there are *k*^8^ different genotypes for each fixed value of *g*_9_ and *N_p_*(*g*_9_) genotypes per phenotype, there should be *k*^8^*/N_p_*(*g*_9_) different phenotypes for a fixed value of *g*_9_. With this, we can simply sum over all values of *g*_9_ with greater or equal neutral set size to compute the rank. Because the neutral set size is the same for all 5 *≤ g*_9_, we need to handle this case separately:

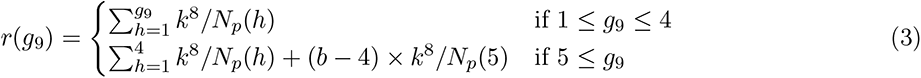

Here, *b* is the number of different values *g*_9_ can take (we assume that one is the lowest allowed value for *g*_9_): in our case, *g*_9_ can take any value from 1 to 8, so we have *b* = 8 and *b −* 4 = 4.

We can simplify the rank calculation in eq 3 by noting that the terms with the smallest neutral set sizes dominate the sums and putting in the expressions for *N_p_* from eq 2. This means that we can approximate the full expression as:

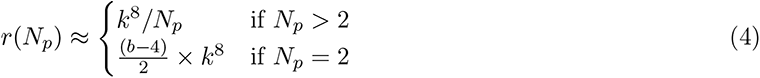

Thus, for a range of neutral set sizes, the rank is proportional to *N ^−^*^1^, and conversely, the frequencies are proportional to 1*/r*(*N_p_*) and so the distribution follows Zipf’s law. This relationship is reminiscent of the power laws found in other GP maps, such as the Fibonacci model [3], for which the constrained/unconstrained approach was first developed. However, note that while eq 3 is exact for the analytic model, the reductions of the sums to their largest terms, which gave eq 4, are only an approximation that will lead to underestimates of the true sums, and thus the true ranks.

#### S1.3 Complexity estimates

In our analytic calculations, we do not draw the biomorphs figures in 2D and so it is not possible to estimate their complexities from the drawn images. However, it is possible to derive an upper bound on the complexity of a biomorph phenotype without drawing the corresponding figure as follows: one way of describing a biomorph phenotype is by recording one corresponding genotype as well as the instructions on how to generate the figure from the genotype. As in previous applications of AIT arguments to GP maps [7], we will ignore the second part because this is a constant term that is the same for all phenotypes. The genotypes, however, are different for different phenotypes: since only the constrained genotype positions are required to fully define the phenotype, the length of the essential part of the genotype varies from phenotype to phenotype. The length of this essential part is proportional to the number of constrained positions per genotype. This is one upper bound on the complexity since the complexity is defined as the shortest possible description length and the genotype is one way of describing the phenotype. Thus, we have an upper bound on complexity *K̃*

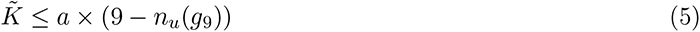

where *a* is the constant of proportionality that is set by the description length per encoded phenotype position. In our analysis, sites *g*_1_ to *g*_8_ can take one of seven discrete values and *g*_9_ can take one of eight discrete values - thus any genotype position can be encoded in *a* = log_2_ 8 = 3 bits. Using eq 2 to express *n_u_* in terms of neutral set sizes then gives a log-linear upper bound:

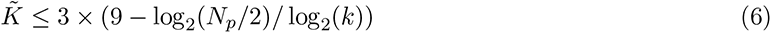

Rearranging for *N_p_* gives:

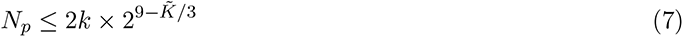

#### S1.4 Genotype and phenotype robustness

In order to find the robustness of a genotype *ρ_g_*, we need to compute the fraction of mutations that leave its phenotype unchanged. By definition, all mutations at unconstrained sites leave the phenotype intact, whereas all mutations at constrained sites change the phenotype. Thus, we only need to know the number of constrained sites and the number of possible mutations at each site. The number of possible mutations at each site is two (changing the value at the site by +1 or -1) if we neglect the fact that in our model, the values are confined to fixed ranges (for example *−*3 is the lowest value *g*_1_ can take and cannot be decreased further). With this assumption, we have:

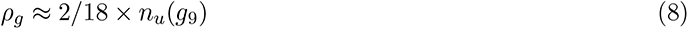

This only depends on *g*_9_ and all genotypes in the neutral set of a phenotype share a single value of *g*_9_, all genotypes in a neutral set have the same robustness. Thus, the average robustness of a phenotype is simply:

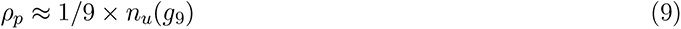

So far, we have written phenotypic robustness *ρ_p_* as a function of *g*_9_. We can rewrite this as a function of phenotypic frequency using eq 2 and relying on the fact that the phenotypic frequency is simply the neutral set size normalized by the total number of genotypes, which is *bk*^8^:

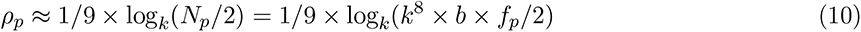

#### S1.5 Genotype evolvability

If we start with a given initial genotype, each distinct mutation in a constrained site changes the phenotype in a distinct way and so there are 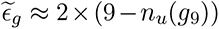 distinct phenotypic changes (again assuming there are always exactly two mutations at each site). Using Eq 9, we have the following relationship between genotype robustness and evolvability:

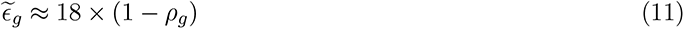

#### S1.6 Phenotype evolvability

Calculating the phenotype evolvability is a little more complex: for this, we need to compute how many distinct phenotypic changes are possible from the entire neutral set of the initial phenotype *p*, a task similar to calculations in ref [5]. This can be higher than the evolvability of an individual genotype if different phenotypic changes are possible for different genotypes in the neutral set of *p*. In our biomorphs model, this is the case for mutations that raise *g*_9_ by +1: since such a mutation adds one recursion in the construction process, the mutation can cause additional vectors to appear in the figure. This means that there can be sites, which were unconstrained before the mutation, but play a determining role for the phenotype after the mutation. Thus, a single mutation - raising *g*_9_ by +1 - can generate several distinct phenotypic changes when applied to different genotypes in the neutral set of *p*.

The number of distinct phenotypic changes that can be achieved from a given initial phenotype by raising *g*_9_ by +1 can thus be computed by identifying the number of sites that switch from being unconstrained to constrained, (9 *− n_u_*(*g*_9_ + 1)) *−* (9 *− n_u_*(*g*_9_)), and the number of values each of these sites could take, *k* = 7. Thus, we have 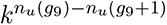 distinct phenotypic changes that can be achieved from a given initial phenotype by raising *g*_9_ by +1.

All other non-neutral mutations - lowering *g*_9_ or changing a constrained ‘vector component’ site - have the same phenotypic effect regardless of which genotype in the neutral set of *p* they are applied to. The number of such non-neutral mutations is 2 *×* (9 *− n_u_*(*g*_9_)) *−* 1. Thus, we can add both contributions to obtain an expression for the phenotypic evolvability of *p*:

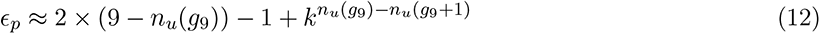

The first term of this expression scales like the genotype evolvability and is anti-correlated with robustness. It is therefore the last term that gives us a positive correlation between robustness and evolvability on the phenotypic level. As postulated in previous theoretical work [5], this term is due to mutations that change the sequence constraints. This role corresponds to mutations in the ‘stop codon’ [5], and thus to mutations in *g*_9_ in the biomorphs model.

So far, eq 12 is only a parametric equation. However, we can put in values of 1 *≤ g*_9_ *≤* 8 and use equations 1 & 9 to get the following cases, depending on the value that *n_u_*(*g*_9_) *− n_u_*(*g*_9_ + 1) can take:

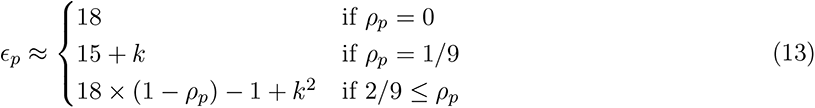

#### S1.7 Mutation probabilities

Mutation probabilities *φ_pq_* quantify the likelihood that a new phenotype *p* is generated by a random mutation applied to a random genotype in the neutral set of an initial phenotype *q* [8]. Both mutation probabilities and evolvability values capture phenotypic changes and thus our calculations of mutation probabilities *φ_pq_* will use the same arguments as in the previous paragraphs: Most mutations bring about the same phenotypic change for all genotypes in the neutral set. Phenotypic changes produced by such mutations are generated by one in every eighteen mutations since there are eighteen mutations for each genotype and only one mutation gives the specific phenotypic change. The one exception is again a mutation that raises the value of *g*_9_: since there are *k^n^^u^*^(*g*9)^*^−nu^*^(*g*9+1)^ different phenotypic changes produced by this type of mutation in a given initial neutral set, each of these occurs with *φ_pq_ ≈* 1*/*(18 *× k^n^^u^*^(*g*9)^*^−nu^*^(*g*9+1)^). To sum up, the possible phenotypic changes have the following likelihood:

*•* Decreasing *g*_9_ by one: this gives a new phenotype *p* with *φ_pq_ ≈* 1*/*18. The neutral set size of *p* is given by *N_p_*(*g*_9_ *−* 1).
*•* Increasing *g*_9_ by one: this gives a new phenotype *p* with *φ_pq_ ≈* 1*/*(18 *× k^n^^u^*^(*g*9)^*^−nu^*^(*g*9+1)^). The neutral set size of *p* is given by *N_p_*(*g*_9_ + 1).
*•* Mutating any other non-neutral site: this gives a new phenotype *p* with *φ_pq_ ≈* 1*/*18. The neutral set size of *p* is given by *N_p_*(*g*_9_).

These three combinations of *φ_pq_*and neutral set size give the analytic prediction plotted in the main text (where the initial phenotype has *g*_9_ = 3). It is important to note that not all phenotypic changes are possible: even if *q* and *p* share the same value of *g*_9_, there is no way of mutating from *q* to *p* if one of the constrained sites differs by *≥* 2 since constrained sites are constant in an entire neutral set by definition and can only change by *±*1 in a single non-neutral mutation. Thus, the analytic model predicts that most phenotypic changes cannot be achieved in a single mutation (i.e. *φ_pq_* = 0).

Thus, frequent phenotypic changes occur with *φ_pq_ ≈* 1*/*18 and correspond to changes in the vector part of the genotype or to changes in *g*_9_ that lower the number of developmental stages, i.e. broadly examples of heterometry or a specific type of heterochrony if we follow the classification of developmental changes in ref [9] (depending on how we assume that the developmental stages in the biomorphs system are controlled by gene-regulatory events). Rarer phenotypic changes occur with *φ_pq_ ≈* 1*/*(18 *× k^n^^u^*^(*g*9)^*^−nu^*^(*g*9+1)^) and correspond to changes in *g*_9_ that increase the number of developmental stages, i.e. to a specific type of heterochrony.

### S2 Shape space covering property

**Figure S1:**
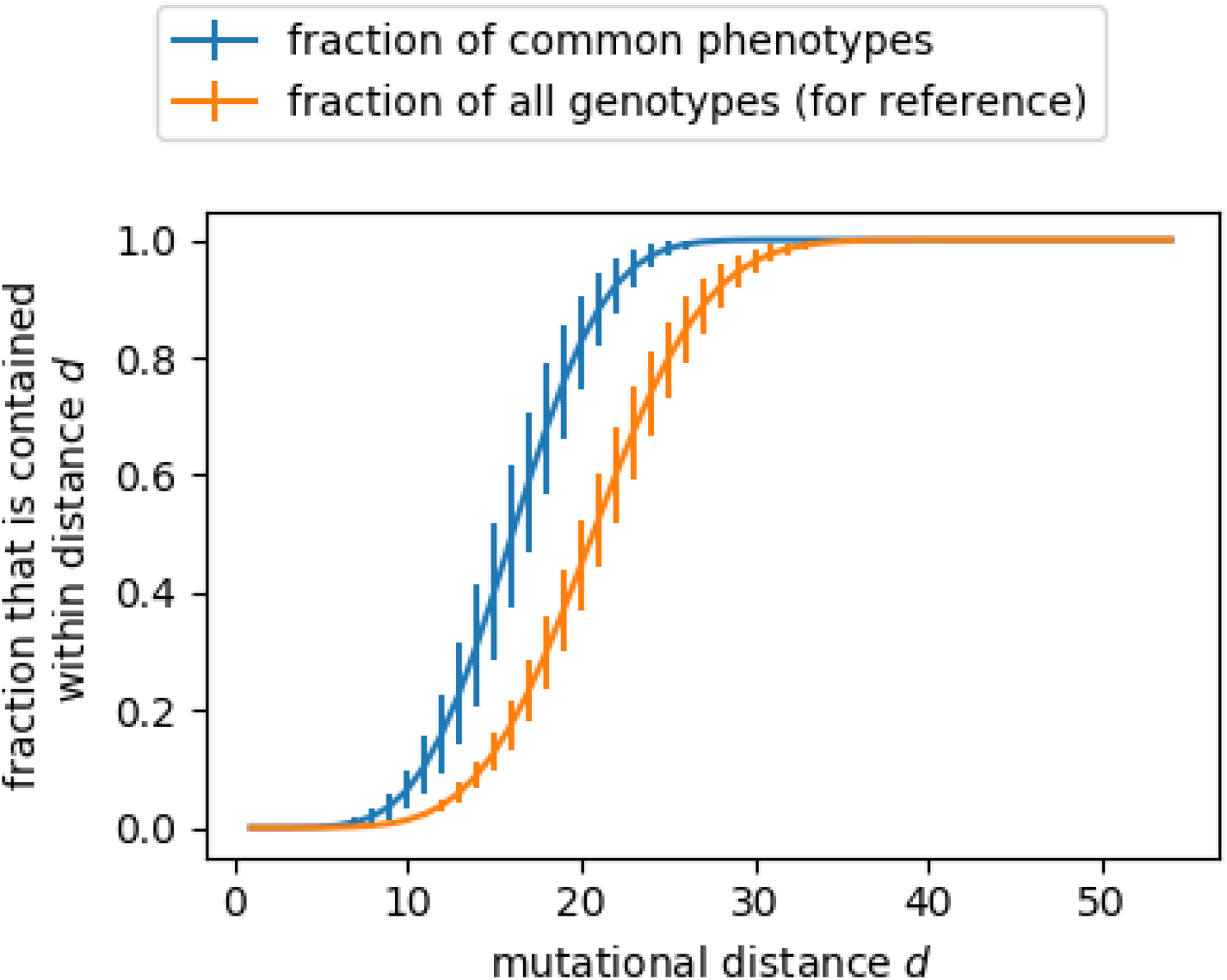
Test of the shape space covering concept: For ten randomly selected initial genotypes, all genotypes within a mutational distance of d are enumerated and their phenotypes recorded. We evaluate, what fraction of high-frequency phenotypes are found within this set for a given value of d (blue). For reference, we also show, what fraction of all genotypes are contained within this set (orange). The plot shows the mean and standard deviation of the values for the ten different initial genotypes. Note that the fraction of all genotypes that are contained within a distance d depends on the initial genotype due to our definition of the biomorphs’ genotype space: if the initial value at a given site is −3, then it can take up to six mutations to reach every value in the valid range [−3, 3], whereas this would only take up to three mutations if the initial value was 0.

In the main text, we performed many analyses that have been applied to a range of molecular GP maps. For completeness, we include one further aspect here, the concept of ‘shape space covering’: this concept postulates that when we start with a given initial genotype and consider all genotypes within a mutational distance of at most *d* from that genotype, then most high-frequency phenotypes exist among this set of genotypes, even when the value of *d* is small [10]. Other authors use a slightly different definition of ‘shape space covering’ that is not limited to high-frequency phenotypes, but includes all phenotypes [11]. However, here we work with the original definition.

Here, we test the hypothesis of ‘shape space covering’ by following the methods in ref [10]: we start with a randomly selected genotype and evaluate the fraction of phenotypes found within at most *d* mutations from that genotype. This analysis is repeated for several initial genotypes and the results are shown in Fig S1: we find that the number of high-frequency phenotypes covered within *d* mutational steps increases rapidly with *d* and that about 50% of these frequent phenotypes are found after around *d ≈* 15 mutations, even though the maximum distance between two genotypes is 55 if each integer value had to change from the lowest permitted value (*−*3 for [*g*_1_*, …, g*_8_] and 1 for *g*_9_) to the highest permitted value (3 for [*g*_1_*, …, g*_8_] and 8 for *g*_9_). In this analysis, we have followed the definition by Grüner et al. [10] and considered a phenotype to be among the high-frequency phenotypes if its phenotype frequency is higher than the average phenotype frequency of all phenotypes.

Two reasons are given in the literature for why the ‘shape space covering’ property is found in many GP maps: first, the set of genotypes within a mutational distance of at most *d* grows rapidly with mutational distance *d* due to the many ways in which *d* mutations can be combined along the genotype (i.e. because the mutational space is high-dimensional) [12]. Secondly, high-frequency phenotypes are often so frequent that they are likely to be found among even a small set of random genotypes [13]. We can investigate the first aspect by recording the fraction of all valid biomorph genotypes that are found within *d* mutations of an initial genotype (orange line in Fig S1). We find that this fraction increases quickly in only a few mutational steps, as expected. However, we also find that for a given *d*, the fraction of high-frequency phenotypes within *d* is even higher than the fraction of all genotypes. Therefore, the second argument also applies: high-frequency phenotypes are likely to be found in a relatively small set of genotypes, simply because there are many genotypes mapping to each of these phenotypes.

### S3 Robustness of GP map properties to changes in the phenotype definition

In the methods section of the main text, we describe, how we convert each 2D biomorphs figure into a discrete phenotype for our computational analysis. This process relies on two parameters: the grid size and the threshold above which a pixel is set to one. In our analysis, we used a 30 *×* 30 grid and set a pixel to one if the total length of all unique line segments within that pixel equaled at least *≥* 20% of the width/length of the pixel. In this section, we vary these two parameters and repeat key aspects of the GP map analysis, in order to test how robust our results are to the details of the phenotype definition.

#### S3.1 Grid size

GP map data for different values of the grid size are shown in Figs S2-S3: a lower resolution of 20 *×* 20 is used in Fig S2 and a higher resolution of 40 *×* 40 is used in Fig S3. We find that the qualitative results of the analysis are unchanged: we still find phenotypic bias over several orders of magnitude, this bias is towards a subset of the simple phenotypes, phenotype robustness is correlated with the logarithm of the neutral set size, mutation probabilities (if non-zero) tend to be higher for higher-frequency phenotypes and the relationship between robustness and evolvability is negative on a genotypic level, but (weakly) positive on a phenotypic level. All these results continue to be in agreement with the simple analytic model, except the observed simplicity bias, which cannot be directly compared to the analytic model, but is instead consistent with the log-linear upper bound predicted by Dingle et al. [7].

While the qualitative trends and results are all robust to different parameter choices, the quantitative data does show differences: for example phenotypic evolvability values tend to be higher when using a more coarse-grained treatment on a 20x20 grid than when choosing a more fine-grained treatment on a 40x40 grid (Fig S2F compared to Fig S3F). This observation can be explained as follows: in a more coarse-grained treatment, more sequences belong to a given neutral set and thus neutral set sizes are higher. Such changes can have a big effect on evolvability since a single transition from *p* to a new phenotype *q* from a single phenotype in the neutral set of *p* is sufficient to raise the evolvability by one for the entire neutral set.

#### S3.2 Discretization

Similarly, the analysis was repeated for different values of the discretization threshold: this threshold *t* determines whether a pixel, which contains a number of line segments with a total length of *l*, is set to zero or one: it is set to one if *l ≥ t* and zero otherwise. Here, we repeat the GP map analysis with values of 10% of the pixel size (Fig S4) and 50% of the pixel size (Fig S5). Again, we find that the qualitative GP map characteristics, as well as the agreement with the analytic model and the predictions from [7], are unaffected.

**Figure S2:**
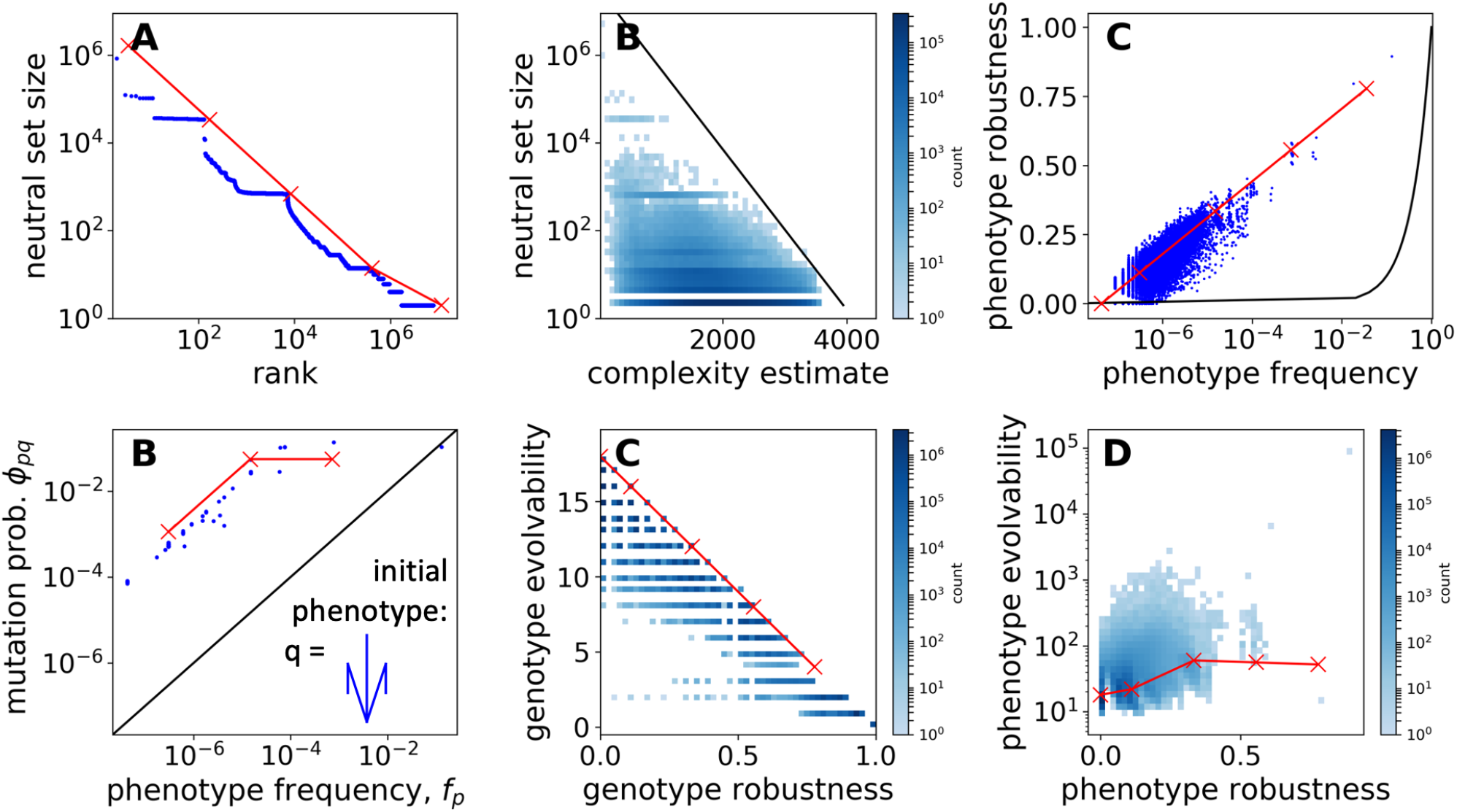
GP map analysis with a different parameter choice in the coarse-grained phenotype definition: Here, a 20 × 20 grid is used (instead of 30 × 30). The analysis shows the GP map data (blue) as well as the predictions from the analytic model (red) for the following quantities: **(A) Neutral set size vs frequency rank. (B) Neutral set size vs estimated complexity.** The black line indicates an approximate log-lin upper bound to guide the eye, as predicted in [7]. **(C) Phenotype robustness vs phenotype frequency** f_q_. The black line (ρ_q_ = f_q_) shows what we would expect in the null model from refs [8, 14]. **(D) Phenotype mutation probability** φ_pq_ **vs. phenotype frequency** f_p_ **for one specific initial phenotype** q. The black line (φ_pq_= f_p_) shows what we would expect in the null model from refs [8, 14]. Data points with φ_pq_ = 0 are excluded on this log-scale. **(E) Genotype evolvability vs genotype robustness. (F) Phenotype evolvability vs phenotype robustness.** Only the computational data depends on the grid size; the analytic data is included just for reference.

**Figure S3:**
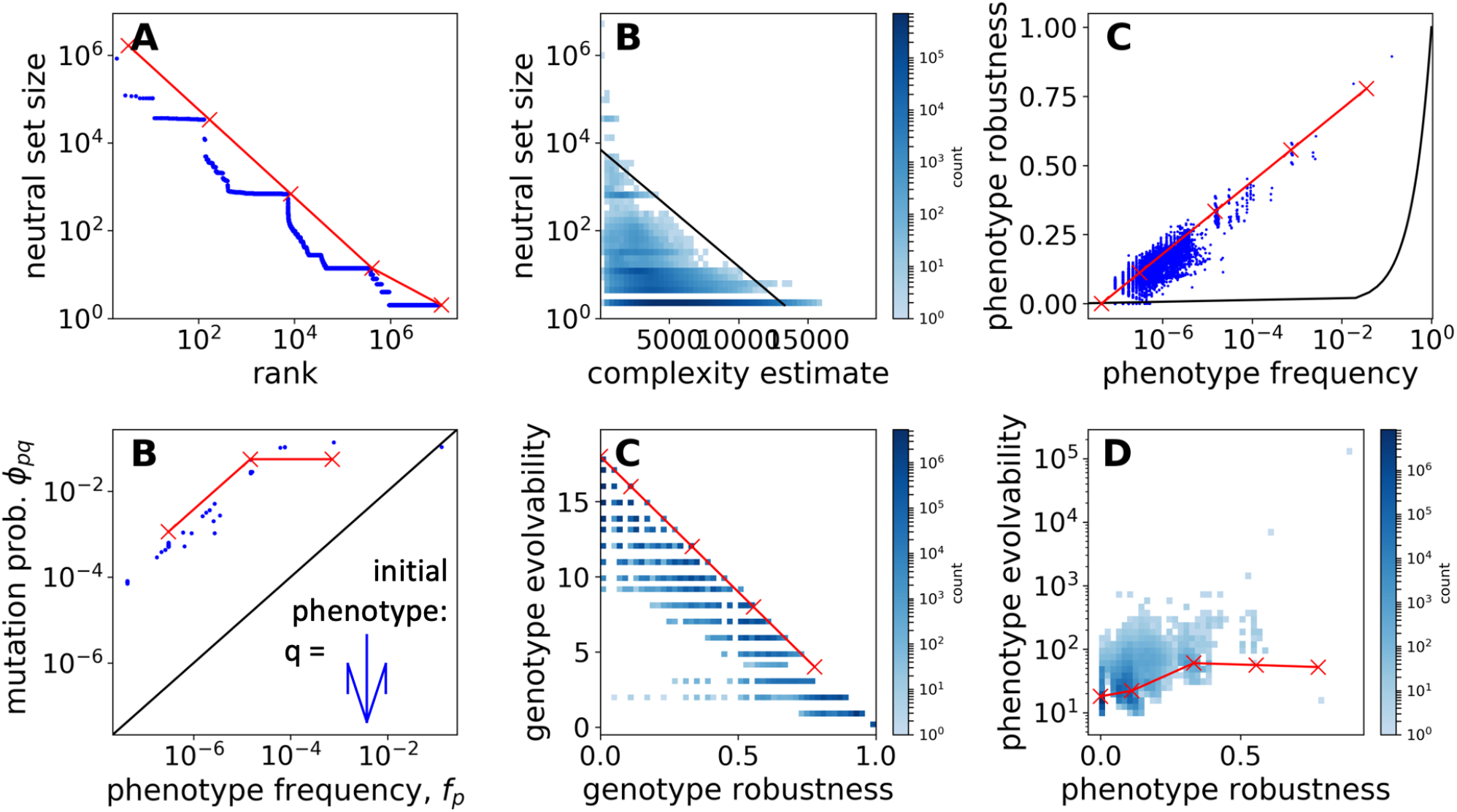
GP map analysis with a different parameter choice in the coarse-grained phenotype definition: same as Fig S2, but here a 40 × 40 grid is used.

**Figure S4:**
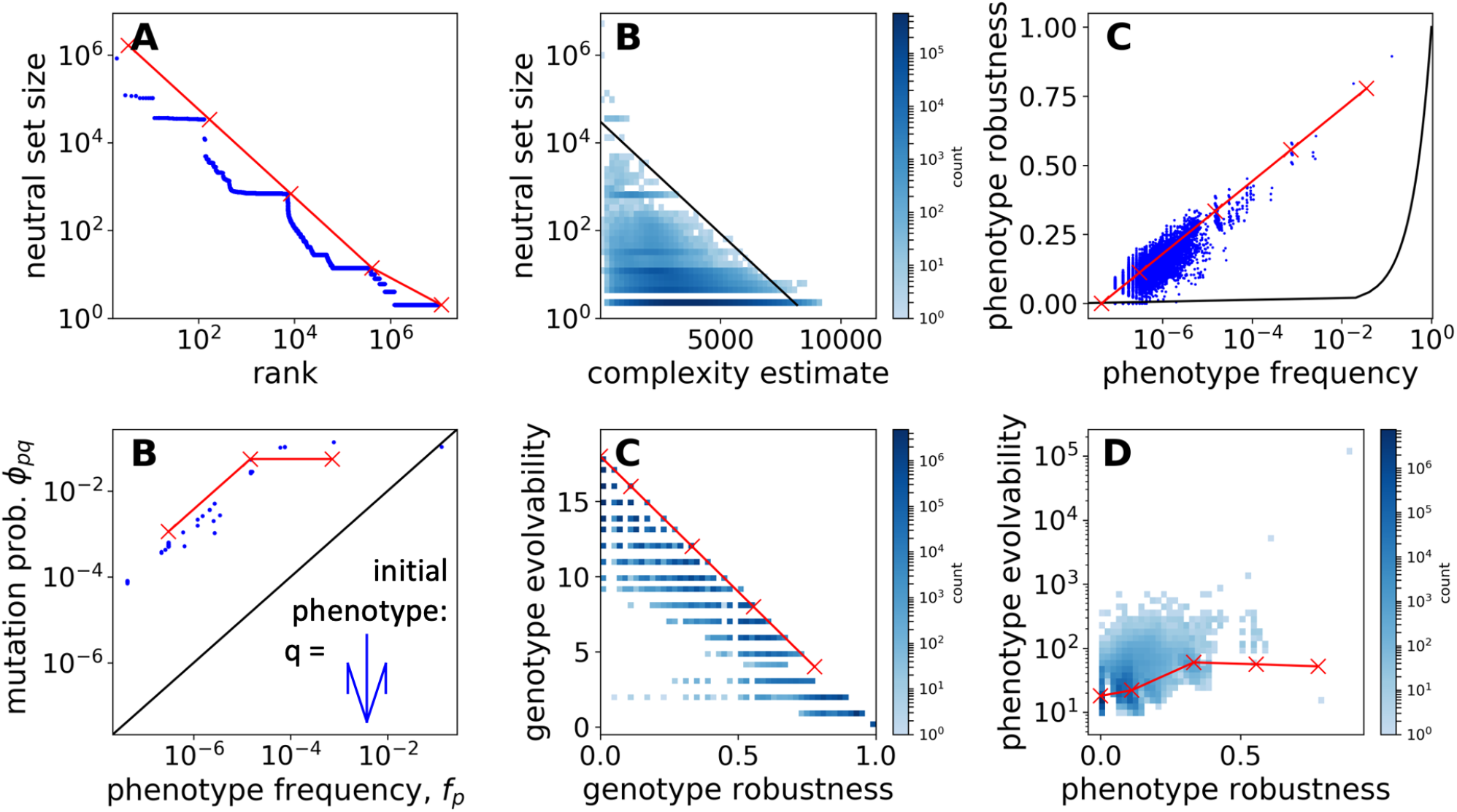
GP map analysis with a different parameter choice in the coarse-grained phenotype definition: same as Fig S2, but here the threshold is different from the one in the main text: 10% instead of 20%. The grid size, 30 × 30, is the same as in the main text.

**Figure S5:**
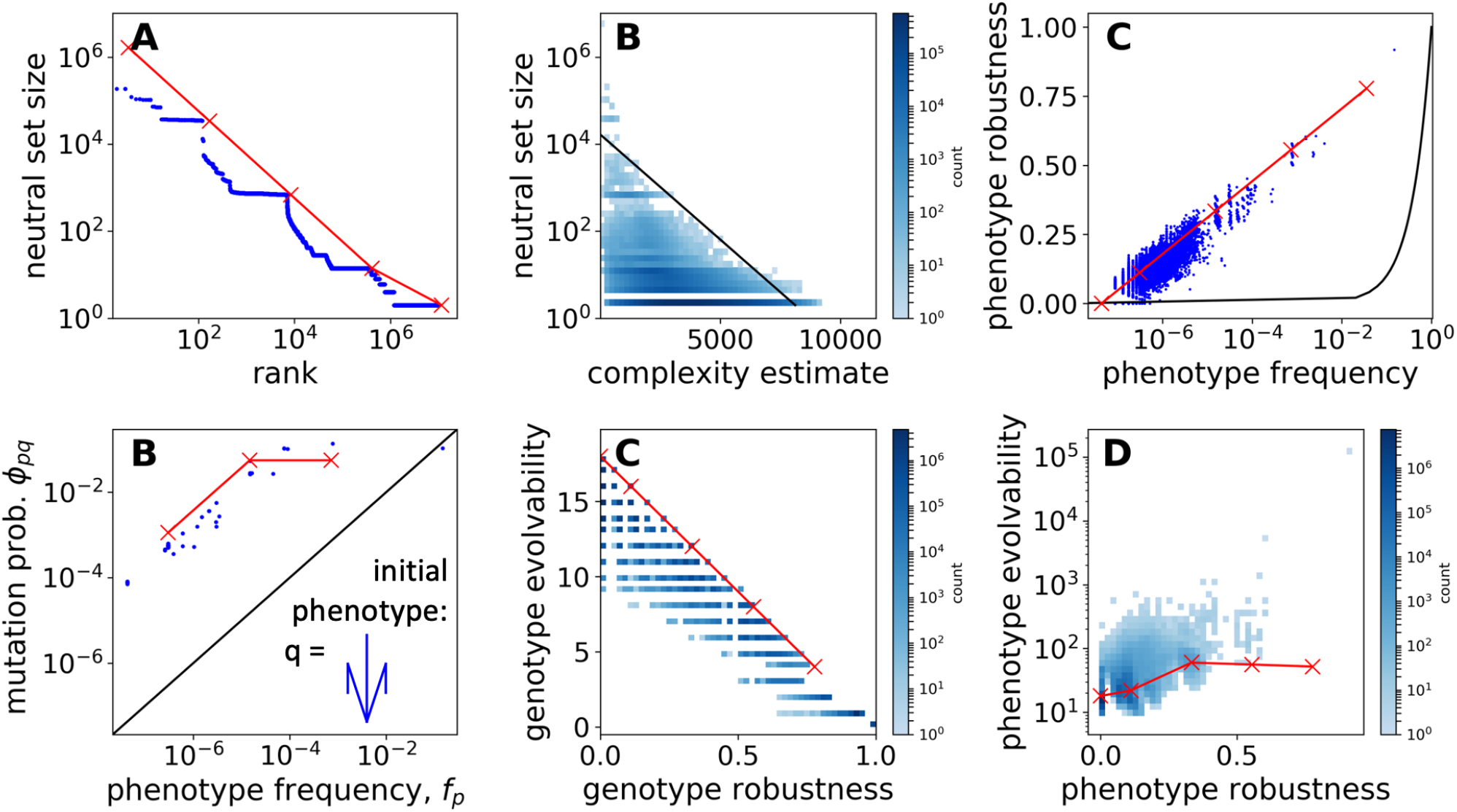
GP map analysis with a different parameter choice in the coarse-grained phenotype definition: same as Fig S2, but here the threshold is different from the one in the main text: 50% instead of 20%. The grid size, 30 × 30, is the same as in the main text.

**Figure S6:**
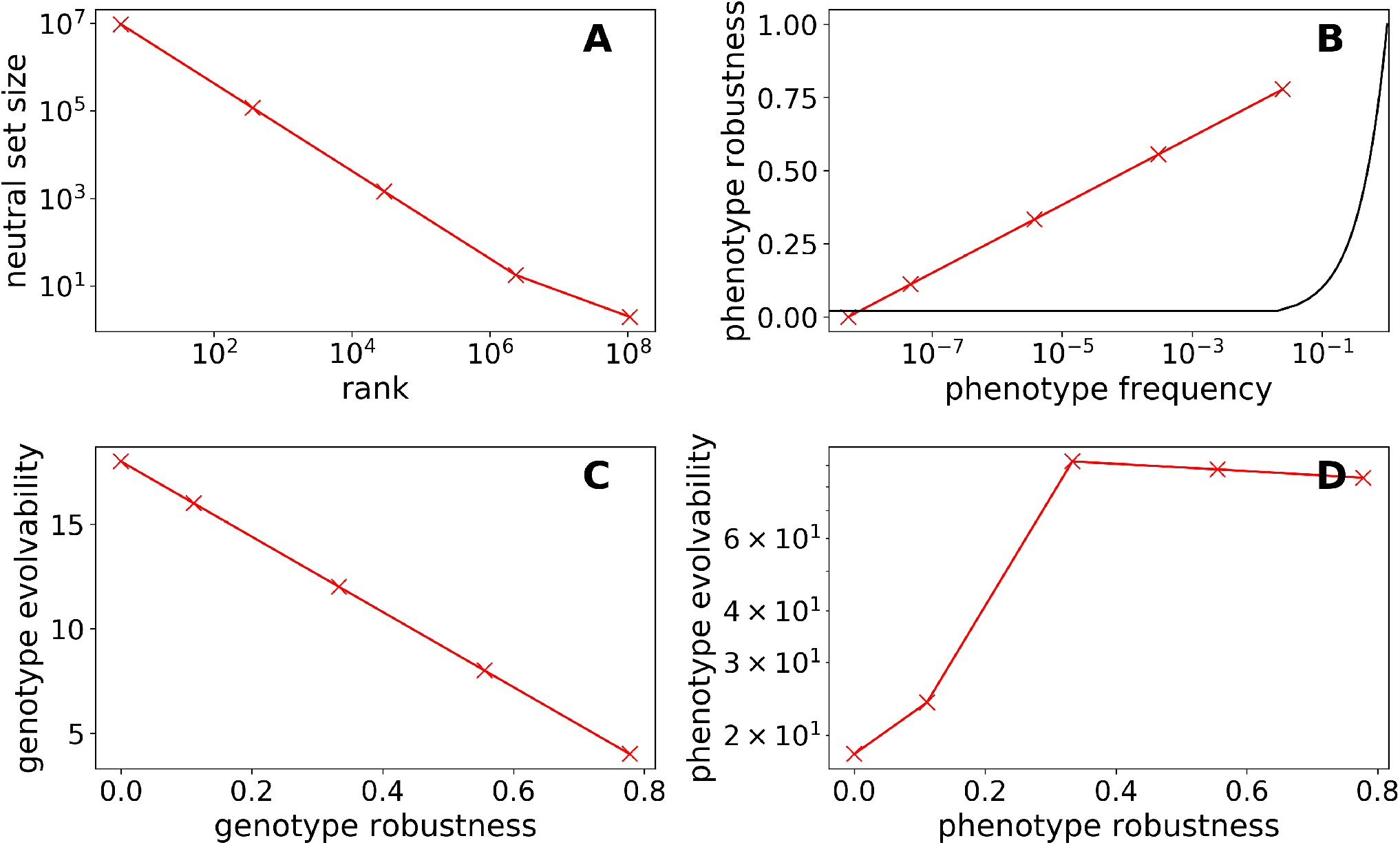
Predictions from the simplified analytic model for a larger range of permitted values at each genotype position: here, a larger range of values is permitted at each position of the genotypes than in the main text: nine values for each of the ‘vector genes’ (−4 ≤ g_i_ ≤ 4 for i ∈ [1, .., 8]) and nine values for the ninth gene (1 ≤ g_9_ ≤ 9). With these parameters, there are 9^9^ ≈ 4 × 10^8^ genotypes and so a full computational analysis is no longer feasible. However, approximate predictions from the analytic model can be made and these are shown in this figure. The plots show: A) Neutral set size vs. frequency rank (Eq 4). B) Phenotype robustness vs. phenotype frequency (Eq 10). C) Genotype evolvability vs. genotype robustness (Eq 11). D) Phenotype evolvability vs. phenotype robustness (Eq 13).

With our analytic model, we can allow an arbitrary range of values in the genotypes, without computational difficulties. Here we consider a GP map, where the genotypes can take on a wider range of values: nine values for each of the ‘vector genes’ (*−*4 *≤ g_i_ ≤* 4 for *i ∈* [1*, ..,* 8]) and nine values for the ninth gene (1 *≤ g*_9_ *≤* 9).

The data in Fig S6 indicate that permitting a wider range of integers in the genotype would not affect our qualitative results.

#### S4.1 Examples of phenotypes and their estimated complexities

In order to visualize the data in Fig 3C of the main text, we focus on phenotypes with different combinations of complexity values and neutral set sizes: Fig S7 shows examples of simple phenotypes with low neutral set sizes, simple phenotypes with high neutral set sizes and complex phenotypes with low neutral set sizes (complex phenotypes with high neutral set sizes do not exist since we have observed simplicity bias in this GP map). While the analytic model cannot be compared directly to this data, which is based on the computational phenotype treatment, we can still use insights from the analytic model to guide our interpretation: we find that, as we might expect from the analytic model, phenotypes with few lines (and hence low *g*_9_ and high neutral set sizes) are simple. Phenotypes with many lines (and hence high *g*_9_ and low neutral set sizes) can have high complexity, but they can also have low complexity in the coarse-grained computational model, for example, if each vector is used exactly once or if some vectors overlap.

#### S4.2 Alternative method of estimating phenotypic complexity

Since Kolmogorov complexity cannot be measured exactly, just estimated, we repeated the analysis in the main text for a different method of estimating complexity: we use a compression-based method, the Lempel-Ziv approach [15], relying on the implementation from ref [7]. This implementation takes a binary string as an input, but our phenotypes are 2D binary grids. Therefore, we concatenated the rows of our grid before passing it to the complexity estimator. Since Dingle et al. [7] argue that the complexity of a string is best estimated as the mean of the estimated complexity of the string and the estimated complexity of its reverse, we performed an analogous calculation on our 2D array: we took the transpose of the binary grid and included it in the estimate. This means that we did not only take a mean of the estimated complexity of the string and its reverse, but also the corresponding concatenated string of the transposed array and its reverse.

The results from this compression-based complexity estimator are shown in Fig S8: we still find that there are no complex phenotypes with high neutral set sizes. As before, the data approximately falls below a log-linear line, derived theoretically in ref [7].

#### S4.3 Alternative method of estimating phenotypic complexity in the analytic model

In the analytic model, we have a clear criterion for when two genotypes fall into the same neutral sets, formulated in terms of constrained and unconstrained sites, but we do not have a visual description of each corresponding phenotype. Thus, we estimated phenotype complexities using the information that needs to be encoded in the genotype (section S1.3). However, there is one way of approximating the visual complexity of each phenotype: if we simply assume that each line in a phenotype drawing takes the same amount of information to encode, then the number of lines in the biomorphs figure is a good proxy for the total description length, i.e. the complexity. This is only an approximation since a set of parallel lines can be encoded more efficiently than a set of arbitrary lines (in the same way that repeating strings can be compressed, but arbitrary strings cannot). However, if we use the number of lines *n_l_* that are drawn in the biomorphs image, whether they are overlapping or not, as a first approximation, we get:

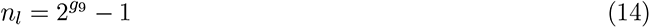

Since we also have an expression for the neutral set size as a function of *g*_9_ (eq 2), we can use this parametric relationship to obtain the data in Fig S9. We find that the qualitative trend is the same: more complex phenotypes with a higher number of lines tend to have lower neutral set sizes. However, this relationship no longer follows the log-linear relationship predicted in ref [7]. This is because the number of lines in the biomorphs figure can exceed the number of distinct vectors (which is eight). Then the same vectors appear with different scaling factors multiple times in the figure and so the number of lines increases more rapidly than the amount of genotypic information needed to encode them.

#### S4.4 Distribution of phenotypic complexities for arbitrary genotypes

In the main text, we found that a phenotype with a large neutral set is likely to be simple. Thus, a given simple phenotype is more likely to have a high phenotype frequency and appear in a small random sample of genotypes than a given complex phenotype. However, there may be many different simple phenotypes and many different complex phenotypes, so it is not clear, how many simple phenotypes we expect to find overall in a random sample of genotypes. Arguments in the SI of ref [17] imply that while the neutral set size of an individual complex phenotype is small, the number of distinct complex phenotypes is much larger than the number of distinct simple phenotypes, and so overall, the likelihood of drawing any complex phenotype from a random sample of genotypes is approximately equal to the likelihood of drawing any simple phenotype.

Here we test both parts of this argument for the biomorphs GP map, using our computational results. First, we consider the number of simple and complex phenotypes (Fig S10): we find that there is only a small number of simple phenotypes, as expected from the information-theoretic arguments in ref [17] . However, the number of very complex phenotypes is also limited. This deviation from the information-theoretic arguments in ref [17] is likely due to the constraints of the biomorphs system, which only permits certain geometric forms (for example no phenotype can have two disconnected parts and this fact alone severely restricts the number of possible 2D drawings). Secondly, we calculate the fraction of *genotypes* with simple or complex phenotypes (Fig S11). We find that there is a range of complexity values, for which the probability of finding a phenotype with that complexity is approximately constant, in agreement with the information-theoretic arguments in ref [17]. However, at high complexities, we see a deviation from the information-theoretic expectation, which is consistent with our results from Fig S10.

So far, complexity distributions like this have not been discussed in much detail (exceptions are in the SI of ref [17], and for one matrix-rewriting grammar GP map in ref [18] and digital logic gates in ref [19]) and so future work should investigate these distributions and their implications more thoroughly, both for the biomorphs and for other GP maps.

**Figure S7:**
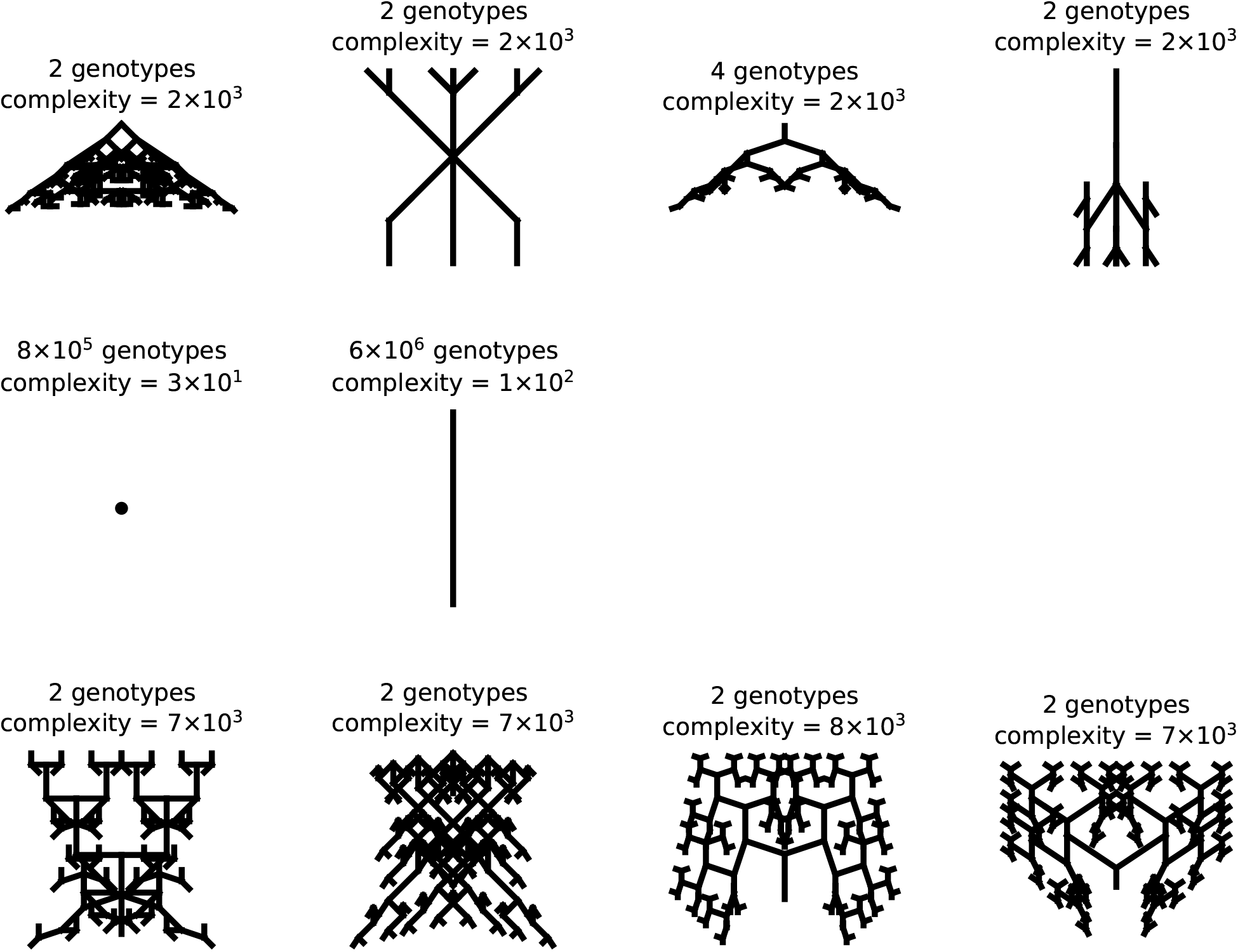
Neutral set sizes and complexity estimates for a few example phenotypes: the figures show examples of rare and simple phenotypes (top row), of frequent and simple phenotypes (middle row), and of rare and complex phenotypes (third row). Note that the labels ‘rare’/‘frequent’ and ‘complex’/‘simple’ are discrete categories that represent a range of values - not all of the ‘simple’ phenotypes have the same complexity in this figure.

**Figure S8:**
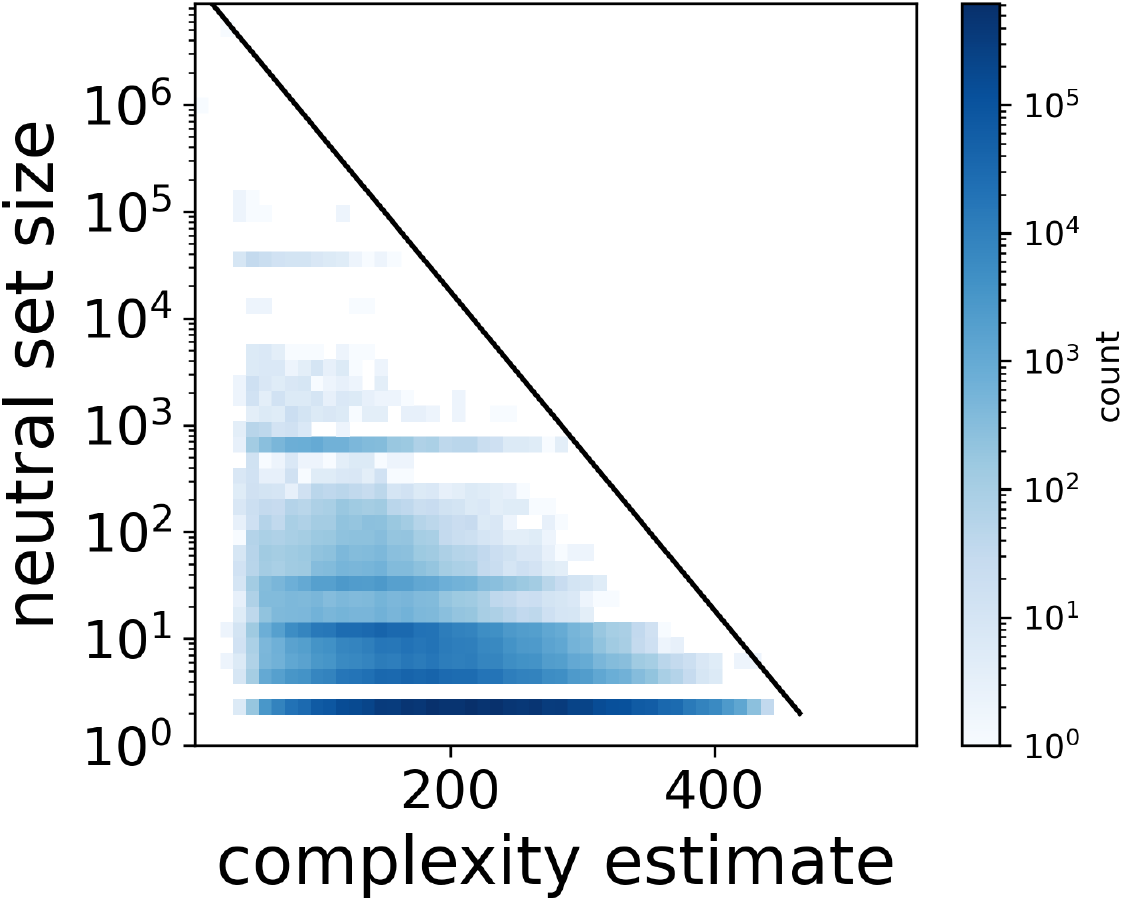
Neutral set size vs complexity estimate with a compression-based complexity estimator: as in Fig 3C in the main text, we plot the neutral set size of each phenotype (on a log scale) against an estimate of its complexity. Here, this complexity is computed by feeding a concatenated version of each phenotype’s binary pixel array into the Lempel-Ziv implementation from ref [7]. A log-linear line, as predicted as an upper bound [7], is drawn to guide the eye.

**Figure S9:**
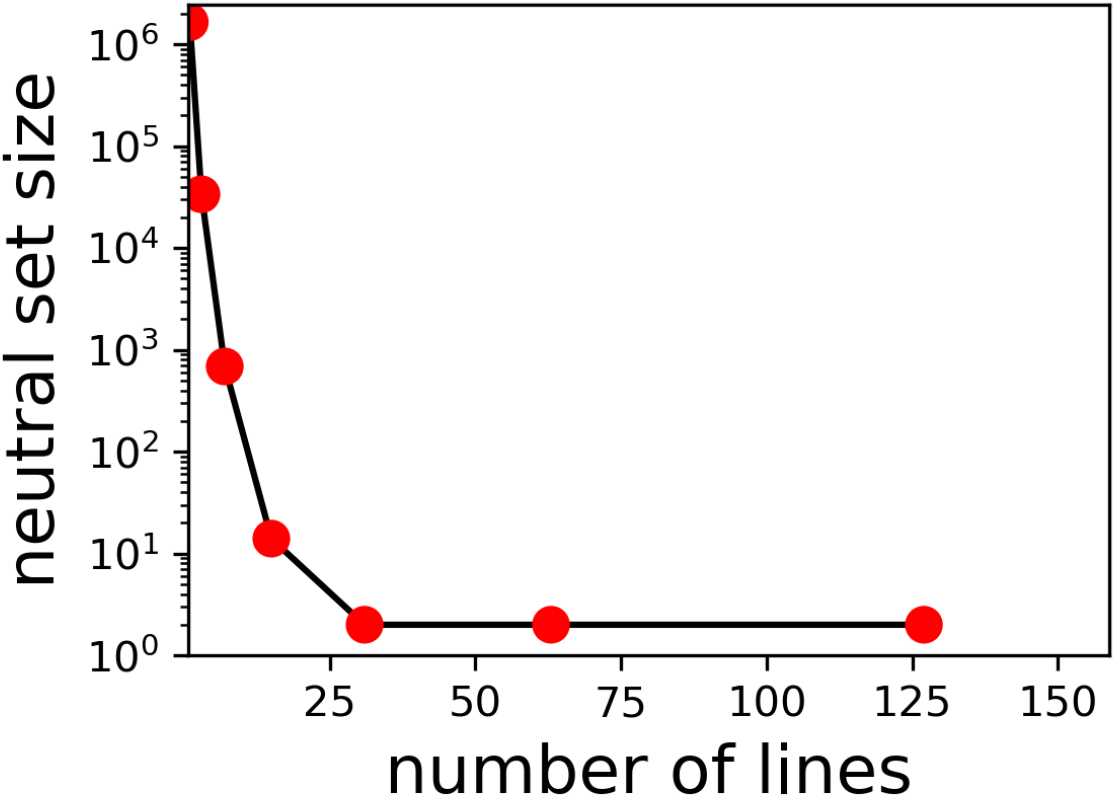
Neutral set size vs number of lines in the figure: both quantities are estimated with the analytic model, using eq 2 and eq 14.

**Figure S10:**
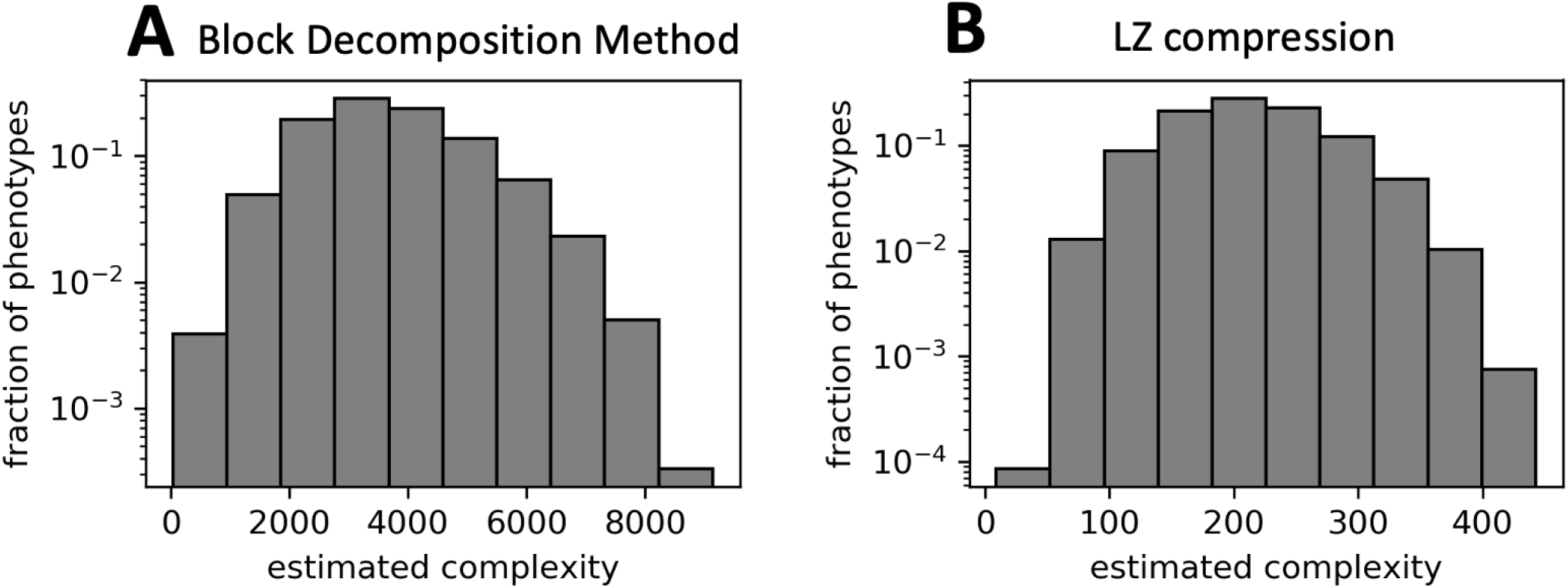
Probability P_p_(K) of obtaining a phenotype of complexity K upon random sampling of phenotypes. Most of the ≈ 10^7^ phenotypes have intermediate complexity values - phenotypes with very low or very high complexities are rare. A) Phenotypic complexities based on the Block Decomposition Method [16] (as in the main text). B) Phenotypic complexities based on Lempel-Ziv compression (as in section S4.2 above).

**Figure S11:**
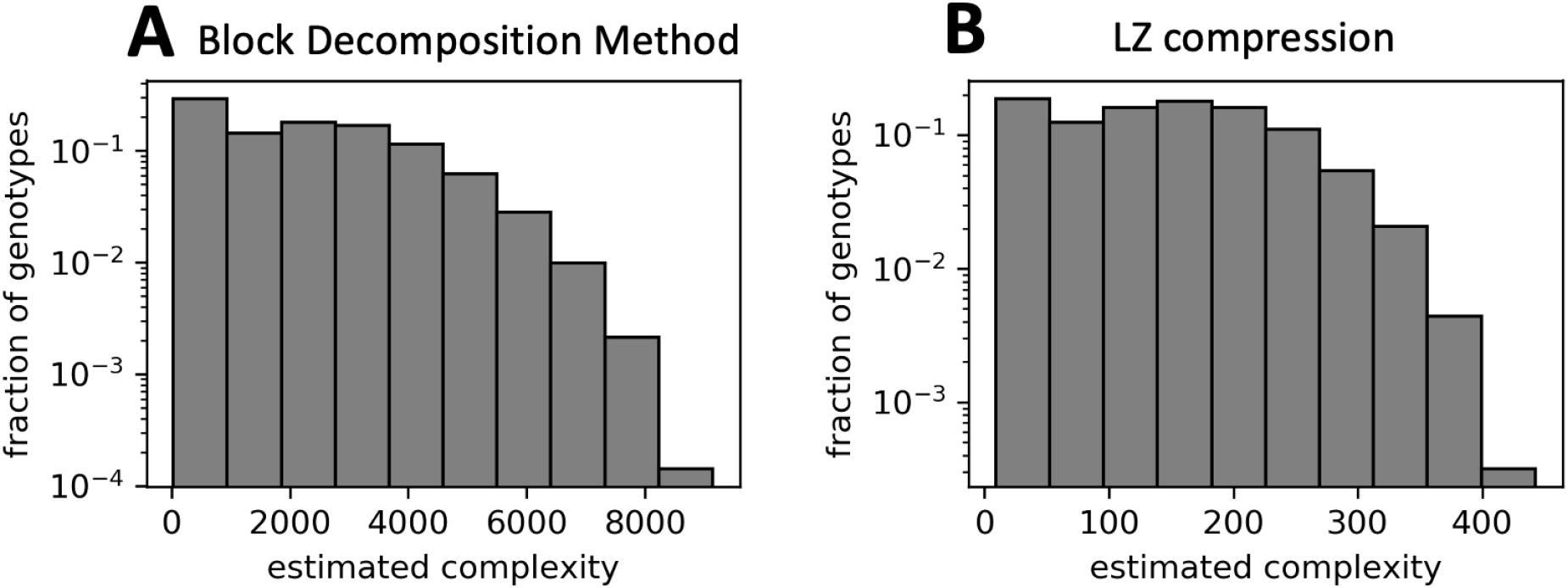
Probability P_g_(K) of obtaining a phenotype of complexity K upon random sampling of genotypes. A) Phenotypic complexities based on the Block Decomposition Method [16] (as in the main text). B) Phenotypic complexities based on Lempel-Ziv compression (as in section S4.2 above).

**Figure S12:**
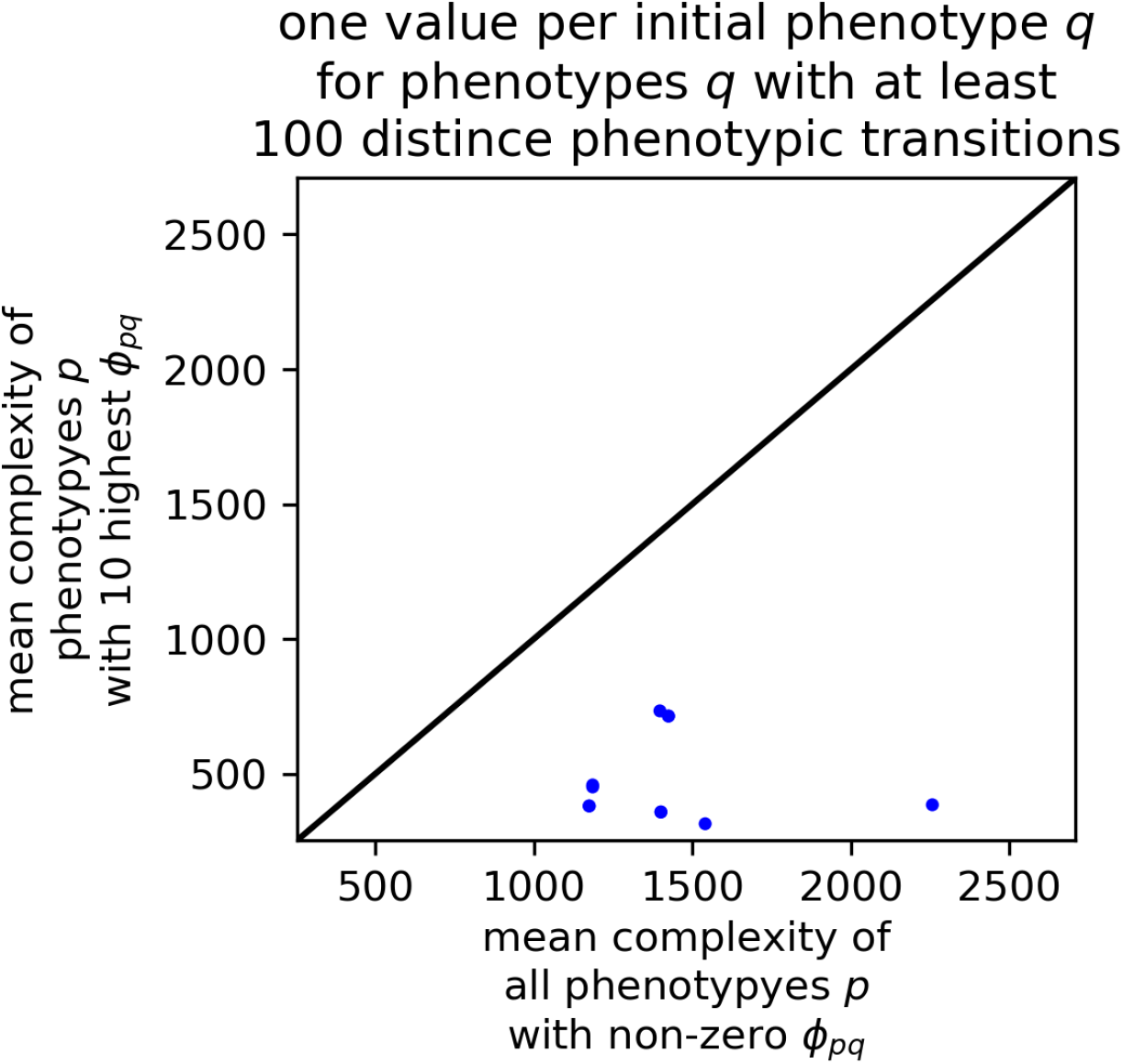
Mutational bias towards simple phenotypes: we consider all phenotypes q, from which at least 100 different phenotypes p can be reached through mutations (i.e. E_q_ ≥ 100). For each initial phenotype q, we plot the mean complexity of the 10 phenotypes p with the highest φ_pq_ values against the mean complexity of all phenotypes with non-zero φ_pq_ values, in order to compare likely phenotypic transitions to the full set of possible phenotypic transitions. The black line indicates equality (x = y). We find that for all initial phenotypes q, the complexity of the high-φ_pq_ phenotypes is lower, indicating that mutation probabilities to simple phenotypes q tend to be higher. The data in this plot is for our computational approach based on the coarse-grained images.

#### S4.5 Simplicity bias in mutation probabilities

In the main text, we argued that the strong simplicity bias found in the biomorphs GP map means that a random mutation on a random genotype is more likely to give a simple phenotype than a complex one. Here, we test whether this continues to hold when we consider mutations for a fixed initial phenotype (i.e. whether there is simplicity bias in the *φ_pq_* values). The data is shown in Fig S12: for a fixed initial phenotype *q*, we compare the complexity of the ten phenotypes which are most likely to appear after mutations (i.e. with the highest *φ_pq_* values), to the complexity of all phenotypes which can appear after mutations. This data indicates that, regardless of the initial phenotype *q*, phenotypic changes that happen with a high probability through mutations tend to be towards simpler phenotypes than phenotypic changes that occur with a lower probability. A caveat of this analysis is that we limit the set of initial phenotypes *q* to phenotypes, from which at least 100 different phenotypic changes are possible through mutations since the top-ten values are only relevant if there are *»* 10 non-zero *φ_pq_* values.

### S5 Phenotypic Bias when the number of developmental stages is kept constant

**Figure S13:**
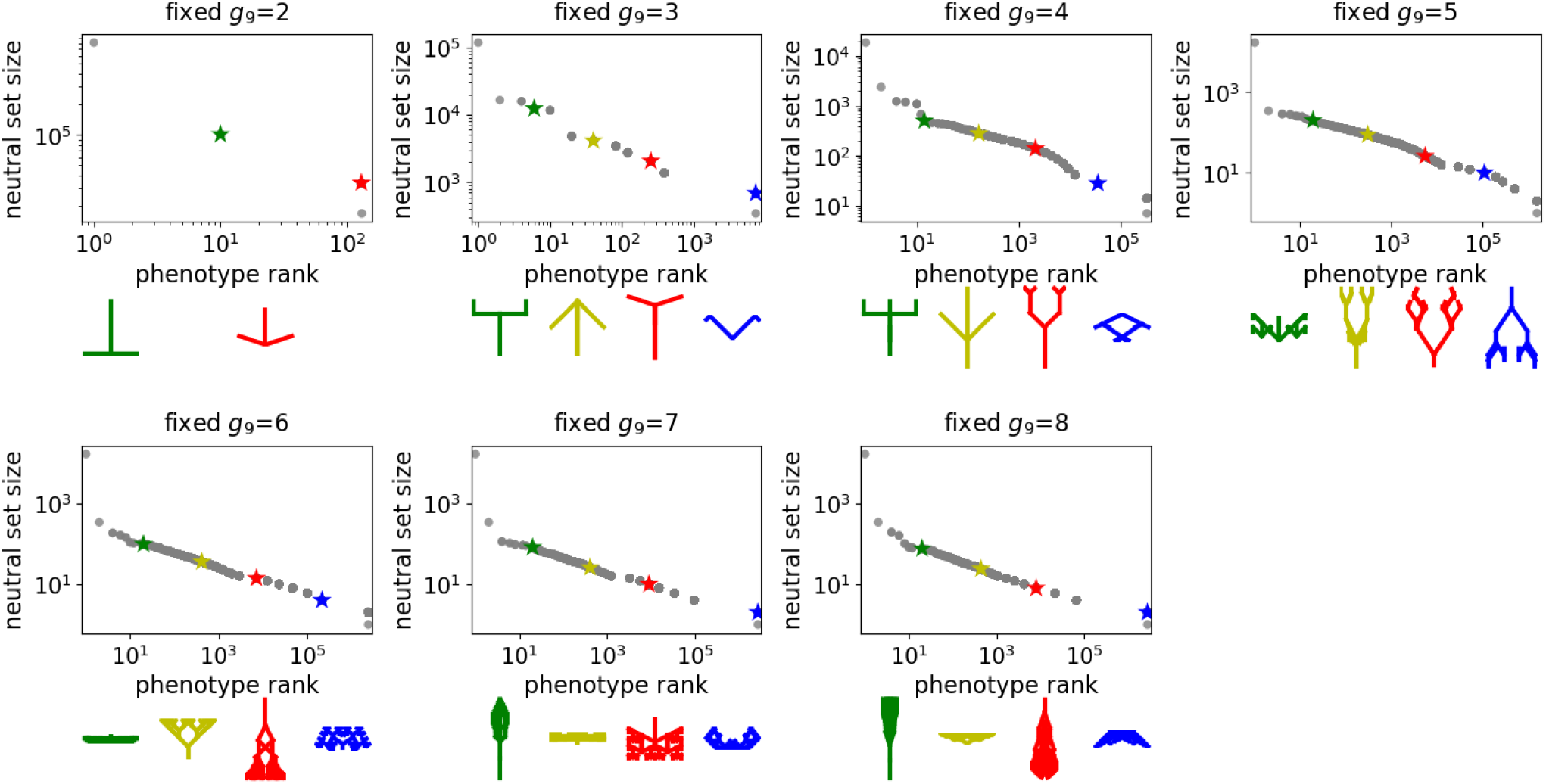
Neutral set size vs rank for fixed values of g_9_: each plot analyses the slice of the GP map that is defined by a fixed value of g_9_between g_9_ = 2 and g_9_ = 8 (indicated in the plot titles). In each case, the number of genotypes per phenotype (i.e. the neutral set size at fixed g_9_) is computed for all phenotypes present in the given slice and plotted as a rank plot using our computational approach with the same parameters as in the main text. To illustrate, what kind of phenotypes are frequent/rare in each case, a few phenotypes are highlighted in each plot and drawn in corresponding colors underneath the plot.

In the analytic model, all genotypes that map to a given phenotype have the same value of *g*_9_, the final site of the genotype. This value then determines the neutral set size of the given phenotype. If the analytic model was perfect, this would mean that there would be no phenotypic bias at all if *g*_9_ was kept fixed. Fig S13 shows that this is not the case and that there is bias even at constant *g*_9_, albeit over fewer orders of magnitude. The reason for this is the following: the analytic model assumes that changing any vector component of any vector in the figure will change the phenotype. However, if all x-components of all vectors are set to zero, then the choice of y-components does not matter since the phenotype will be a single vertical line along the y-direction for any choice of y-components. Thus, this is a simple example, where the analytic model would underestimate the neutral set size by a large factor. We can therefore conclude that the analytic model is only an approximation and that neutral set size differences in the biomorph model are more subtle than predicted by the analytic model.

### S6 Identifying an evolutionary path with the smallest number of phenotypic changes

In the main text, we illustrated how a line-shaped initial phenotype can be transformed into a specific insect- shaped phenotype in single point mutations, such that the number of phenotypic changes during the process is as small as possible. Here, we describe the computational approach we used to obtain this shortest series of phenotypic changes.

It is useful to reframe this optimization problem in terms of neutral components (NCs): a NC is a subset of the neutral set of *p*, which is defined [20] such that two genotypes are in the same NC if they can be connected by a series of neutral mutations. Therefore, it is possible to get from any genotype in a NC to any other genotype in the same NC during a period of neutral evolution, but any transition to another NC through mutations has to include phenotypic changes. NCs for a given GP map can be identified efficiently, for example by building on methods from ref [10].

Once we have enumerated all NCs, identifying the path with the smallest number of phenotypic changes is equivalent to finding the sequence of mutations with the smallest number of mutations that change the neutral component (NC) since two NCs that are connected by point mutations always correspond to different phenotypes, by definition. This allows us to solve our optimization problem by focusing on NCs and not individual genotypes. Thus, we created a unique ID for each NC in the GP map and evaluated, which NCs can be reached from a given NC through single point mutations. This defines a network, where each NC is a node and each edge indicates that there are point mutations connecting two NCs. Then we found the shortest path in this network using Dijkstra [21]’s algorithm. The resulting list of NCs gives the shortest list of NC transitions that have to be made in order to convert the initial phenotype into the target phenotype through point mutations. Then we simply mapped the NCs to the corresponding phenotypes to obtain the final figure.

### S7 Two-peaked landscapes - beyond the first fixation

In the main text, we studied a scenario based on refs [8, 22], where a population with an initial phenotype *p*_0_ evolved adaptively to one of two fitness peaks, *p*_1_ or *p*_2_. We reported which of these two phenotypes was the first to go into fixation, the mutationally more accessible phenotype *p*_1_ or the phenotype with higher selective advantage *p*_2_. Here, we investigate what happens after the first fixation event. Since there are no point mutations that can convert *p*_1_ into *p*_2_ directly (or vice versa), the only possible phenotypic changes are either back to the initial phenotype *p*_0_, which is less fit, or directly to the other phenotype through a rare event involving multiple mutations at once. To investigate, if one of these phenotypic changes is likely to happen, we ran additional simulations that spanned 10^6^ generations each and were not terminated with the first fixation event. This is about ten times longer than the typical number of generations until the first fixation event, which is between *≈* 10^5^ and *≈* 10^5^ (depending on the parameters *s*_1_ and *s*_2_). We ran 100 such simulations for each combination of parameters *s*_1_ and *s*_2_. We found no instances of direct fixations from *p*_1_ to *p*_2_, implying that the requirement for multiple specific mutations to coincide makes such transitions extremely unlikely. Thus, the only new fixations that we observe after the first fixation event is a reversion to *p*_0_, the phenotype with the lowest fitness. This only happens in rare cases (*<* 0.5% of cases) and only if the phenotype that fixes first only has a low selective advantage over *p*_0_ (Fig. S14).

**Figure S14:**
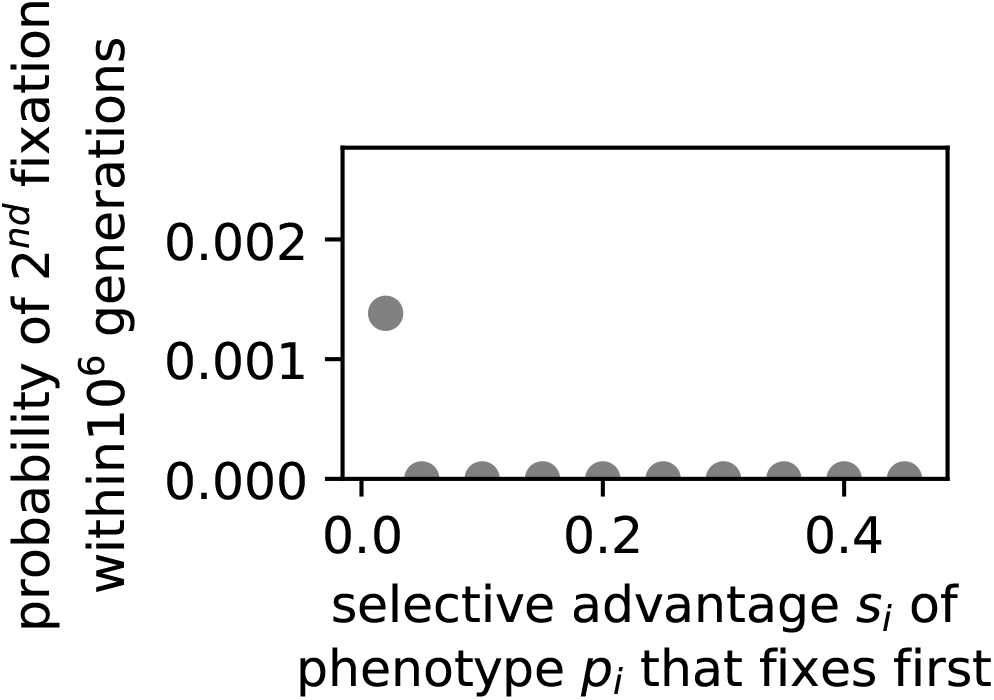
Likelihood of reversion to fitness valley in the two-peaked landscape: We simulate evolving populations in the two-peaked landscape described in the main text for 10^6^ generations. The first fixation of one of the local maxima (p_1_ or p_2_) typically occurs after 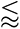 10^5^ generations. Here we investigate how likely the population is to experience a renewed fixation to the initial phenotype p_0_ (the fitness valley). We plot this likelihood against the selective advantage s_i_ of the phenotype that was the first to fix. As might be expected, reversions to the fitness valley via drift are rare since we have strong selection with Ns_i_ ≥ 10. The same parameters are used as in the main text, but the data is only based on 100 repetitions for each combination of the parameters s_1_ and s_2_ because of the higher computational cost of this analysis.

### S8 Selection for tree-like shapes

Here, we repeat the evolutionary simulation from Fig 5 in the main text, but with a slightly more complex fitness landscape inspired by one of the scenarios in Johnston et al. [17]: we assume that all tree-like phenotypes are equally fit (fitness *F* = 1) and all other phenotypes are completely unviable (*F* = 0). To obtain a reproducible and simple definition of what a tree-like shape is, we proceed as follows: we crop empty margins from the biomorph’s grid representation and then consider the resulting cropped grid to be tree-like if there is a ‘stem’ in the lower part of the figure (i.e. the pixels on the lower fifth of the y-axis are filled in) and this stem is surrounded by some space (i.e. the pixels immediately to the right and left of the stem are not filled in). We also consider a single vertical line to be a tree-like phenotype.

The data is shown in Fig S15: we find phenotypic bias over several orders of magnitude even among the more restricted set of tree-like phenotypes. This phenotypic bias is reflected in the evolutionary simulation: while selection confines the population to tree-like shapes, the phenotypic bias still plays an important role in determining *which* of the tree-like shapes appear more often in the evolving population. Here, some of the frequent shapes are actually observed more often than expected based on their phenotypic frequencies, which is likely due to selection for high robustness at this high mutation rate of *µ* = 0.1 (the ‘survival of the flattest’ [23] effect).

**Figure S15:**
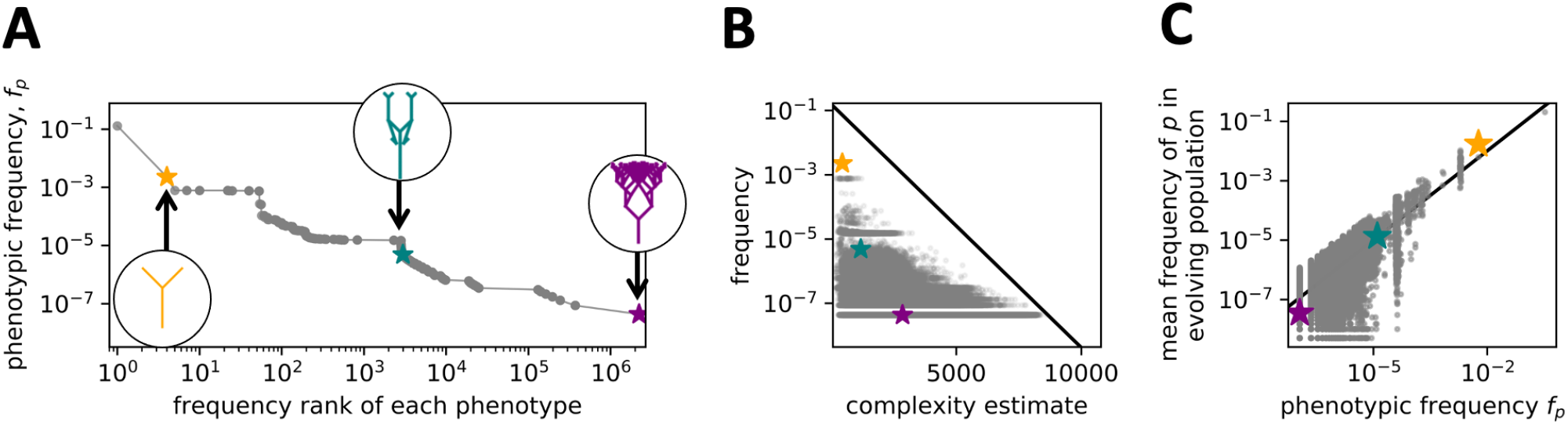
Phenotypic bias towards simple shapes if the analysis is restricted to tree-like biomorphs: The concepts of this plot are the same as in Fig 5 in the main text, but here only tree-like phenotypes have non-zero fitness in the evolutionary simulation: (A) Phenotype frequency vs rank for all tree-like phenotypes. Three phenotypes are selected from this plot: one with high frequency (yellow), one with medium frequency (teal), and one with low frequency (purple). We find bias even among tree-like phenotypes. (B) Phenotype frequency vs estimated complexity for all tree-like phenotypes, with the three selected phenotypes from (A) highlighted in color. We observe simplicity bias among the tree-like shapes. (C) As a simplified model of an evolutionary process, we assume that all tree-like phenotypes are equally fit (fitness F = 1) and all other phenotypes are completely unviable (F = 0). We model a population of 2000 individuals with a mutation rate of µ = 0.1 for 10^5^ generations. In this process, we record, how frequently we each of the selected shapes from (A) occurs. In (C), the normalized number of times each phenotype occurs in the population is plotted against a renormalized version of its phenotypic frequency (such that the frequencies of all tree-like shapes sum to one). We find that the phenotypic bias among the tree-like shape is reflected in their frequency in the evolving population.

The strong bias in the frequency of different tree-like shapes in the simulation is not surprising since transitions between different tree-like biomorphs are neutral in this scenario. However, it constitutes a simple example, where there is selection on some features of the phenotype only, which may be a realistic approximation under some conditions.

### S9 Historical background to the discussion on ultimate and proximate causation in development

#### S9.1 Historical theme 1: Developmental constraints or biases

We begin by exploring the language of constraints. In a highly influential 1985 paper, John Maynard-Smith et al. [24] opened with the definition: “*A developmental constraint is a bias on the production of variant phenotypes or a limitation on phenotypic variability caused by the structure, character, composition, or dynamics of the developmental system.”*. They distinguished between “universal” constraints which hold for all organisms, and “local constraints” which hold for a limited range of species or taxa, but were careful to emphasize that this is not a sharp dichotomy.

Universal physical constraints on phenotypes are everywhere in the natural world, setting limits on viable phenotypic variation. They can arise from the basic laws of mathematics, physics, and chemistry or from principles of self-organization, as emphasised by Kauffman [25]. Universal constraints are not contingent, in the sense that if one were to re-run the tape of life again, they would still hold in much the same form. Other universal constraints may have an element of contingency: one example given by Maynard-Smith et al. [24] is that the mapping from codons to amino acids in the genetic code could have been different [26, 27] but once it is fixed, it is unlikely to change again.

It is also not controversial that local developmental constraints are active everywhere in the natural world, shaping the spectrum of variation upon which natural selection can act. This point is made repeatedly in *The Blind Watchmaker* [1]. For example, in the addendum on evolvability, Dawkins argues that: “*You can’t make an elephant by mutation if the existing embryology is octopus embryology*”. He further suggests that such constraints can facilitate more rapid evolution in certain, possibly favorable directions [28]. These constraints are local because they only affect certain taxa. Moreover, the developmental constraints may themselves change due to natural selection, a process Dawkins coined as the “*evolution of evolvability*” [1, 2]. He illustrates this concept with the axial symmetry of the biomorphs, a constraint that can only be overcome by modifications to the developmental program itself. Dawkins suggests that such changes may be related to major transitions in evolution.

The language around local developmental constraints has changed since the iconic paper by Maynard-Smith et al. [24], not least due to the rise of the field of evo-devo [29–33]. There is a modern preference [34, 35] for the term “*developmental bias*” over developmental constraint, in part because constraints are seen as too binary: something can or cannot happen because it is or is not constrained, while many developmental processes are more subtle in their influence [34]. In addition, development may facilitate the generation of new variation rather than only limiting it [35]. The broad spectrum of effects captured by the terms *developmental bias* and *developmental constraint*, the subtle differences in their meanings, as well as variations in how they are used by different research communities all contribute to the historical complexity of this debate [36]. The idea that hard binary constraints –such as those imposed by octopus or elephant developmental programmes –affect adaptive evolutionary outcomes is much less controversial than the claim that potentially weaker developmental biases do (see e.g. [37, 38] for some historical examples of such arguments). That said, there are further reasons for differing attitudes to biases and/or constraints that relate to background issues to be discussed in the next subsection.

#### S9.2 Historical theme 2: Mutation and selection as opposing pressures

The second theme that has long influenced the debate between internalism and structuralism arises from historical population genetic arguments which state that the “pressure” from mutation is too weak to overcome selective effects (see [38] for an overview of this history): Statements of this type are often attributed to the founders of the modern synthesis including influential work by Haldane [39] and Wright [40]. The mutation pressure argument has historically been used to counter structuralist claims that biases in the arrival of variation affect *adaptive* evolutionary outcomes [38]. By contrast, it is not controversial at all that mutational biases can play an important role in neutral evolution [37, 40, 41].

The prevalence of the mutation pressure argument was influenced by a historical focus on selective changes in allele frequencies within a gene pool of standing variation (see [38]). This gene-pool regime is certainly an extremely important evolutionary scenario, especially for highly polygenic quantitative traits in populations of sexual organisms, the same regime accessed by breeders who artificially select. Within this picture, the main source of variation that natural selection can act on comes from recombination within the gene pool [42]. The primary role of mutation is then to act as a fuel that replenishes the gene pool with new variability; it is not thought to otherwise impart any directionality.

The emphasis on the gene-pool picture can also help explain the historic neglect of models that explicitly treat the introduction of new mutations. As reviewed by McCandlish and Stoltzfus [43], models that include both the introduction of a new allele by mutation and its fixation or loss only became important four decades after the opposing pressures argument first appeared with the pioneering 1969 papers [44, 45] on neutral evolution, and then only for the case of neutral mutations. It took a further 30+ years before Yampolsky and Stoltzfus [22] applied these origin-fixation models in an adaptive context and showed that fixation rates can strongly depend on the rate at which novel mutants are introduced into a population without needing to be in a regime with high mutation. These predictions, valid for a regime beyond the “gene pool” scenario, have been corroborated and extended with more detailed population genetic calculations [8, 38, 46–48].

A series of important recent experiments and retrospective analyses in molecular evolution have demonstrated that mutational biases^2^ have significant effects on adaptive evolutionary outcomes. Examples include single celled organisms such as *Escherichia coli* [49], *Mycobacterium tuberculosis* [48, 50], *Pseudomonas fluorescens* [51] and *Saccharomyces cerevisiae* [48]. In particular, good agreement was found with theoretical calculations of the effects of the bias for such organisms, which can range over as much as two orders of magnitude, although much smaller biases can be picked out in the data [48, 52]. A meta-analysis of parallelisms found clear evidence for transition bias across taxa ranging from insects to mammals [53]. A related analysis of mutations related to altitude adaptation in the haemoglobin of high-flying birds found that CpG mutations were highly over-represented, as expected from mutation bias [54]. While some controversy over the full implications of this evidence remains [37, 55, 56], there is little doubt that mutational biases can significantly affect adaptive outcomes in predictable ways [52].

In summary then, the old “mutation pressure” argument should be retired from employment in opposing biases in the introduction of variation affecting adaptive evolutionary outcomes. Theoretical calculations, detailed experiments, and retrospective analyses of mutational biases have robustly shown that this argument does not hold in key regimes where the introduction of new mutations plays an important role.

#### S9.3 Historical theme 3: Ultimate and proximate causes

Another principal axis around which these disagreements revolve, and the one we think is the most interesting for this paper, concerns views on evolutionary causation. These are often framed through the lens of Ernst Mayr’s famous distinction between ultimate and proximate causes [57]. The former relate to “why is it like this” questions asked by evolutionary biology whereas the latter encompass the “how something works” questions of functional biology. This distinction has been influential in shaping the landscape of the discussion, not least because development is often a priori labelled as a proximate cause. See for example Mayr’s 1994 response to reading the literature on evo-devo [58]: “ *.. the two kinds of causations were hopelessly mixed up. All I can say is that I hope that the day will come when people realize that the decoding of a genetic program is something very different from the [adaptive] making and changing of genetic programs that is done in evolution.* ". In other words, Mayr’s argues here that developmental bias is a proximate cause because it ultimately arises from an adaptive historical process behind the developmental program. Mayr’s views were likely more complex [59], but that has not stopped some authors from simply equating ultimate causes with adaptive ones [60]. Needless to say, the reception of the ultimate-proximate distinction has a tangled intellectual history [33, 59, 61–67], with much disagreement about the definition and the usefulness of the distinction.

One influential way to explore the possibilities for ultimate and proximate causation is to frame them in terms of Gould’s nomenclature of isotropic variation [68]: “*Under these provisos [variation is small, copious, and undirected and evolution proceeds in gradual steps], variation becomes raw material only – an isotropic sphere of potential about the modal form of a species . . . [only] natural selection . . . can manufacture substantial, directional change.*”. Scholl and Pigliucci use this concept to explain the differences between those who do and those who don’t believe that developmental bias can be an ultimate evolutionary cause [64].

If variation is isotropic, then development is not an evolutionary cause. But if variation is anisotropic and biased, development could be a causal factor in evolutionary outcomes if natural selection cannot overcome the biases. While this way of formulating the questions has the appeal of clarity, it suffers from the fact that many adaptationists agree that variation can be anisotropic, but may still not consider it to be an ultimate cause. For example, Dawkins criticizes the assumption of isotropic variation as a “caricature of a Darwinian” in Chapter 11 of *The Blind Watchmaker* [1]. Nevertheless, isotropic variation is often an unspoken assumption in arguments around evolutionary causes [64]. It can also function as a “*useful fiction*" to help clarify arguments.

Because ultimate and adaptive causes are so closely entwined in the literature, we will take a small detour to explore different types of adaptationism. Consider, for example, how Dawkins describes his view of the role of processes such as neutral mutations or developmental bias in chapter 11 of *The Blind Watchmaker* [1]: “*Of course, large quantities of evolutionary change may be non-adaptive, in which case these alternative theories [neutralism, mutationism] may well be important in parts of evolution, but only in the boring parts of evolution . . .* ". To put this statement into a broader context, it is helpful to use Godfrey-Smith’s[69] influential threefold categorization of adaptationism ^3^. At one end of the spectrum, there is the strongest category, *Empirical Adaptationism*, where natural selection is the dominant explanatory mechanism behind evolutionary outcomes. At the other end of the spectrum, *Methodological Adaptationism* makes the much weaker assumption that evolutionary research should focus on adaptation because non-adaptive processes are harder to study. The third category is *Explanatory Adaptationism*, where selection has unique explanatory importance because, on this view, the big and interesting questions of evolutionary biology revolve around the origins of adaptations. Thus, selection will be “*of central important even if it is rare*” [69]. Godfrey-Smith argues that Dawkins is an explanatory adaptationist for whom even strong anisotropies in the introduction of variation that affect evolutionary outcomes are “*boring* " because they are random w.r.t. improvement of the organism and therefore don’t explain the adaptations he is primarily interested in.

These different attitudes to adaptationism will have implications for the interpretation of new evidence related to the role of developmental bias. For empirical and explanatory adaptationists, ultimate explanations will by definition be selective ones. For the methodological adaptationist, the question is open but could be (too) hard to adjudicate. While an empirical adaptationist may be willing to include developmental bias as a significant evolutionary cause if sufficiently strong evidence demonstrated its influence on evolutionary outcomes, such evidence is unlikely to be convincing for explanatory adaptationists because theirs is fundamentally a philosophical position on what is important or interesting.

To avoid the complexities surrounding adaptationism, we turn to an important definition of ultimate causes put forward by Ariew [71], who equates them with “statistical population-level explanation[s]” that answer questions about the prevalence and maintenance of traits in a population. Such ensemble-level causes of course include natural selection, but could also include genetic drift and other causes that act on populations at a population-level. We will then add to Ariew’s definition the additional stipulation that such causes be universal, that is they are not limited only to certain taxa. This more stringent requirement has the advantage of clarity. If it can be shown that certain developmental biases are universal and also affect long-term evolutionary outcomes at a population level, then it is hard to see why these should not be classed as ultimate rather than proximate causes.

#### S9.4 Historical theme 4: Contingency, convergence, and counterfactuals

The final set of implicit assumptions that bedevil this debate relates to the role of contingency in evolution. On the one hand, we have Gould’s memorable expression of radical contingency in his book *Wonderful Life*: [72]: “*..any replay of the tape [of life] would lead evolution down a pathway radically different from the road actually taken*”. This famous gedankenexperiment of rerunning the tape of life is effectively an exercise in counterfactuals – things that could have happened but did not – which play an important role in these kinds of big-picture arguments about evolution. In this case, Gould was writing about the development, more specifically, the emergence of body plans (baupläne or blueprints) during the Cambrian explosion. Once established, these developmental patterns are largely fixed and strongly constrain the spectrum of future evolutionary innovations. Gould argued that contingent accidents of history determine which bodyplans survived, and which did not. But the arguments he made about contingency were broader than just this example, and multifaceted, as emphasized by Beatty [73]. One can ask what would happen if one reran the tape of life from a time well before the Cambrian explosion. If similar conditions to the Cambrian explosion occurred, would the same initial set of bodyplans emerge? While we are not sure exactly what Gould’s view was on this specific question, given his take on contingency, it is likely that he would have thought that the bodyplans that first appear on a replay would be different, and therefore a subset of a much larger possibility space of body plans that could have appeared but didn’t.

The unimaginably vast size of genotype spaces also plays a role in arguments for contingency (see [74] for a broader discussion). Nature can only have explored a minuscule fraction of the space of all genotypes, and so it is natural to assume that if life were to restart in a different part of that genotypic space, we would observe different evolutionary outcomes.

There range of views on contingency is broad. On the other side of this debate, we have important thinkers such as Simon Conway Morris, one of the scientific heroes of Gould’s book [72] who, rather ironically, has a radically different interpretation of the research that Gould was popularizing. He argues that something close to replaying the tape of life is “*ubiquitous*" as evidenced by many striking evolutionary convergences, where similar phenotypes evolve independently [75]. In other words, his view on counterfactuals is that we should expect something similar to evolve again when we replay the tape of life. Presumably, he believes that the possibility space of biologically feasible body plans is much more constrained (by what he calls “*deeper structures*") than someone emphasizing contingency would think. Dawkins’s concept of the evolution of evolvability, and Mayr’s arguments about the ultimate origin of developmental programs that we quoted above, don’t necessarily assume that the possibility space of body plans is small. But they do assume that natural selection has access to alternative body plans. Behind the Gouldian perspective on contingency, there may also be an assumption that it is very hard for alternate body plans to appear as potential selectable variation, once a given set has fixed. Such a pre-commitment will affect how useful one thinks the concept of ultimate (adaptive) evolutionary causes are.

Where someone sits on this contingency to convergence continuum, and what specific background assumptions they hold about the space of phenotypic possibilities and the possibilities of changing developmental programs, can influence how arguments about the relative role of developmental bias and natural selection in determining evolutionary outcomes are weighed [74]. If one believes that developmental programs (e.g. those of elephants or octopuses) are contingent accidents of history, that many different options could have occurred, but that once fixed they are hard to change, then it is easier to believe that developmental bias is akin to an ultimate cause of evolutionary outcomes, or perhaps that Mayr’s distinction itself is not that useful. On the other hand, if one thinks that developmental programs are themselves quite amenable to natural selection, then developmental bias is less likely to be interpreted as an ultimate cause. This latter viewpoint also makes the implicit assumption that the timescales on which natural selection can access and traverse the landscape of developmental programs are accessible in practice.

## Footnotes

1 We note that the metaphor of Occam’s razor in a GP map was used, to our knowledge, for the first time for the particular example of L-systems by Lehre and Haddow [10]

2 To avoid misunderstandings, this relatively small number of sequences is what is needed to find all the level-5 coarse-grained structural classes found in the databases. And while the structure is important for facilitating function, it is only part of what needs to be specified. For example, at the finer level of individual secondary structures (inferred) frequencies of individual secondary structure frequencies found in the database varied over 15 orders of magnitude for this length [14] (with a distribution that closely follows that of sampled frequencies *fp*). At an even finer level, a catalytic ribozyme, for example, needs not only a specific structure but also a particular sequence in its active site. Thus, the probability of finding a ribozyme in random samples is smaller than the number needed to just find its secondary structure. The main point from [9] is that this number is many orders of magnitude smaller than one might naively expect based on the full size of the sequence space

3 We only consider one half of the phenotype since all biomorphs are axially symmetric. We use default parameters in the block

4 For example, the simplifying assumptions in the analytic model mean that each genotype in a given phenotype’s neutral set has the same value of *g*_9_. In the computational data, this is not fully true: any genotype with a zero for all positions that affect x-components is a vertical line, regardless of the value of *g*_9_ and this genotypic diversity in the neutral set of the ‘line’ phenotype may facilitate additional phenotypic changes.

5 The median number of transitions is 16 in the computational results, which coincides with the analytic value for the phenotypes with *Np* = 2, which dominate the data due to their high number.

6 The criterion for the strong-selection weak-mutation regime is that the product of mutation rate and population size is small and the product of the population size and selective advantages large [75]: here these quantities are 9 *× µN* = 0.45 (where the factor of nine accounts for the fact that the mutation rate is per-site) and *N × s ≥* 10.

7 Inspired by a biomorph shape in ref [77].

8 Strictly speaking, not all genotypes in a neutral set are connected by neutral mutations and Wagner’s [36] phenotype evolvability needs to be computed for each mutationally connected subset, i.e. each neutral component [76].

9 Note that since phenotype evolvability values depend sensitively on the coarse-graining, the exact nature of possible single-step phenotypic transitions and thus the shortest phenotypic paths also depend on the coarse-graining. However, our argument holds as long as phenotypic evolvabilities are higher than genotypic evolvabilities, since this ensures that exploring neutral spaces enables a higher number of transitions than are possible from a single genotype. This condition is met even in the fine-grained analytic model (see Fig 4).

10 One might worry that the correlation with phenotypic frequencies is a spandrel, a byproduct of some other adaptive process. For example, higher phenotypic frequency correlates with increased stability [106] and phenotype robustness. However, the effect of stability is unlikely to explain the orders of magnitude range of observed frequencies, see e.g. [12] for a discussion, let alone the tight correlations observed in [9, 14, 107]. Adaptive evolution for mutational robustness may also occur, but its effect is likely much smaller than the effect that phenotype bias has on mutational robustness [3, 14].

1 In the context of previous models based on constrained/unconstrained approaches, this is equivalent to having multiple neutral components (NCs) per neutral set, for example in ref [6]: in that case, the constrained-unconstrained calculations give the size of a single connected NC, but the final result has to be multiplied by the total number of NCs to obtain the neutral set size.

2 For example, on average, individual transitions, nucleotide changes within a class (purines or pyrimidines) are significantly more likely to occur than individual transversions (nucleotide changes between classes).

3 The distinctions can be refined further, e.g. Lewens has seven categories [70].

